# Invasive alien mammals of European Union concern

**DOI:** 10.1101/2021.04.21.440832

**Authors:** Lisa Tedeschi, Dino Biancolini, César Capinha, Carlo Rondinini, Franz Essl

**Affiliations:** Global Mammal Assessment programme, Department of Biology and Biotechnologies, Sapienza University of Rome, Viale dell’Università 32, 00185 Rome, Italy; Centro de Estudos Geográficos, Instituto de Geografia e Ordenamento do Território – IGOT, Universidade de Lisboa, Lisboa, Portugal; BioInvasions, Global Change, Macroecology-Group, Department of Botany and Biodiversity Research, University of Vienna, Rennweg 14, 1030 Vienna, Austria

**Keywords:** biological invasions, environmental impact, pathways of introduction, spread, zoonotic diseases

## Abstract

Biological invasions have emerged as one of the main drivers of biodiversity change and decline, and numbers of alien species are rapidly rising. The European Union established a dedicated regulation to limit the impacts of invasive alien species (IAS), which is focused on a Union List of IAS of particular concern. However, no previous study has specifically addressed the ecology of invasive alien mammals included in the Union List.
We performed a systematic review of published literature on these species. We retrieved 262 studies dealing with 16 species, and we complemented these with the most up-to-date information extracted from global databases on IAS.
We show that most of the study species reached Europe as pets that escaped from captivity or were intentionally released. On average, 1.2 species’ new first records/year were documented in European countries in the period 1981-2020, and most species are still expanding their alien ranges colonising neighbouring territories. France, Germany, Italy, and The Netherlands are the most invaded nations, and the muskrat (*Ondatra zibethicus*), the raccoon dog (*Nyctereutes procyonoides*), and the American mink (*Neovison vison*) are the most widespread species, having invaded at least 27 countries each. Invasive mammals of European Union concern are threatening native biodiversity and human well-being: worryingly, 81.3% of the study species are implicated in the epidemiological cycle of zoonotic pathogens.
Containing the secondary spread to further countries is of paramount importance to avoid the establishment of new populations of invasive mammals and the related impacts on native communities, ecosystem services, and human health.
Our results offer the most updated compendium on the ecology of invasive mammals of European Union concern, that can be used to assist environmental policies, identify and subsequently fill knowledge gaps, and inform stakeholders.

**GRAPHICAL ABSTRACT:** 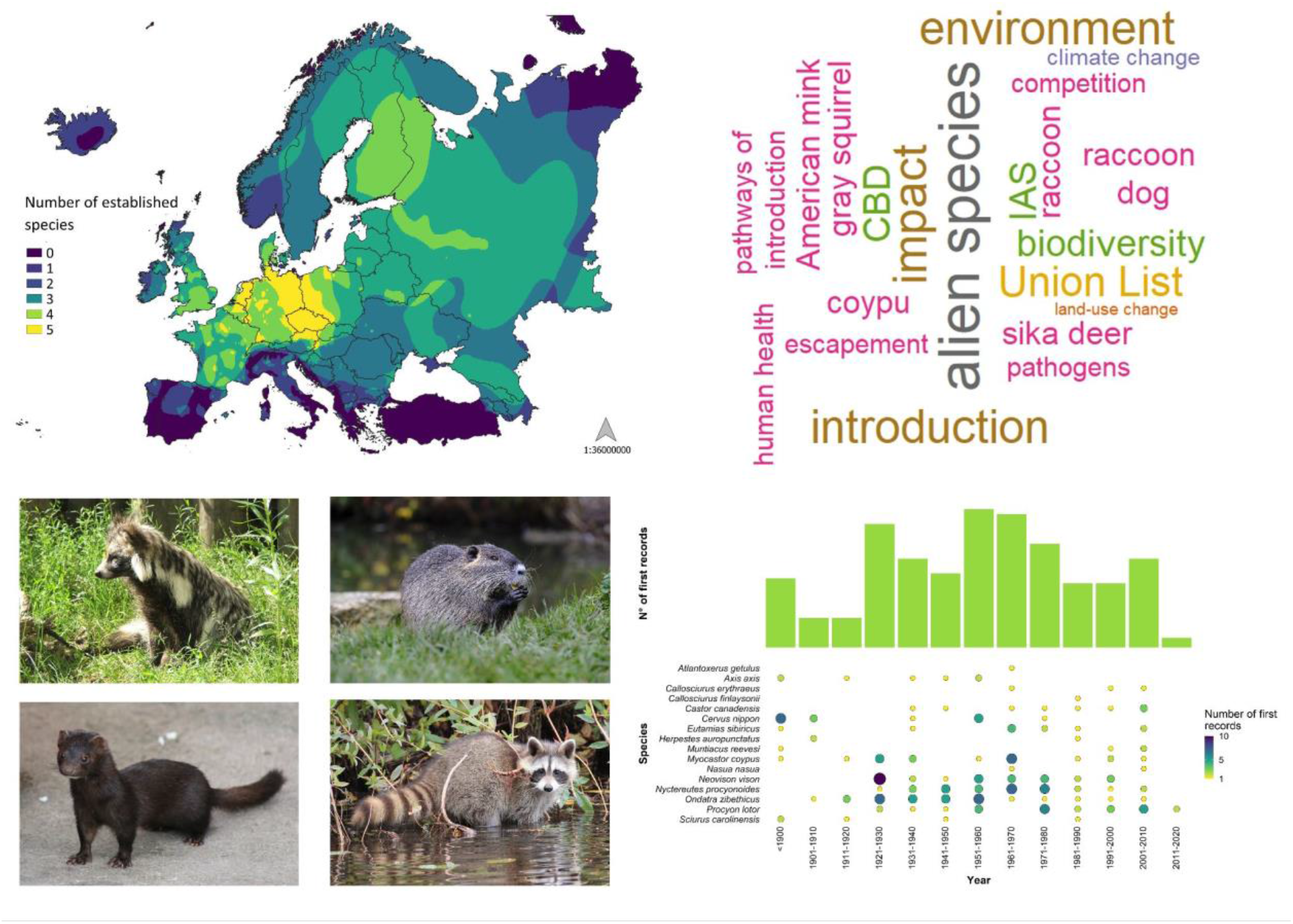

Graphical abstract: Invasive alien mammals of European Union concern.

The figure illustrates how the introduction of a species in few new areas, followed by a lag phase of adaptation and sometimes enriched by further subsequent releases, can rapidly lead to the colonisation of large parts of a continent. On the top left, a heat map with species’ richness in countries of Europe. On the top right, a word cloud with the main keywords of our literature search and some of the study species’ names. On the bottom left, four out of 16 study species: in clockwise order, the raccoon dog (*Nyctereutes procyonoides*), the muskrat (*Ondatra zibethicus*), the American mink (*Neovison vison*), and the raccoon (*Procyon lotor*). On the bottom right, the temporal distribution of the first records of the study species in the countries of Europe.

## INTRODUCTION

The human-mediated introduction of species to regions outside their native range has become one of the main drivers of biodiversity change and decline in recent human history (IPBES 2019). Despite a rise in awareness, and the adoption of legislation to reduce these introductions, the number of newly-introduced species has risen strongly in recent decades (Seebens et al. 2017) and is expected to continue to do so in the future (Seebens et al. 2020). International trade, global transportation networks (Hulme 2009), land-use change (Essl et al. 2020b), and climate change (Diez et al. 2012, Bellard et al. 2018) are the main drivers promoting species introduction and spread, and they continue to intensify. Many species introduced in new regions fail to establish self-sustaining populations or remain localised, whereas others become permanent additions to the receiving ecosystems and spread over substantial distances. In doing so, they can cause severe impacts on native biota (Blackburn et al. 2019) at different biological organisation levels (Hawkins et al. 2015), ecosystems services (Vilà & Hulme 2017), and human livelihoods (Bradshaw et al. 2016), i.e., they become invasive alien species (IAS).

The prevention and mitigation of biological invasions in Europe is a significant challenge, as policies are devoted to the free circulation of goods and people (Genovesi et al. 2015). To address this issue, the European Union (EU) adopted the Regulation (EU) No 1143/2014, aimed at the prevention of IAS introduction and spread (EU 2014). The Regulation, informed by years of invasion science research (Genovesi et al. 2015), called for the creation of a list of IAS of Union concern, the Union List. Each member state of the EU is required to collect information and take actions on introduction, detection, and eradication of these species and to mitigate their impact (EU 2014). Furthermore, this subset of IAS is subject to a ban on intentional importation and trade in the EU.

Out of the 66 species currently included in the Union List, 11 (~ 17%) are mammals, highlighting the perceived impact of this taxon across Europe. Indeed, mammals represent 60% of the worst invasive terrestrial vertebrates in Europe (DAISIE 2009, Polaina et al. 2020) and, overall, more than 50 species of mammals are currently established in this continent (Biancolini et al. 2021). Furthermore, seven species with high invasive potential are in the initial phase of the invasion (i.e., restricted to the initial location of introductions and without established populations in Europe): the four-toed hedgehog (*Atelerix albiventris*), the American bison (*Bison bison*), the leopard cat (*Prionailurus bengalensis*), the Northern palm squirrel (*Funambulus pennantii*), the striped skunk (*Mephitis mephitis*), the sugar glider (*Petaurus breviceps*), and the Caucasian squirrel (*Sciurus anomalus*). Alarmingly, due to climate change, suitable climatic space is projected to increase for most invasive mammals in Europe (Polaina et al. 2020). For instance, this is the case for the coypu (*Myocastor coypus*; Schertler et al. 2020), the raccoon (*Procyon lotor*; Louppe et al. 2019), and the small Indian mongoose (*Herpestes auropunctatus*; Louppe et al. 2020). Invasive mammals exert negative impacts on biodiversity through competition (Mazzamuto et al. 2017), disease transmission (Collins et al. 2014), habitat alteration (Nogales et al. 2014), hybridisation (McFarlane et al. 2020), and predation (Dahl & Åhlén 2019).

Here, we provide a comprehensive synthesis of the invasion process, current distribution, and impacts of the invasive mammals of Union concern, by reviewing the literature for these species. Specifically, we (1) analyse trends in the published literature regarding 16 mammal species of Union concern (and candidate species to be included in the Union List) in the last 15 years (2005-2020), (2) summarise pathways of introductions, (3) reconstruct the temporal trajectories of mammal invasions, (4) illustrate geographic distribution patterns, (5) investigate environmental and (6) socio-economic impacts, with a focus on human health. This review updates the current knowledge on a subset of highly impacting mammals, that is crucial especially in the light of (and to accomplish the) recent developments of international agreements to protect native biodiversity (EU 2014, CBD 2020), and inform a wide audience of stakeholders and practitioners.

## METHODS

We searched for relevant publications on invasive mammals of Union concern. To provide a wider geographic context, the study area was not limited to the EU, but we considered the 47 member states of the Council of Europe, including also the outermost regions of the EU located in the North Atlantic (i.e., Azores, Madeira, and Canary Islands), but excluding the remaining ones (e.g., French Guiana, Guadeloupe). We selected 10 mammal species of the 11 included in the Union List (thus excluding the fox squirrel *Sciurus niger*, as no established populations are currently present in the study region); further, based on the list from Carboneras et al. (2018), we included another six species recommended for future inclusion in this list, excluding the species currently absent from Europe and species in the initial phase of the invasion (not established). Therefore, a total of 16 species were included in this review (Table 1).

**Table 1.**
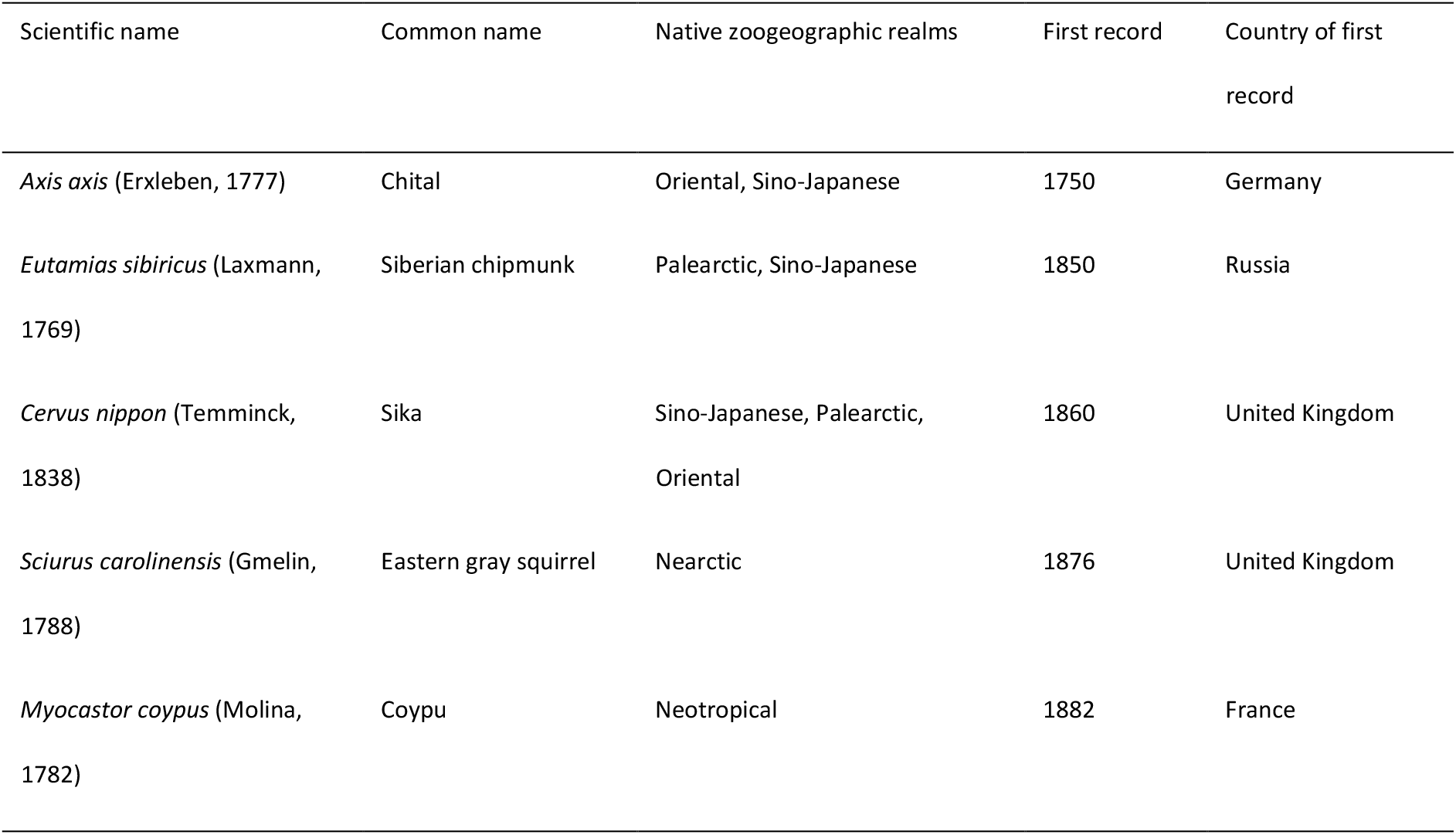
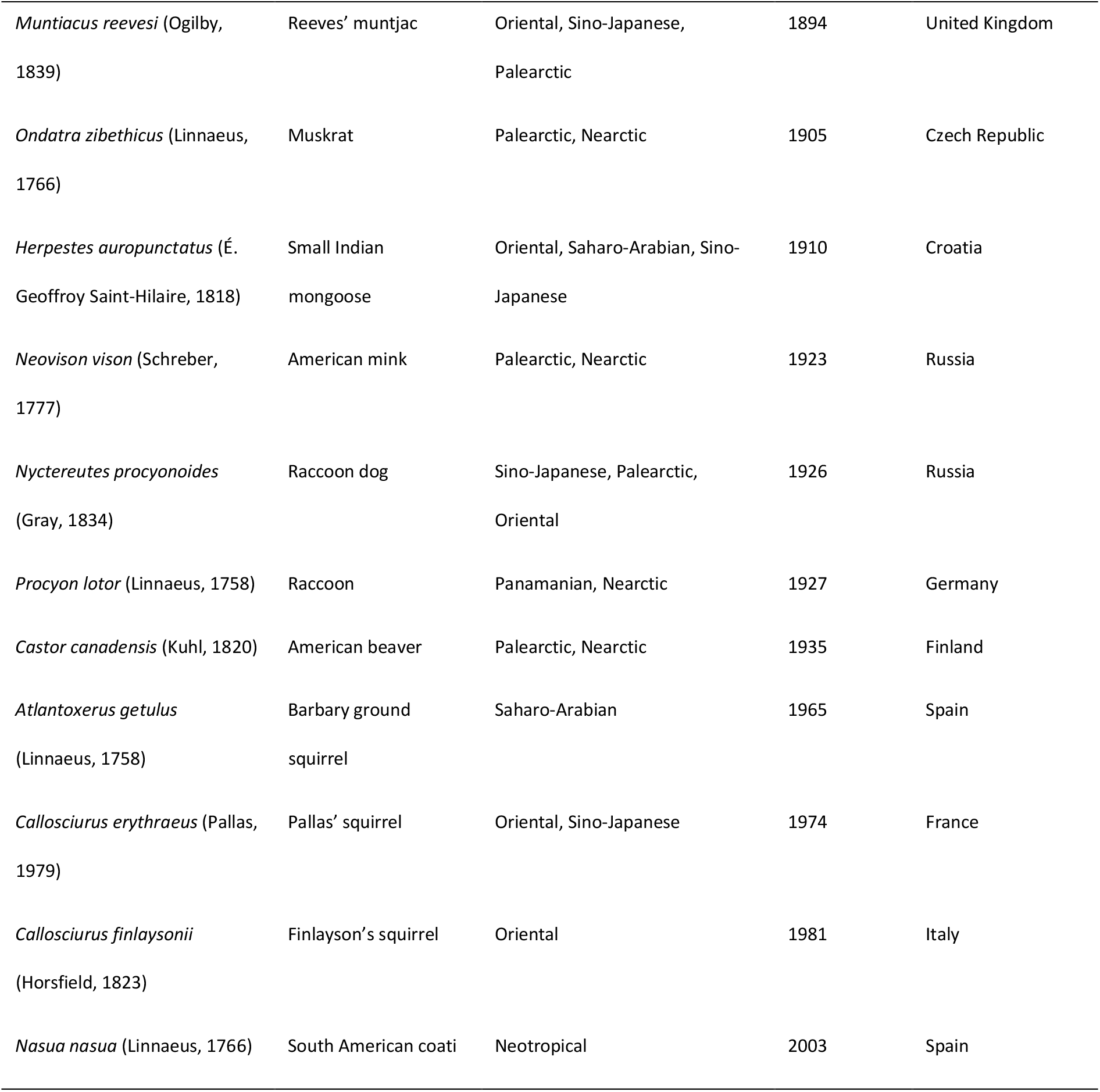
The 16 species included in this review. Scientific name, common name, native zoogeographic realms (following Holt et al. 2013), year of first record in Europe, and country of first record in Europe are indicated. Native zoogeographic realms are given for each species in decreasing order, based on the percentage of native range located in each realm.

### Literature search

The literature search was carried out by the first author following the Preferred Reporting Items for Systematic Reviews and Meta-Analyses (PRISMA) methodology (Appendix S1; Moher et al. 2009) in August and September 2020. For each species, we downloaded available information from the EU Commission CIRCA website (https://circabc.europa.eu/ui/welcome) in the form of the EU Non-Native Risk Assessment Scheme or the Great Britain Non-Native Risk Assessment Scheme. In addition, we downloaded CABI species datasheets (www.cabi.org) and the NOBANIS factsheets (www.nobanis.org). Hereafter, for brevity, we will refer to all these documents as “datasheets”. As these datasheets were highly comprehensive on the scientific knowledge of the study species at the time of completion, the time range of the search for additional articles was adapted for each species, depending on the date of the most recent datasheet. If no prior datasheet was found, the search in the literature databases was performed without a temporal filter.

Subsequently, we searched for additional recent information on the species in Scopus and Web of Science (WoS). On Scopus, we conducted an advanced search refined for the sub-areas of Agricultural and Biological Sciences and Environmental Sciences. On WoS, we performed a basic search without sub-areas limitations, except for the American beaver, for which we filtered WoS results due to the large literature retrieved on unrelated topics (such as engineering or fluid mechanics). For each species we conducted a separate search with a combination of the scientific name and its synonyms, common name(s) and the relevant keywords, linked by the Boolean operators AND/OR. The list of countries encompassing the alien range, to be used as species-specific keywords, was obtained from the global Distribution of Alien Mammals database (DAMA; Biancolini et al. 2021). Keywords were identified *a priori* based on the known alien distribution of each species, European Regulation, invasion history, characteristics linked to invasiveness and caused impacts (Appendix S1).

Two species are identified with different scientific names in the Union List and the IUCN Red List, namely the small Indian mongoose (*Herpestes javanicus* changed in *Herpestes auropunctatus*) and the Siberian chipmunk (*Tamias sibiricus* changed to *Eutamias sibiricus*). We are aware of the recent taxonomic revision and, in this work, we choose to follow the IUCN taxonomy (IUCN 2020).

### Data extraction and preparation

To be included in the review, literature results had to fulfil the following criteria: refer to the European territory (defined as described above), be written in English and contain information related to at least one of the following: (i) year(s) of first record of a study species, (ii) point(s) of first record, (iii) pathway(s) of introduction, and/or (iv) impact(s).

A primary research topic was assigned to each publication, based on its aims, as follows: community ecology, datasheet (sub-topics: CABI, NOBANIS), economic impacts, environmental impacts (sub-topics: competition, disease transmission, habitat alteration, hybridisation, native species replacement, predation), general ecology (sub-topics: activity pattern, behavioural responses, diet, ecological modelling, reproduction, space use), genetics (sub-topics: genotyping, methodology, phylogeny, population genetics), health status, management, population status, review, risk assessment (sub-topics: EU NNRA, GB NNRA, other), social impacts, and systematics. A distinction was made amongst species’ pathogens studies, classifying them based on the threat posed to native fauna (topic: environmental impacts/disease transmission), to humans (social impacts) or a general investigation of species’ pathogens (health status).

The same paper investigating two (or more) species was counted only once when illustrating trends in the literature and was referred to the first species in alphabetical order (scientific name) to be investigated. For instance, Bertolino and Lurz (2013) investigated both Pallas’ squirrel and Finlayson’s squirrel, but the paper was counted for Pallas’ squirrel. In such cases the relevant information (e.g., regarding pathways of introduction) was extracted for all the studied species.

In tables and figures, countries are indicated by their ISO country code. RU refers to the European part of the Russian Federation. Species with occasional occurrences (i.e., not established) or with an unknown status are indicated as “casual presences”. Alien ranges for the study species were obtained from DAMA (Biancolini et al. 2021), as well as the list of all established mammals in Europe, regardless of their inclusion in the Union List, to get a more comprehensive picture of alien mammals’ status in Europe. Native zoogeographic realms (Holt et al. 2013) for the study species and all established mammals in Europe were obtained based on species native ranges (IUCN 2020). Marginal parts of native ranges occurring in less than 1% of a zoogeographic realm were not considered.

Capellini et al. (2015) and Blackburn et al. (2017) identified body size, litter size, litters per year, and generation length as species’ traits favouring introduction, establishment, and spread of invasive mammals. We extracted these trait values from the recently developed Coalesced Mammal Database of Intrinsic and Extrinsic traits database (COMBINE; Soria et al. 2021). Reproductive life span was calculated as the difference between maximum longevity and age at first reproduction (Soria et al. 2021).

Each species was assigned to one or more pathway(s) of introduction following CBD categorization (CBD 2014, Biancolini et al. 2021). First records of the species were mainly obtained from the Version 2 (last updated in March 2021) of the Alien Species First Records Database (Seebens et al. 2017). For first records obtained from articles encountered during the literature review, the earliest year was retained in cases of multiple and/or continuous introduction in a country. Information regarding species’ pathogens (e.g., prevalence) was extracted both from original papers and from reviews encountered during the literature search.

## RESULTS

### Relevant literature and publication trends

The literature search yielded 3322 papers published between 2005 and 2020 that were subjected to screening by reading the title and abstract; if these elements did not provide definite information, the full text was screened. All the species but one (Barbary ground squirrel) had at least one datasheet available for download, for a total of 36 published datasheets. After this screening, 591 articles were retained, and their full text assessed for eligibility. A backward reference search (“snowballing”) was performed on the reference list of each of these articles to identify other relevant publications, adding further 30 studies. Duplicate records resulting from an overlap of the database outcomes were removed. 26 articles could not be assessed due to access’ restrictions or because they were not written in English. Eventually, 262 publications were included in the review (Appendix S3).

Published information was available mostly for the raccoon (that accounted for 15% of all publications), the American mink (14%), and the sika (12%) (Appendix S2). The majority of the datasheets collected (88.2%) were published from 2009 to 2014 (Appendix S2) and, due to the temporal filters adopted, for most of the study species the literature search supplied mainly papers published after 2015. Accounting for these filters adopted in the literature search, mainly species’ environmental impacts were investigated (24.1% of all papers), with a peak of publications in 2017-2018, followed by studies on health-related issues (17.6%) and social impacts (11.8%).

### Taxonomic characterization, traits, and native ranges

The 16 study species belong to three orders and nine families. Half of them belong to the order Rodentia (Appendix S2), whereas the remaining are either from Carnivora (31.3%) or Artiodactyla (18.7%). The Sciuridae family is the most represented, accounting for 31.3% of all species, followed by Cervidae (18.8%) and Procyonidae (12.5%). In comparison, the full ensemble of alien mammals established in Europe is divided into seven orders and 17 families. The order Artiodactyla is the most numerous (28.3%), followed by Rodentia (22.6%) and Carnivora (17%; Appendix S2). The most represented family is Cervidae (15.1%), followed by Leporidae (13.2%) and Bovidae and Mustelidae (11.3% each).

Adult body mass (in grams) for the study species varied between 85 for the Siberian chipmunk and 53000 for the sika (mean 9762.4; Appendix S2). Litter size was between 1 for sika and Reeves’ muntjac and 6.4 for the muskrat (mean 3.3). Litters per year ranged from 1 for American beaver, sika, American mink, and raccoon to 2.6 for the muskrat (mean 1.5). Lastly, generation length (in days) fluctuated between 2941 for the Barbary ground squirrel and 8504 for the sika (mean 5781).

The study species originate mainly from the Palearctic, Sino-Japanese, and Oriental zoogeographic realms (Appendix S2). Similarly, the full ensemble of alien established mammals in Europe originate mainly from Palearctic, Saharo-Arabian, and Sino-Japanese realms (Appendix S2).

### Pathways of introduction to Europe

The main pathway of introduction for the study species in Europe was the pet trade (Fig. 1). A total of 68.8% of the species escaped after they were introduced at least once through pet trade (i.e., private owners), 50% escaped from zoos (i.e., public exhibitions), and 37.5% escaped after they were introduced to be bred in fur farms. One species was released in nature for biological control (the mongoose in Croatia), and another one for conservation purposes (the American beaver in Finland). The chital, introduced in Croatia, was the only species with an unknown introduction pathway, although subsequent repeated introductions within Croatia were reported for hunting purposes (Šprem & Zachos 2020). No study species was reported to be introduced as contaminant, stowaway, or via a corridor.

**Fig. 1.**
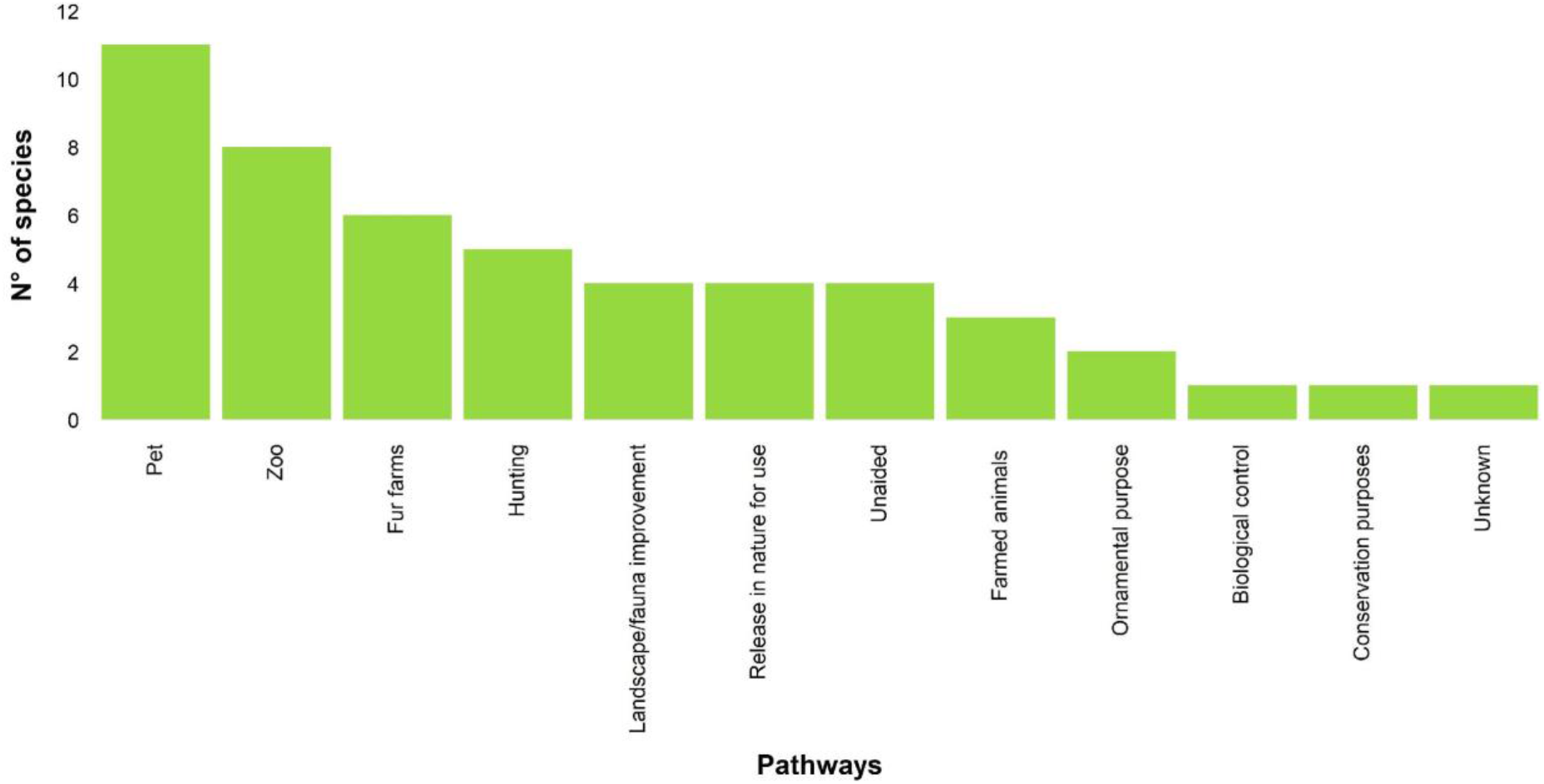
The CBD’s pathways of introduction of the study species in Europe (n = 50). Each species was assigned to one or more pathways. Pathways with zero occurrences and nomenclature not relevant for terrestrial mammals are not shown. Pathways names are abbreviated following CBD (2014).

### Temporal trajectories of mammal invasions in Europe

The rate of first records (of both established and casual presences) of the study species in countries of Europe increased on average from 1.4 new records/year over a 40-year period (1900-1940) to 2.3 records/year in 1941-1980, and then dropped to 1.2 records/year in 1981-2020 (Fig. 2, Appendix S4). Overall, the American mink, the raccoon dog, and the muskrat accounted together for 47.1% of first records.

**Fig. 2.**
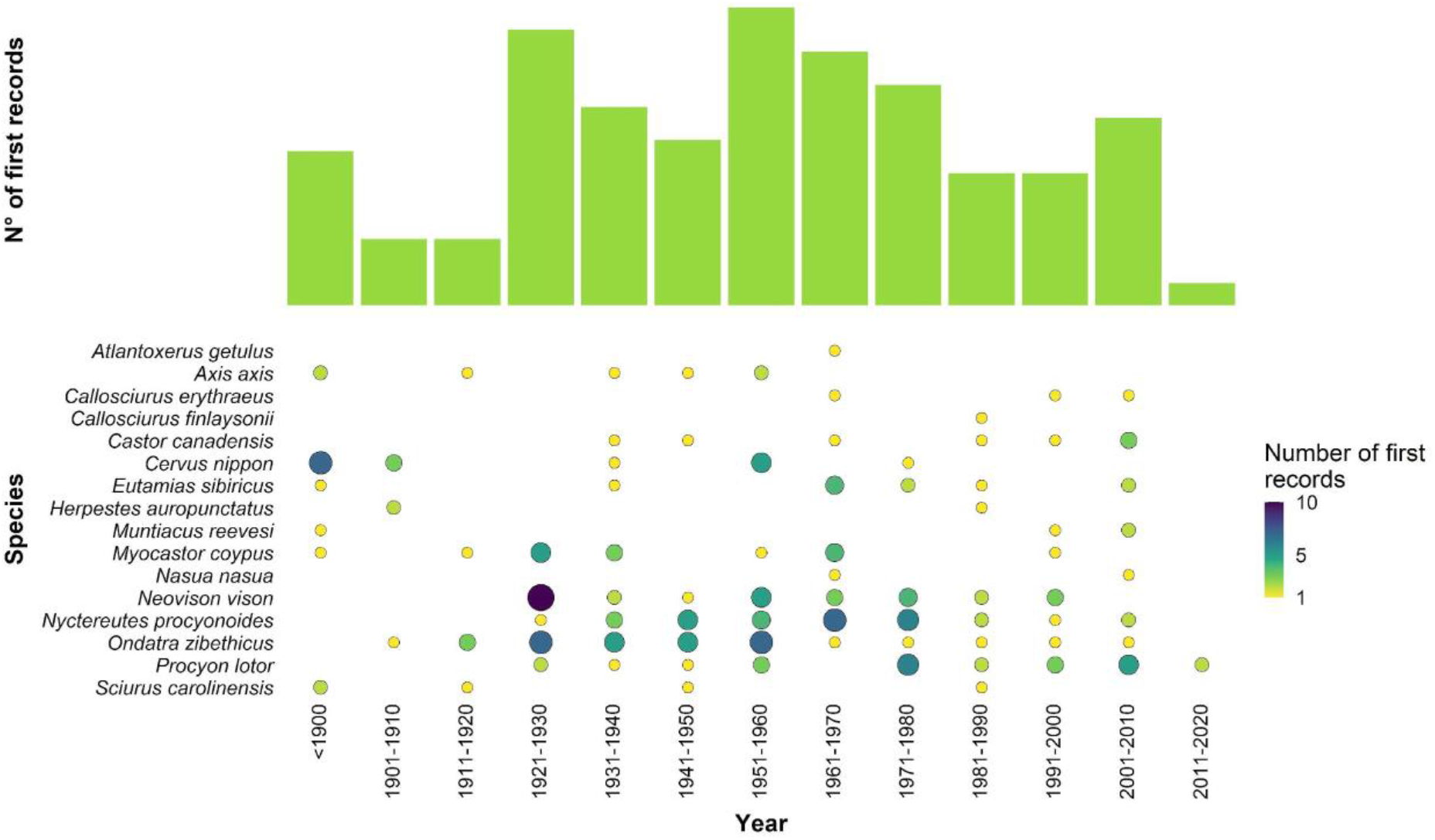
Temporal distribution of first records of the study species in the countries of Europe (n = 197). Point sizes represent the number of records per species and time period.

### Geographic distribution patterns in Europe

The United Kingdom and the Russian Federation first recorded three of the study species each, namely the sika (1860), the Eastern gray squirrel (1876), and the Reeves’ muntjac (1894) for the former, and the Siberian chipmunk (1850), the American mink (1923), and the raccoon dog (1926) for the latter (Appendix S4).

Considering the number of countries occupied, the most widespread species was the muskrat (established in 32 countries and with casual presences in three countries; Appendix S2), followed by the American mink (established in 28, casual in seven), the raccoon dog (established in 27, casual in seven), and the coypu (established in 24, casual in four). However, with respect to the area occupied only by the established species, the order slightly changes, with the raccoon dog becoming the most widespread, followed by the muskrat, the American mink, and the raccoon (Fig. 3).

**Fig. 3.**
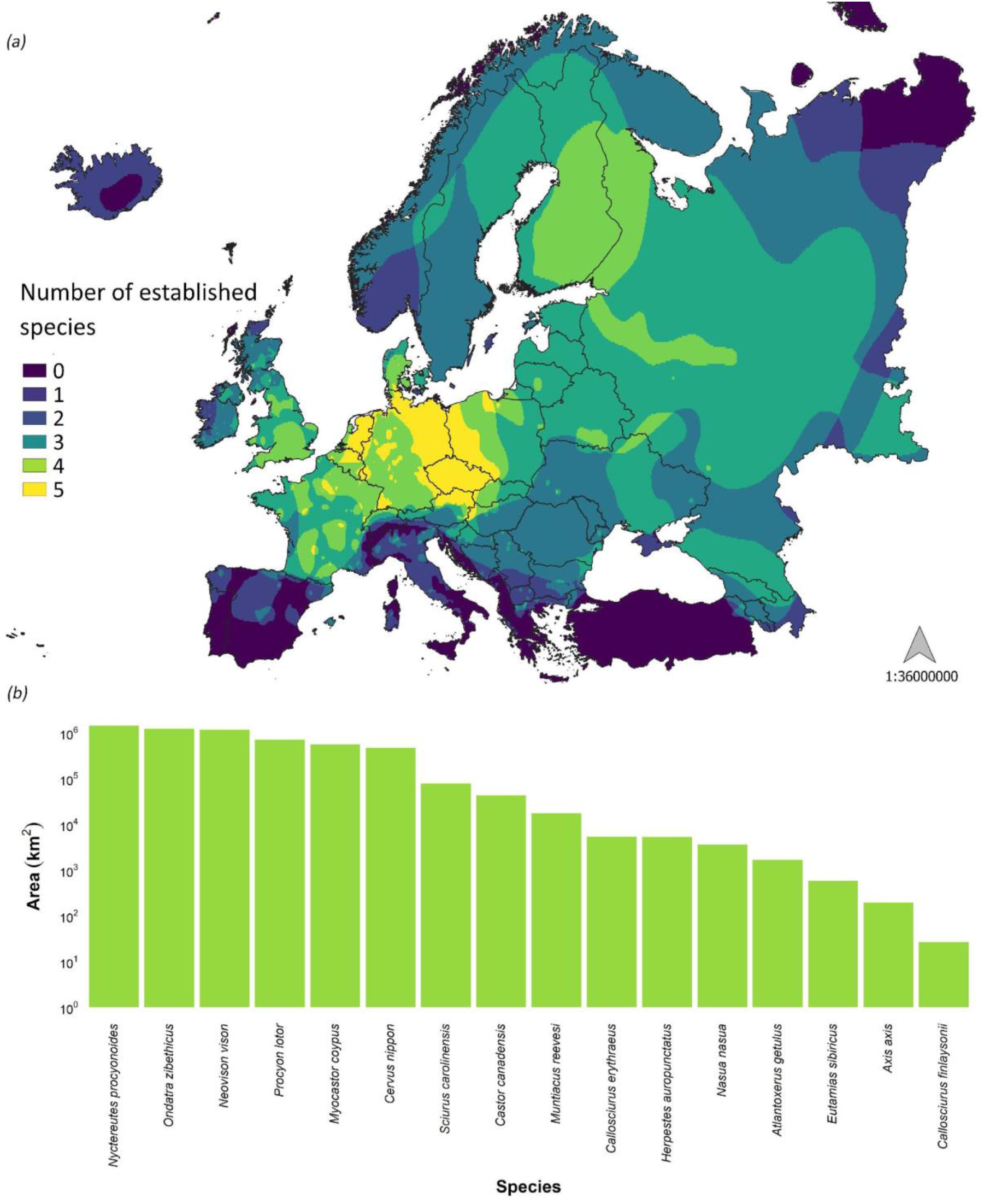
Established presences of the study species in Europe: (a) heat map showing study species’ richness in the study area; (b) area (log scale, km^2^) occupied by the study species.

Fig. 4 illustrates the invasion waves of the four species that invaded most of the European territory (the raccoon dog, the muskrat, the American mink, and the raccoon), including established presences and casual records.

**Fig. 4.**
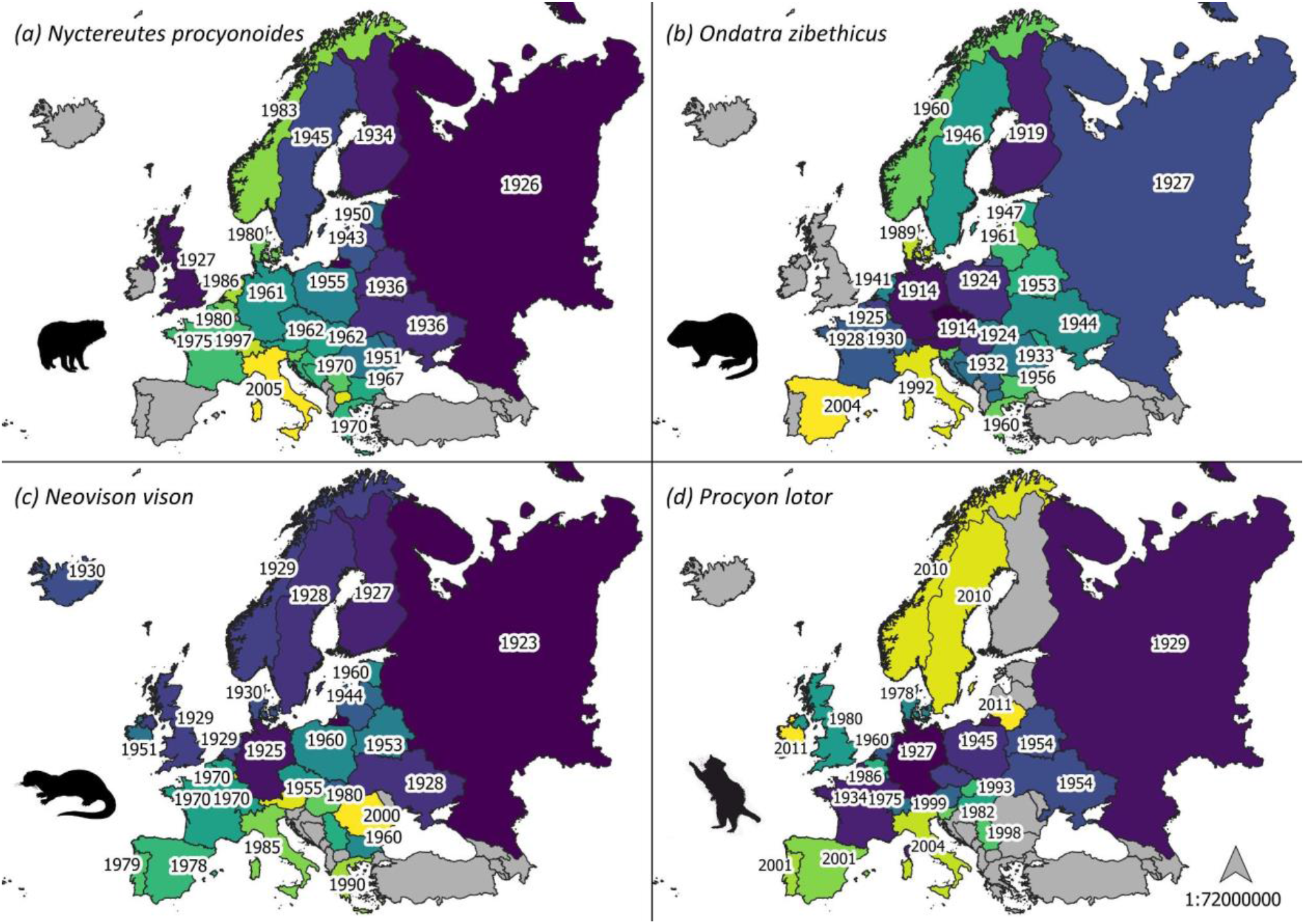
The spread trajectories of the four species that invaded most of the European territory: (a) the raccoon dog, (b) the muskrat, (c) the American mink, and (d) the raccoon. Countries are graded from the country invaded earliest (darker) to the latest (lighter). Year of the first record (when available) is shown. Countries without presences (established or casual) of the species are shown in grey.

### Environmental and socio-economic impacts in Europe

Among the studies retrieved during this search process, environmental impacts of invasive mammals have been broadly investigated, as shown by the number of papers published on this topic (n = 63; Appendix S2). Disease transmission was the most studied sub-topic (30.2% of the total number of papers related to environmental impacts), followed by predation (23.8%) and habitat alteration (14.3%). Studies on pathogens of invasive mammals (n = 19) revolved mainly around their helminthofauna (50.6% of the papers on pathogens, Appendix S2). Some of these pathogens were introduced in Europe with the study species, such as the nematode *Strongyloides callosciureus*, introduced with Pallas’ squirrel and potentially infecting the native red squirrel *Sciurus vulgaris* due to spill-back and spill-over processes (Mazzamuto et al. 2016) or the Squirrelpox virus, which can be often lethal for red squirrels and was introduced in UK and Ireland with gray squirrels (IUCN 2005, Invasive Species Ireland 2012).

The study species were found to be infected by 224 pathogens, of which 143 (63.8%) have zoonotic potential; 13 study species serve as potential reservoirs or are implicated in their epidemiological cycle (Fig. 5; Appendix S5). Specifically, regarding the most widespread study species, 48.6% of the pathogens known to infect the American mink have zoonotic potential; the percentage rises to 66.7% for the raccoon dog, 77.8% for the raccoon, and 100% for the muskrat (Fig. 5). Overall, studies on *Echinococcus multilocularis* (14 studies, three species), *Toxoplasma gondii* (nine studies, six species), and *Baylisascaris procyonis* (nine studies, one species) were particularly abundant among the study species (Appendix S5). Prevalence rates presented a high geographical and taxonomical variability: the prevalence of *E*. *multilocularis* ranged between 0% (in the raccoon and the raccoon dog in various countries; Kornyushin et al. 2011, Wahlström et al. 2012, EFSA 2015, Karamon et al. 2016, Oksanen et al. 2016, Duscher et al. 2017) and 28% (in the racoon dog in Slovakia; Oksanen et al. 2016); for *T*. *gondii*, it ranged from 0% (in mink in Spain and the raccoon in the Czech Republic; Criado-Fornelio et al. 2018, Kornacka et al. 2018) to 78.8% (in mink in Spain; Ribas et al. 2018); lastly, the prevalence of *B*. *procyonis* in raccoons ranged from 1.9% in Poland (Karamon et al. 2014) to 80% in Germany (Hohmann et al. 2002). Moreover, there are recent reports from Denmark and the Netherlands regarding SARS-CoV-2 infection in minks (Oreshkova et al. 2020).

**Fig. 5.**
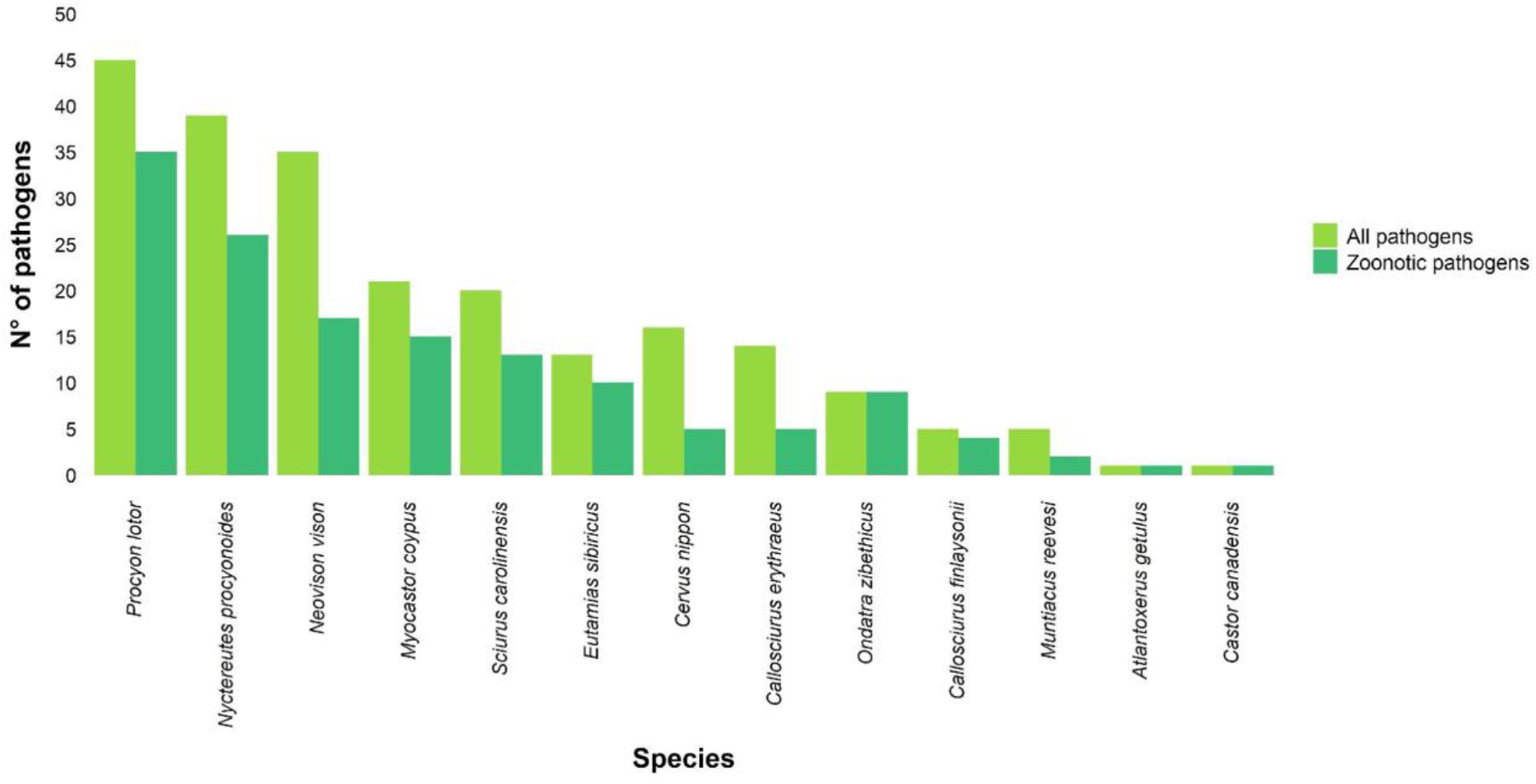
Total number of pathogens known to be infecting the study species (all pathogens) and pathogens with zoonotic potential (zoonotic pathogens). Species without recorded pathogen infections are not shown.

Regarding the second most investigated sub-topic (n = 15), the majority of studies analysed the predatory effects of American minks (40% of the total number of papers related to predation), raccoon dogs (26.7%), and Eastern gray squirrels (20%). Interestingly, 66.7% of American minks’ studies were performed in Poland, the whole of raccoon dogs’ studies in Scandinavia, and 66.7% of Eastern gray squirrels’ studies were conducted in the UK. Lastly, regarding habitat alteration (n = 9), 55.5% of the articles investigated the role of Barbary ground squirrels on native communities of the Canary Archipelago, in Spain.

## DISCUSSION

The majority of invasive mammals of Union concern reached Europe as pets that escaped from captivity, or were intentionally released. Although introductions of alien mammals have declined in Europe for more than 50 years, many study species are still expanding their alien ranges, colonising neighbouring countries. France, Germany, Italy, and The Netherlands are the most invaded countries, and the raccoon dog, the muskrat, the American mink, and the raccoon are the most widespread species. Invasive mammals of Union concern are threatening native biodiversity and human health with consequences largely overlooked in the past, such as new roles in epidemiological cycles of zoonotic pathogens (Oreshkova et al. 2020).

### Relevant literature and publication trends

Geographical and impact-related biases emerged from the reviewed literature. Charismatic, widespread, and detrimental species received more attention – in terms of publication numbers – than others, a trend already observed in invasion ecology (Pyšek et al. 2008). Apparently, documented environmental or socio-economic impacts and alien range’s size are also related to the number of publications. For instance, species having localized alien distributions, such as island invaders (the Barbary ground squirrel, the chital, the small Indian mongoose, and the South American coati) or urban dwellers (Finlayson’s squirrel) have been less investigated than more widespread species, such as the raccoon or the coypu. The well-acknowledged invasive potential of this species urgently calls for additional studies on their impacts and possible future spread. For example, the small Indian mongoose is a devastating island invader across the World, which could irremediably harm native biota in the Balkans mainland (Ćirović & Toholj 2016).

However, our results regarding publication trends shall be taken with caution, as our search did not include grey literature nor articles in languages other than English; this language barrier may generate biases and lead to knowledge gaps (Angulo et al. 2021).

### Taxonomic characterization, traits, and native ranges

Humans pose an initial “filter” to introduction (Clout & Russell 2008), selecting mammal species based on key traits (Blackburn et al. 2017), such as a large body mass, long reproductive lifespan, and large litter size. These last two key traits have been shown to also promote the subsequent phases of establishment and spread, along with frequent litters per year (Capellini et al. 2015). The mean adult body mass for our species was high – especially if compared with the mean adult body mass for mammals – but 75% of the study species not weighed much, as few of them (i.e., sika and chital) heavily skewed the mean. Regarding litter size, it is interesting to note that the most widespread species in Europe (both in terms of countries and area occupied) had also an above-average litter size, confirming the importance of this trait in the invasion stages consecutive to introduction (Capellini et al. 2015). As for the litters per year, the species above-average (more than 1.5 litter per year) were mainly rodents. Accordingly, litter size is larger in these socially monogamous species (West & Capellini 2016). Although a longer reproductive life span promotes introduction and establishment in mammals, apparently the study species with a higher value of this trait have rather localised distributions (the sika, the Eastern gray squirrel, the Reeves’ muntjac), possibly as an outcome of a low colonization pressure. On the contrary, widespread species (like the muskrat and the American mink) present a short reproductive life span. The discordance of some study species’ traits (adult body mass and reproductive life span) with what was found previously in the literature can be the result of the relative over-representation of mammals introduced in the past for goods and services (hunting, fur farming, transport; Blackburn et al. 2017), rather than a depiction of more recent introductions of species used as pets (such as squirrels) or for other aesthetic purposes.

With regards to the provenience of the study species, the Palearctic, Sino-Japanese, and Oriental realms were equally relevant. Previous studies (Genovesi et al. 2009, 2012) showed that the former and the Nearctic were the realms harbouring native ranges of more introduced mammals. Similarly as for species’ traits, this could be linked to the over-representation of species introduced in the past to be utilised by humans for other commercial purposes. Contrarily, the study species are mostly used as pets, and originate from eastern realms.

### Pathways of introduction to Europe

Overall, the study species were mainly kept in private or public collections or to be bred for fur, and subsequently escaped or have been released. We showed that pet trade was still a relevant pathway of introduction to Europe in the last 15 years: for instance, the Siberian chipmunk was first recorded in Ireland in 2007, probably released in nature by (or escaped from) private owners (Invasive Species Ireland 2019). Indeed, all the Sciuridae have been introduced at least once for companionship, enjoyment, recreation and/or trading. These species are charismatic and often released for “fauna improvement” of urban parks (as in the case of Siberian chipmunks in Italy; Mori et al. 2018).

Higher rates of establishment and spread are related to multiple releases and, in general, to a higher introduction effort (Clout & Russell 2008, Capellini et al. 2015). However, in the absence of accurate introduction records it is often challenging to distinguish between the natural spread of a species from invasion foci in adjacent countries and a deliberate release (for instance, from private owners) or an escape, especially for highly vagile species such as ungulates and carnivores. For example, recent genetic analyses have shown that new Eastern gray squirrel populations in Italy (supposedly originated by natural dispersal of individuals) derived in fact from other populations established almost 200 km far away (Signorile et al. 2016). Therefore, in the absence of clear evidence of unaided dispersal, it is inappropriate to assign this pathway of introduction to some species (Pergl et al. 2020).

### Temporal trajectories of mammal invasions in Europe

Despite the continuous alien range expansion throughout Europe, first records of alien mammals declined from the 1960s onwards (Fig. 2). This pattern has already been recorded at a global level for this taxon, and it is likely influenced by the most recent first records (Seebens et al. 2017). For instance, there were almost no first records of the study species in the last 10 years. However, longer monitoring is needed to assess the reliability of these trends (Seebens et al. 2017), especially to clarify if a saturation has been finally reached or if these patterns depend on other factors. As a matter of fact, the rapid decline in new introduction events can be attributed to the synergistic effects of increased awareness and stricter regulations on alien mammals bred for fur, exploited as game species, or used as pets across Europe (Seebens et al. 2017), especially since the implementation of the EU IAS Regulation (EU 2014).

First records in Europe were not evenly distributed among countries, as the UK and the Russian Federation first recorded three study species each. Interestingly, two of the most common species (the American mink and the raccoon dog) were first recorded in the Russian Federation, where they were introduced for fur farming. This comes as no surprise, as this country was one of the world’s largest producers and consumers of fur (Balakirev & Tinaeva 2001).

### Geographic distribution patterns in Europe

In general, the introduction of a species into few localities, and subsequent further releases, can rapidly lead to the colonisation of large parts of the European continent. We show that, in Europe, the raccoon dog, the muskrat, the American mink, and the raccoon are the most widespread species (in terms of area occupied with established presences), having invaded at least 19 countries each and being present for at least 90 years in the European continent (the most recent invader was the raccoon, introduced in 1927 in Germany). The wide distribution of these species can be attributed to several factors, including adaptability and capacity of colonising different environments (Birnbaum 2013), wide trophic niches (Bartoszewicz 2011), and high reproduction potentials (Pitra et al. 2010). It is of paramount importance to monitor the secondary spread (Essl et al. 2020a) of these species in the European territory and to prevent the establishment of new populations of invasive mammals. Secondary spread would foster alien ranges’ expansion and would counteract the stringent regulations adopted hitherto to prevent new introductions (and mitigate IAS impacts). In the EU, the main drivers of potential impacts of biological invasions (trade and transport, climate change, and socio-economy; Essl et al. 2020b) are highly relevant (Kovats et al. 2014). This, combined with the free circulations of goods and people (Genovesi et al. 2015), may promote a rise of impacts of IAS.

### Environmental and socio-economic impacts in Europe

The wide distribution of alien mammals in the European territory raises many concerns, as they can also transmit diseases to native species, act as a reservoir, and introduce zoonotic pathogens. The latter can be hosted by the majority of the study species and, worryingly, some widespread species carry many of them. Associated infectious diseases such as echinococcosis, toxoplasmosis, and baylisascariasis may pose a serious threat to human health. For comparison, only 11% of the IUCN list of the 100 World’s Worst Invasive Alien Species are reservoirs for zoonotic pathogens (Vila et al. *in press*).

Studies on *E*. *multilocularis* (the pathogen most analysed among all the articles on disease transmission) revolved mainly around raccoon dogs, as they are the definitive hosts (Bagrade et al. 2016), that is the host where the parasite attains sexual maturity. However, muskrats can be intermediate hosts (i.e., a host in which a parasite passes one or more of its asexual stages), and only two studies (out of 14) investigated the prevalence of the pathogen in this rodent. Dedicated health surveillance, in general of these widespread species of invasive mammals, would be beneficial for many people, as the study species are often found in cities or are bred in captivity for commercial purposes.

In this context, the outbreaks of SARS-CoV-2 reported in the Netherlands and in Denmark in 2020 (Molenaar et al. 2020, Oreshkova et al. 2020) are notable. It is currently unknown which role American minks and other wild animals (especially the ones that are regularly in contact with humans, such as stray cats or their prey) may play in SARS-CoV-2 cycle and if they can spread new strains of the virus (mutation affecting the spike protein have already been found in American minks; Molenaar et al. 2020, Oreshkova et al. 2020, WHO 2020) or act as a wild reservoir. This possibility could seriously hinder the vaccination campaigns in Europe. American minks appear to be very susceptible to the virus, and cases are being reported from further countries such as Spain, Sweden, Italy, and the United States. Following the huge outbreaks of SARS-CoV-2, mink industry in the Netherlands and in Sweden will be banned in 2021 (Human Society International 2020, 2021), while Italy and Denmark suspended American mink fur farms activity until the end of 2021 (DW 2020, Ministero della Salute 2021).

Large-scale studies investigating zoonotic (and not) pathogen’s prevalence and the possible roles of invasive mammals of Union concern in their epidemiological cycles are still largely missing. The spread of many pathogens follows similar invasion stages as animals and plants (Vila et al. *in press*). The unknown role of these mammals as a reservoir in the wild could easily jeopardise the efforts in place to prevent, manage or eradicate zoonotic diseases. Due to the many analogies between invasion science and human emerging infectious diseases, the management of IAS can be very useful to tackle future human epidemics (Vila et al. *in press*).

However, predation is probably the most well-known mechanism through which alien species are infamously acknowledged for imperil native biodiversity. This is the case for the American mink, which can exert a negative effect on endangered species such as the water vole (Mori & Mazza 2019) or threaten genetically distinct populations of prey species (Flávio et al. 2020). Heavier egg-predation on ground-nesting birds (compared to previous studies) has recently been reported for the raccoon dog (Dahl & Åhlén 2019), and the muskrat was found to be a major threat to endangered freshwater bivalves in Germany (Stoeckl et al. 2020).

Endangered species (e.g., endemic gastropods) have also been found to be a prey item for the Barbary ground squirrel in the Canary Archipelago (López-Darias et al. 2008). Moreover, this African squirrel disrupts seed dispersal of endemic plant species (Nogales et al. 2005), contributes to invasional meltdown (i.e., an alien species facilitates one another’s invasion; López-Darias & Nogales 2008), and may consume endangered plant species (Bañares et al. 2003). However, all these environmental impacts have been recorded, at present, only in the squirrel’s alien range. The ecology of the other species of invasive mammals in their alien ranges needs to be addressed as soon as possible (Gethöffer & Siebert 2020, Polaina et al. 2020) to understand, contain, and reduce potential environmental impacts.

To summarise, our review illustrates that pet trade is still the main pathway of introduction for alien mammals into Europe. It is currently unclear if the recent decline in first records derived from the stricter measures adopted by the European Union or if it is the result of a saturation effect. To answer this question, longer and accurate monitoring of first records and secondary spread of the invasive mammals of Union concern is necessary. Moreover, the eradication of the study species with a wide distribution is likely unfeasible. However, alien species are not either “bad” or “good”: it is rather the population of the species that has become invasive, that can be problematic (Simberloff et al. 2013) and that should be managed. In this context, the identification of problematic populations or invaded areas may help to mitigate future impacts.

## ACKNOWLEDGEMENTS

The authors appreciate funding by Sapienza University of Rome, the 2017-2018 Belmont Forum and BiodivERsA joint call for research proposals, under the BiodivScen ERA-Net COFUND programme, and with the funding organisations FWF (AlienScenarios, FWF project no I 4011-B32), and the Portuguese National Funds through Fundação para a Ciência e a Tecnologia (CEECIND/02037/2017; UIDB/00295/2020 and UIDP/00295/2020).

## SUPPORTING INFORMATION

Additional supporting information may be found in the online version of this article at the publisher’s web-site.

**Appendix S1.** Process of literature search and keywords used.

**Appendix S2.** Figures illustrating the trends in the published literature, species’ taxonomy, traits, native zoogeographic realms, and pathogens classification.

**Appendix S3.** List of the papers obtained through the literature search process for each study species in Europe.

**Appendix S4.** Year of first record and presences for the study species in Europe.

**Appendix S5.** List of pathogens known to have been recorded to infect the study species in Europe and list of additional references.

## Notes

### Competing Interest Statement

The authors have declared no competing interest.

## REFERENCES

Angulo E, Diagne C, Ballesteros-Mejia L, Adamjy T, Ahmed DA, Akulov E et al. (2021) Non-English languages enrich scientific knowledge: The example of economic costs of biological invasions. Science of The Total Environment: 144441.

Bagrade G, Deksne G, Ozoliņa Z, Howlett SJ, Interisano M, Casulli A, Pozio E (2016) Echinococcus multilocularis in foxes and raccoon dogs: an increasing concern for Baltic countries. Parasites and Vectors 9: 1–9.

Balakirev NA, Tinaeva EA (2001) Fur Farming in Russia: the Current Situation and the Prospects. Scientifur 25: 7–10.

Bañares A, Blanca G, Güemes J, Moreno JC, Ortiz S (2003) Atlas y Libro Rojo de la Flora Vascular Amenazada de España. Dirección General de Conservación de la Naturaleza, Madrid.

Bartoszewicz M (2011) NOBANIS - Invasive Alien Species Fact Sheet - Procyon lotor. Online Database of the European Network on Invasive Alien Species - NOBANIS: 1–9.

Bellard C, Jeschke JM, Leroy B, Mace GM (2018) Insights from modeling studies on how climate change affects invasive alien species geography. Ecology and Evolution 8: 5688–5700.

Bertolino S, Lurz PWW (2013) Callosciurus squirrels: Worldwide introductions, ecological impacts and recommendations to prevent the establishment of new invasive populations. Mammal Review 43: 22– 33.

Biancolini D, Vascellari V, Melone B, Rondinini C (2021) DAMA: the global Distribution of Alien Mammals database. Ecology.

Birnbaum C (2013) NOBANIS - Invasive Alien Species Fact Sheet - Ondatra zibethicus. Online Database of the European Network on Invasive Alien Species - NOBANIS: 1–11.

Blackburn TM, Bellard C, Ricciardi A (2019) Alien versus native species as drivers of recent extinctions. Frontiers in Ecology and the Environment 17: 203–207.

Blackburn TM, Scrivens SL, Heinrich S, Cassey P (2017) Patterns of selectivity in introductions of mammal species worldwide. NeoBiota 33: 33–51.

Bradshaw CJA, Leroy B, Bellard C, Roiz D, Albert C, Fournier A et al. (2016) Massive yet grossly underestimated global costs of invasive insects. Nature Communications 7: 1–8.

Capellini I, Baker J, Allen WL, Street SE, Venditti C (2015) The role of life history traits in mammalian invasion success. Ecology Letters 18: 1099–1107.

Carboneras C, Genovesi P, Vilà M, Blackburn TM, Carrete M, Clavero M et al. (2018) A prioritised list of invasive alien species to assist the effective implementation of EU legislation. Journal of Applied Ecology 55: 539–547.

CBD (2014) Pathways of introduction of invasive species, their prioritisation and management.

CBD (2020) Biodiversity and The 2030 Agenda for Sustainable Development - Policy Brief.

Ćirović D, Toholj D (2016) Distribution of Small Indian Mongoose (Herpestes auropunctatus) in the Eastern Herzegovina – spreading inside mainland. Balkan Journal of Wildlife Research 2: 33–37.

Clout MN, Russell JC (2008) The invasion ecology of mammals: a global perspective. Wildlife Research 35: 180–184.

Collins LM, Warnock ND, Tosh DG, McInnes C, Everest D, Montgomery WI et al. (2014) Squirrelpox virus: Assessing prevalence, transmission and environmental degradation. PLoS ONE 9: 1–8.

Criado-Fornelio A, Martín-Pérez T, Verdú-Expósito C, Reinoso-Ortiz SA, Pérez-Serrano J (2018) Molecular epidemiology of parasitic protozoa and Ehrlichia canis in wildlife in Madrid (central Spain). Parasitology Research 117: 2291–2298.

Dahl F, Åhlén PA (2019) Nest predation by raccoon dog Nyctereutes procyonoides in the archipelago of northern Sweden. Biological Invasions 21: 743–755.

DAISIE (2009) Handbook of Alien Species in Europe, Invading N. Springer, New York.

Diez JM, D’Antonio CM, Dukes JS, Grosholz ED, Olden JD, Sorte CJB et al. (2012) Will extreme climatic events facilitate biological invasions? Frontiers in Ecology and the Environment 10: 249–257.

Duscher T, Hodžić A, Glawischnig W, Duscher GG (2017) The raccoon dog (Nyctereutes procyonoides) and the raccoon (Procyon lotor)—their role and impact of maintaining and transmitting zoonotic diseases in Austria, Central Europe. Parasitology Research 116: 1411–1416.

DW (2020) Danish lawmakers ban mink farming until 2022 amid coronavirus outbreak.

EFSA (2015) Scientific opinion – Update on oral vaccination of foxes and raccoon dogs against rabies. EFSA Journal 13: 70.

Essl F, Latombe G, Lenzner B, Pagad S, Seebens H, Smith K, Wilson JRU, Genovesi P (2020a) The Convention on Biological Diversity (CBD)’s Post-2020 target on invasive alien species – what should it include and how should it be monitored? NeoBiota 121: 99–121.

Essl F, Lenzner B, Bacher S, Bailey S, Capinha C, Daehler C et al. (2020b) Drivers of future alien species impacts: An expert-based assessment. Global Change Biology 26: 4880–4893.

EU (2014) Regulation (EU) No 1143/2014 of the European Parliament and of the Council of 22 October 2014 on the prevention and management of the introduction and spread of invasive alien species. Official Journal of the European Union L317: 35–55.

Flávio H, Caballero P, Jepsen N, Aarestrup K (2020) Atlantic salmon living on the edge: Smolt behaviour and survival during seaward migration in River Minho. Ecology of Freshwater Fish: 1–12.

Genovesi P, Bacher S, Kobelt M, Pascal M, Scalera R (2009) Handbook of Alien Species in Europe. Alien Mammals of Europe.

Genovesi P, Carboneras C, Vilà M, Walton P (2015) EU adopts innovative legislation on invasive species: a step towards a global response to biological invasions? Biological Invasions 17: 1307–1311.

Genovesi P, Carnevali L, Alonzi A, Scalera R (2012) Alien mammals in Europe: Updated numbers and trends, and assessment of the effects on biodiversity. Integrative Zoology 7: 247–253.

Gethöffer F, Siebert U (2020) Current knowledge of the Neozoa Nutria and Muskrat in Europe and their environmental impacts. Journal of Wildlife and Biodiversity 4: 1–12.

Hawkins CL, Bacher S, Essl F, Hulme PE, Jeschke JM, Kühn I et al. (2015) Framework and guidelines for implementing the proposed IUCN Environmental Impact Classification for Alien Taxa (EICAT). Diversity and Distributions 21: 1360–1363.

Hohmann U, Voigt S, Andreas U (2002) Racoons take the offensive. A current assessment. Biologische Invasionen. Herausforderung zum Handeln?: 191–192.

Holt BG, Lessard JP, Borregaard MK, Fritz SA, Araújo MB, Dimitrov D et al. (2013) An Update of Wallace’s Zoogeographic Regions of the World. Science 339: 74–79.

Hulme PE (2009) Trade, transport and trouble: managing invasive species pathways in an era of globalization. Journal of Applied Ecology 46: 10–18.

Human Society International (2020) Dutch mink fur farms to be permanently closed by March 2021 following 41 COVID-19 farm infections.

Human Society International (2021) Sweden suspends mink fur farming in wake of COVID-19.

Invasive Species Ireland (2012) Squirrel pox virus alert.

Invasive Species Ireland (2019) Siberian Chipmunk.

IPBES (2019) Summary for policymakers of the global assessment report on biodiversity and ecosystem services of the Intergovernmental Science-Policy Platform on Biodiversity and Ecosystem Services. IPBES Secretariat, Bonn, Germany.

IUCN (2005) Datasheet on Sciurus carolinensis. Wallingford, United Kingdom.

IUCN (2020) IUCN Red List of Threatened Species.

Karamon J, Kochanowski M, Cencek T, Bartoszewicz M, Kusyk P (2014) Gastrointestinal helminths of raccoons (Procyon lotor) in western Poland (Lubuskie province) - with particular regard to Baylisascaris procyonis. Bulletin of the Veterinary Institute in Pulawy 58: 547–552.

Karamon J, Samorek-Pieróg M, Moskwa B, Rózycki M, Bilska-Zajac E, Zdybel J, Włodarczyk M (2016) Intestinal helminths of raccoon dogs (Nyctereutes procyonoides) and red foxes (Vulpes vulpes) from the Augustów Primeval Forest (north-eastern Poland). Journal of Veterinary Research (Poland) 60: 273–277.

Kornacka A, Cybulska A, Popiołek M, Kuśmierek N, Moskwa B (2018) Survey of Toxoplasma gondii and Neospora caninum in raccoons (Procyon lotor) from the Czech Republic, Germany and Poland. Veterinary Parasitology 262: 47–50.

Kornyushin V V., Malyshko EI, Malega AM (2011) The Helminths of wild predatory mammals of Ukraine. Cestodes. Vestnik Zoologii 45: 4–11.

Kovats RS, Valentini R, Bouwer LM, Georgopoulou E, Jacob D, Martin E, Rounsevell M, Soussana JF (2014) Europe. In: Barros VR, Field CB, Dokken DJ, Mastrandrea MD, Mach KJ, Bilir TE et al. (eds) Climate Change 2014: Impacts, Adaptation and Vulnerability: Part B: Regional Aspects: Working Group II Contribution to the Fifth Assessment Report of the Intergovernmental Panel on Climate Change, 1267–1326. Cambridge University Press, Cambridge, United Kingdom, and New York, NY, USA.

López-Darias M, Nogales M (2008) Effects of the invasive Barbary ground squirrel (Atlantoxerus getulus) on seed dispersal systems of insular xeric environments. Journal of Arid Environments 72: 926–939.

López-Darias M, Lobo JM, Gouat P (2008) Predicting potential distributions of invasive species: The exotic Barbary ground squirrel in the Canarian archipelago and the west Mediterranean region. Biological Invasions 10: 1027–1040.

Louppe V, Leroy B, Herrel A, Veron G (2019) Current and future climatic regions favourable for a globally introduced wild carnivore, the raccoon Procyon lotor. Scientific Reports 9: 1–13.

Louppe V, Leroy B, Herrel A, Veron G (2020) The globally invasive small Indian mongoose Urva auropunctata is likely to spread with climate change. Scientific Reports 10: 1–11.

Mazzamuto MV, Bisi F, Wauters LA, Preatoni DG, Martinoli A (2017) Interspecific competition between alien Pallas’s squirrels and Eurasian red squirrels reduces density of the native species. Biological Invasions 19: 723–735.

Mazzamuto MV, Pisanu B, Romeo C, Ferrari N, Preatoni D, Wauters LA, Chapuis J-L, Martinoli A (2016) Poor Parasite Community of an Invasive Alien Species: Macroparasites of Pallas’s Squirrel in Italy. Annales Zoologici Fennici 53: 103–112.

McFarlane SE, Hunter DC, Senn H V., Smith SL, Holland R, Huisman J, Pemberton JM (2020) Increased genetic marker density reveals high levels of admixture between red deer and introduced Japanese sika in Kintyre, Scotland. Evolutionary Applications 13: 432–441.

Ministero della Salute (2021) Proroga sospensione delle attività degli allevamenti di visoni. Comunicato n. 4.

Moher D, Liberati A, Tetzlaff J, Altman DG, Altman D, Antes G et al. (2009) Preferred reporting items for systematic reviews and meta-analyses: The PRISMA statement. PLoS Medicine 6.

Molenaar RJ, Vreman S, Hakze-van der Honing RW, Zwart R, de Rond J, Weesendorp E et al. (2020) Clinical and Pathological Findings in SARS-CoV-2 Disease Outbreaks in Farmed Mink (Neovison vison). Veterinary Pathology 57: 653–657.

Mori E, Mazza G (2019) Diet of a semiaquatic invasive mammal in northern Italy: Could it be an alarming threat to the endemic water vole? Mammalian Biology 97: 88–94.

Mori E, Zozzoli R, Mazza G (2018) Coming in like a wrecking-ball: are native Eurasian red squirrels displacing invasive Siberian chipmunks? A study from an urban park. Urban Ecosystems 21: 975–981.

Nogales M, Nieves C, Illera JC, Padilla DP, Traveset A (2014) Effect of native and alien vertebrate frugivores patterns of Rubia fruticosa viability and germination in the eastern Canary Islands (Rubiaceae). Functional Ecology 19: 429–436.

Nogales M, Nieves C, Illera JC, Padilla DP, Traveset A (2005) Effect of native and alien vertebrate frugivores on seed viability and germination patterns of Rubia fruticosa (Rubiaceae) in the eastern Canary Islands. Functional Ecology 19: 429–436.

Oksanen A, Siles-Lucas M, Karamon J, Possenti A, Conraths FJ, Romig T et al. (2016) The geographical distribution and prevalence of Echinococcus multilocularis in animals in the European Union and adjacent countries: A systematic review and meta-analysis. Parasites and Vectors 9: 1–23.

Oreshkova N, Moelnaar RJ, Vreman S, Harders F, Munnink BBO, Van Der Honin RWH et al. (2020) SARS-CoV-2 infection in farmed minks, the Netherlands, April and May 2020. Euro Surveillance 25 (23): 1–7.

Pergl J, Brundu G, Harrower CA, Cardoso AC, Genovesi P, Katsanevakis S et al. (2020) Applying the Convention on Biological Diversity Pathway Classification to alien species in Europe. NeoBiota 62: 333– 363.

Pitra C, Schwarz S, Fickel J (2010) Going west-invasion genetics of the alien raccoon dog Nyctereutes procynoides in Europe. European Journal of Wildlife Research 56: 117–129.

Polaina E, Pärt T, Recio MR (2020) Identifying hotspots of invasive alien terrestrial vertebrates in Europe to assist transboundary prevention and control. Scientific Reports: 1–11.

Pyšek P, Richardson DM, Pergl J,Jarošík V, Sixtová Z, Weber E (2008) Geographical and taxonomic biases in invasion ecology. Trends in Ecology and Evolution 23: 237–244.

Ribas MP, Almería S, Fernández-Aguilar X, De Pedro G, Lizarraga P, Alarcia-Alejos O et al. (2018) Tracking Toxoplasma gondii in freshwater ecosystems: interaction with the invasive American mink (Neovison vison) in Spain. Parasitology Research 117: 2275–2281.

Schertler A, Rabitsch W, Moser D, Wessely J, Essl F (2020) The potential current distribution of the coypu (Myocastor coypus) in Europe and climate change induced shifts in the near future. NeoBiota 58: 129–160.

Seebens H, Bacher S, Blackburn TM, Capinha C, Dawson W, Dullinger S et al. (2020) Projecting the continental accumulation of alien species through to 2050. Global Change Biology: 1–13.

Seebens H, Blackburn TM, Dyer EE, Genovesi P, Hulme PE, Jeschke JM et al. (2017) No saturation in the accumulation of alien species worldwide. Nature Communications 8.

Signorile AL, Reuman DC, Lurz PWW, Bertolino S, Carbone C, Wang J (2016) Using DNA profiling to investigate human-mediated translocations of an invasive species. Biological Conservation 195: 97– 105.

Simberloff D, Martin JL, Genovesi P, Maris V, Wardle DA, Aronson J et al. (2013) Impacts of biological invasions: What’s what and the way forward. Trends in Ecology and Evolution 28: 58–66.

Soria CD, Di Marco M, Pacifici M, Butchart SHM, Rondinini C (2021) COMBINE: A Coalesced Mammal Database of Intrinsic and Extrinsic traits. Ecology.

Šprem N, Zachos FE (2020) Axis Deer Axis axis Erxleben, 1777. In: Hackländer K, Zachos FE (eds) Handbook of the Mammals of Europe, 1–9. Springer Nature Switzerland.

Stoeckl K, Denic M, Geist J (2020) Conservation status of two endangered freshwater mussel species in Bavaria, Germany: Habitat quality, threats, and implications for conservation management. Aquatic Conservation: Marine and Freshwater Ecosystems 30: 647–661.

Vila M, Dunn A, Essl F, Gòmez-Dìaz E, Hulme PE, Jeschke JM et al. Viewing emerging human infectious disease epidemics through the lens of invasion science. BioScience.

Vilà M, Hulme PE (eds) (2017) Impact of biological invasions on ecosystem services, Invading N. Springer.

Wahlström H, Lindberg A, Lindh J, Wallensten A, Lindqvist R, Plym-Forshell L et al. (2012) Investigations and actions taken during 2011 due to the first finding of echinococcus multilocularis in Sweden. Eurosurveillance 17: 1–7.

West HER, Capellini I (2016) Male care and life history traits in mammals. Nature Communications 7.

WHO (2020) SARS-CoV-2 mink-associated variant strain – Denmark.

