## Appendix_S1_S2_S3_S5 for "Invasive alien mammals of European Union concern"

**Appendix S1.** Process of literature search and keywords used.

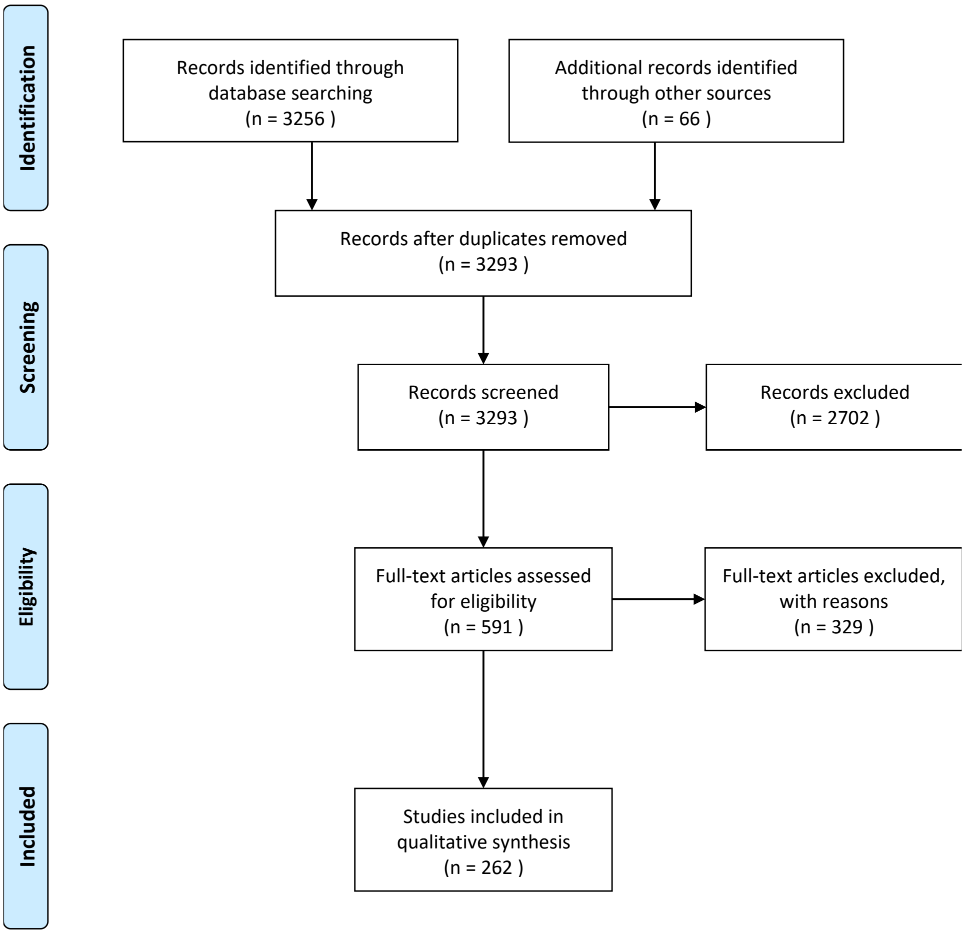


**Fig. S1.** The flowchart illustrating the process of literature search and review, based on PRISMA guidelines, conducted in August and September 2020 (adapted from Moher et al., 2009).

### Scopus and Web of Science search terms used to review the literature for each study species in the study area.

*Atlantoxerus getulus*

( TITLE-ABS-KEY ( "Atlantoxerus getulus" OR "Barbary ground squirrel" ) AND TITLE-ABS-KEY ( europe* OR "european union" OR EU OR Spain OR introduc* OR invasi* OR establish* OR alien OR invasive OR ias OR allochthonous OR exotic OR "Aichi target 9" OR "EU biodiversity strategy" OR "europe* biodiversity strategy" OR "EU IAS regulation" OR "Europe* IAS regulation" OR “Union List” OR "propagule pressure" OR "colonization pressure" OR "life-history trait*" OR "life history trait*" OR trait* OR "risk assessment*" OR "impact assessment*" OR "environmental impact*" OR "socio-economic impact*" OR "socio economic impact*" OR "economic impact*" ) ) AND ( LIMIT-TO ( SUBJAREA , "AGRI" ) OR LIMIT-TO ( SUBJAREA , "ENVI" ) OR LIMIT-TO ( SUBJAREA , "EART" ) ) AND ( LIMIT-TO ( LANGUAGE , "English" ) )

*Axis axis*

( TITLE-ABS-KEY ( "Axis axis" OR "Indian spotted deer" OR chital* OR “Spotted deer” OR “Axis deer”) AND TITLE-ABS-KEY ( europe* OR "european union" OR EU OR Croatia OR Ukraine OR introduc* OR invasi* OR establish* OR alien OR invasive OR ias OR allochthonous OR exotic OR "Aichi target 9" OR "EU biodiversity strategy" OR "europe* biodiversity strategy" OR "EU IAS regulation" OR "Europe* IAS regulation" OR “Union List” OR "propagule pressure" OR "colonization pressure" OR "life-history trait*" OR "life history trait*" OR trait* OR "risk assessment*" OR "impact assessment*" OR "environmental impact*" OR "socio-economic impact*" OR "socio economic impact*" OR "economic impact*" ) ) AND ( LIMIT-TO ( SUBJAREA , "AGRI" ) OR LIMIT-TO ( SUBJAREA , "ENVI" ) OR LIMIT-TO ( SUBJAREA , "EART" ) ) AND ( LIMIT-TO ( LANGUAGE , "English" ) )

*Callosciurus erythraeus*

( TITLE-ABS-KEY ( "Callosciurus erythraeus" OR "Pallas's squirrel" ) AND TITLE-ABS-KEY ( europe* OR "european union" OR EU OR Belgium OR France OR Germany OR Italy OR Netherlands OR “The Netherlands” OR introduc* OR invasi* OR establish* OR alien OR invasive OR ias OR allochthonous OR exotic OR "Aichi target 9" OR "EU biodiversity strategy" OR "europe* biodiversity strategy" OR "EU IAS regulation" OR "Europe* IAS regulation" OR “Union List” OR "propagule pressure" OR "colonization pressure" OR "life-history trait*" OR "life history trait*" OR trait* OR "risk assessment*" OR "impact assessment*" OR "environmental impact*" OR "socio-economic impact*" OR "socio economic impact*" OR "economic impact*" ) ) AND ( LIMIT-TO ( SUBJAREA , "AGRI" ) OR LIMIT-TO ( SUBJAREA , "ENVI" ) OR LIMIT-TO ( SUBJAREA , "EART" ) ) AND ( LIMIT-TO ( LANGUAGE , "English" ) ) AND ( LIMIT-TO ( PUBYEAR , 2020 ) OR LIMIT-TO ( PUBYEAR , 2019 ) OR LIMIT-TO ( PUBYEAR , 2018 ) OR LIMIT-TO ( PUBYEAR , 2017 ) OR LIMIT-TO ( PUBYEAR , 2016 ) OR LIMIT-TO ( PUBYEAR , 2015 ) OR LIMIT-TO ( PUBYEAR , 2014 ))

*Callosciurus finlaysonii*

( TITLE-ABS-KEY ( "Callosciurus finlaysonii " OR "Variable squirrel" OR “Finlayson’s squirrel”) AND TITLE-ABS-KEY ( europe* OR "european union" OR EU OR Italy OR introduc* OR invasi* OR establish* OR alien OR invasive OR ias OR allochthonous OR exotic OR "Aichi target 9" OR "EU biodiversity strategy" OR "europe* biodiversity strategy" OR "EU IAS regulation" OR "Europe* IAS regulation" OR “Union List” OR "propagule pressure" OR "colonization pressure" OR "life-history trait*" OR "life history trait*" OR trait* OR "risk assessment*" OR "impact assessment*" OR "environmental impact*" OR "socio-economic impact*" OR "socio economic impact*" OR "economic impact*" ) ) AND ( LIMIT-TO ( SUBJAREA , "AGRI" ) OR LIMIT-TO ( SUBJAREA , "ENVI" ) OR LIMIT-TO ( SUBJAREA , "EART" ) ) AND ( LIMIT-TO ( LANGUAGE , "English" ) ) AND ( LIMIT-TO ( PUBYEAR , 2020 ) OR LIMIT-TO ( PUBYEAR , 2019 ) OR LIMIT-TO ( PUBYEAR , 2018 ) )

*Castor canadensis*

[excluding] WEB OF SCIENCE CATEGORIES: ( ANTHROPOLOGY OR MATERIALS SCIENCE MULTIDISCIPLINARY OR MATHEMATICAL COMPUTATIONAL BIOLOGY OR OPHTHALMOLOGY OR ORTHOPEDICS OR LINGUISTICS OR PHYSICS FLUIDS PLASMAS OR THERMODYNAMICS OR MATHEMATICS APPLIED OR COMPUTER SCIENCE ARTIFICIAL INTELLIGENCE OR ENGINEERING ELECTRICAL ELECTRONIC OR COMPUTER SCIENCE SOFTWARE ENGINEERING OR DENTISTRY ORAL SURGERY MEDICINE OR HISTORY OR HUMANITIES MULTIDISCIPLINARY OR MECHANICS OR LANGUAGE LINGUISTICS OR EMERGENCY MEDICINE OR COMPUTER SCIENCE INTERDISCIPLINARY APPLICATIONS OR PHYSICS MATHEMATICAL OR COMPUTER SCIENCE THEORY METHODS OR SURGERY OR ART OR CARDIAC CARDIOVASCULAR SYSTEMS OR HEALTH CARE SCIENCES SERVICES OR HISTORY PHILOSOPHY OF SCIENCE OR MATHEMATICS INTERDISCIPLINARY APPLICATIONS OR COMPUTER SCIENCE INFORMATION SYSTEMS OR EDUCATION EDUCATIONAL RESEARCH OR EDUCATION SCIENTIFIC DISCIPLINES OR ENERGY FUELS OR INSTRUMENTS INSTRUMENTATION OR INTERNATIONAL RELATIONS OR HOSPITALITY LEISURE SPORT TOURISM OR PUBLIC ENVIRONMENTAL OCCUPATIONAL HEALTH ) ( TITLE-ABS-KEY ( "Castor canadensis" OR beaver* OR “American beaver”) AND TITLE-ABS-

KEY ( europe* OR "european union" OR EU OR Belgium OR Finland OR France OR Germany OR Luxembourg OR Russia OR “Russian Federation” OR introduc* OR invasi* OR establish* OR alien OR invasive OR ias OR allochthonous OR exotic OR "Aichi target 9" OR "EU biodiversity strategy" OR "europe* biodiversity strategy" OR "EU IAS regulation" OR "Europe* IAS regulation" OR “Union List” OR "propagule pressure" OR "colonization pressure" OR "life-history trait*" OR "life history trait*" OR trait* OR "risk assessment*" OR "impact assessment*" OR "environmental impact*" OR "socio-economic impact*" OR "socio economic impact*" OR "economic impact*" ) ) AND ( LIMIT-TO ( SUBJAREA , "AGRI" ) OR LIMIT-TO ( SUBJAREA , "ENVI" ) OR LIMIT-TO ( SUBJAREA , "EART" ) ) AND ( LIMIT-TO ( LANGUAGE , "English" ) ) AND ( LIMIT-TO ( PUBYEAR , 2020 ) OR LIMIT-TO ( PUBYEAR , 2019 ) OR LIMIT-TO ( PUBYEAR , 2018 ) OR LIMIT-TO ( PUBYEAR , 2017 ) OR LIMIT-TO ( PUBYEAR , 2016 ) OR LIMIT-TO ( PUBYEAR , 2015 ) OR LIMIT-TO ( PUBYEAR , 2014 ) OR LIMIT-TO ( PUBYEAR , 2013 ) OR LIMIT-TO ( PUBYEAR , 2012 ) OR LIMIT-TO ( PUBYEAR , 2011 ) OR LIMIT-TO ( PUBYEAR , 2010 ) )

*Cervus nippon*

( TITLE-ABS-KEY ( "Cervus nippon" OR “Sika deer”) AND TITLE-ABS-KEY ( europe* OR "european union" OR EU OR Austria OR Czechia OR “Czech Republic” OR Denmark OR Finland OR France OR Germany OR Hungary OR Ireland OR Lithuania OR Poland OR Russia OR “Russian Federation” OR Switzerland OR “United Kingdom” OR UK OR Ukraine OR introduc* OR invasi* OR establish* OR alien OR invasive OR ias OR allochthonous OR exotic OR "Aichi target 9" OR "EU biodiversity strategy" OR "europe* biodiversity strategy" OR "EU IAS regulation" OR "Europe* IAS regulation" OR “Union List” OR "propagule pressure" OR "colonization pressure" OR "life-history trait*" OR "life history trait*" OR trait* OR "risk assessment*" OR "impact assessment*" OR "environmental impact*" OR "socio-economic impact*" OR "socio economic impact*" OR "economic impact*" ) ) AND ( LIMIT-TO ( SUBJAREA , "AGRI" ) OR LIMIT-TO ( SUBJAREA , "ENVI" ) OR LIMIT-TO ( SUBJAREA , "EART" ) ) AND ( LIMIT-TO ( LANGUAGE , "English" ) ) AND ( LIMIT-TO ( PUBYEAR , 2020 ) OR LIMIT-TO ( PUBYEAR , 2019 ) OR LIMIT-TO ( PUBYEAR , 2018 ) OR LIMIT-TO ( PUBYEAR , 2017 ) OR LIMIT-TO ( PUBYEAR , 2016 ) OR LIMIT-TO ( PUBYEAR , 2015 ) OR LIMIT-TO ( PUBYEAR , 2014 ) OR LIMIT-TO ( PUBYEAR , 2013 ) OR LIMIT-TO ( PUBYEAR , 2012 ) OR LIMIT-TO ( PUBYEAR , 2011 ) OR LIMIT-TO ( PUBYEAR , 2010 ) OR LIMIT-TO ( PUBYEAR , 2009 ) )

*Eutamias sibiricus*

( TITLE-ABS-KEY ( "Eutamias sibiricus" OR “Tamias sibiricus” OR "Siberian chipmunk" ) AND TITLE-ABS-KEY ( europe* OR "european union" OR EU OR Belgium OR Denmark OR France OR Germany OR Ireland OR Italy OR Netherlands OR “The Netherlands” OR Russia OR “Russian Federation” OR Spain OR Switzerland OR “United Kingdom” OR UK OR introduc* OR invasi* OR establish* OR alien OR invasive OR ias OR allochthonous OR exotic OR "Aichi target 9" OR "EU biodiversity strategy" OR "europe* biodiversity strategy" OR "EU IAS regulation" OR "Europe* IAS regulation" OR “Union List” OR "propagule pressure" OR "colonization pressure" OR "life-history trait*" OR "life history trait*" OR trait* OR "risk assessment*" OR "impact assessment*" OR "environmental impact*" OR "socio-economic impact*" OR "socio economic impact*" OR "economic impact*" ) ) AND ( LIMIT-TO ( SUBJAREA , "AGRI" ) OR LIMIT-TO ( SUBJAREA , "ENVI" ) OR LIMIT-TO ( SUBJAREA , "EART" ) ) AND ( LIMIT-TO ( LANGUAGE , "English" ) )

*Herpestes auropunctatus*

( TITLE-ABS-KEY ("Herpestes javanic*" OR "Herpestes auropunctat*" OR "Urva javanic*" OR "Urva auropunctat*" OR "Small Indian mongoose" ) AND TITLE-ABS-KEY ( europe* OR "european union" OR EU OR Albania OR “Bosnia and Herzegovina” OR “Bosnia-Herzegovina” OR Croatia OR Montenegro OR introduc* OR invasi* OR establish* OR alien OR invasive OR ias OR allochthonous OR exotic OR "Aichi target 9" OR "EU biodiversity strategy" OR "europe* biodiversity strategy" OR "EU IAS regulation" OR "Europe* IAS regulation" OR “Union List” OR "propagule pressure" OR "colonization pressure" OR "life-history trait*" OR "life history trait*" OR trait* OR "risk assessment*" OR "impact assessment*" OR "environmental impact*" OR "socio-economic impact*" OR "socio economic impact*" OR "economic impact*" ) ) AND ( LIMIT-TO ( SUBJAREA , "AGRI" ) OR LIMIT-TO ( SUBJAREA , "ENVI" ) OR LIMIT-TO ( SUBJAREA , "EART" ) ) AND ( LIMIT-TO ( LANGUAGE , "English" ) ) AND ( LIMIT-TO ( PUBYEAR , 2020 ) OR LIMIT-TO ( PUBYEAR , 2019 ) OR LIMIT-TO ( PUBYEAR , 2018 ) OR LIMIT-TO ( PUBYEAR , 2017 ) OR LIMIT-TO ( PUBYEAR , 2016 ) OR LIMIT-TO ( PUBYEAR , 2015 ) )

*Muntiacus reevesi*

( TITLE-ABS-KEY ( "Muntiacus reevesi" OR “Reeves’ muntjac” OR "Reeves muntjac" ) AND TITLE-ABS-KEY ( europe* OR "european union" OR EU OR Belgium OR Denmark OR Ireland OR Netherlands OR “The Netherlands” OR “United Kingdom” OR UK OR introduc* OR invasi* OR establish* OR alien OR invasive OR ias OR allochthonous OR exotic OR "Aichi target 9" OR "EU biodiversity strategy" OR "europe* biodiversity strategy" OR "EU IAS regulation" OR “Union List” OR "Europe* IAS regulation" OR "propagule pressure" OR "colonization pressure" OR "life-history trait*" OR "life history trait*" OR trait* OR "risk assessment*" OR "impact assessment*" OR "environmental impact*" OR "socio-economic impact*" OR "socio economic impact*" OR "economic impact*" ) ) AND ( LIMIT-TO ( SUBJAREA , "AGRI" ) OR LIMIT-TO ( SUBJAREA , "ENVI" ) OR LIMIT-TO ( SUBJAREA , "EART" ) ) AND ( LIMIT-TO ( LANGUAGE , "English" ) )

*Myocastor coypus*

( TITLE-ABS-KEY ( "Myocastor coypus" OR “coypu*” OR “nutria”) AND TITLE-ABS-KEY ( europe* OR "european union" OR EU OR Austria OR Belarus OR Belgium OR Bulgaria OR Croatia OR Czechia OR “Czech Republic” OR Denmark OR France OR Germany OR Greece OR Hungary OR Ireland OR Italy OR Luxembourg OR Macedonia OR Montenegro OR Netherlands OR “The Netherlands” OR Norway OR Poland OR Romania OR Serbia OR Slovakia OR Slovenia OR Spain OR Sweden OR Switzerland OR “United Kingdom” OR UK OR Ukraine OR introduc* OR invasi* OR establish* OR alien OR invasive OR ias OR allochthonous OR exotic OR "Aichi target 9" OR "EU biodiversity strategy" OR "europe* biodiversity strategy" OR "EU IAS regulation" OR "Europe* IAS regulation" OR “Union List” OR "propagule pressure" OR "colonization pressure" OR "life-history trait*" OR "life history trait*" OR trait* OR "risk assessment*" OR "impact assessment*" OR "environmental impact*" OR "socio-economic impact*" OR "socio economic impact*" OR "economic impact*" ) ) AND ( LIMIT-TO ( SUBJAREA , "AGRI" ) OR LIMIT-TO ( SUBJAREA , "ENVI" ) OR LIMIT-TO ( SUBJAREA , "EART" ) ) AND ( LIMIT-TO ( LANGUAGE , "English" ) ) AND ( LIMIT-TO ( PUBYEAR , 2020 ) OR LIMIT-TO ( PUBYEAR , 2019 ) OR LIMIT-TO ( PUBYEAR , 2018 ) OR LIMIT-TO ( PUBYEAR , 2017 ) OR LIMIT-TO ( PUBYEAR , 2016 ) OR LIMIT-TO ( PUBYEAR , 2015 ) OR LIMIT-TO ( PUBYEAR , 2014 ) )

*Nasua nasua*

( TITLE-ABS-KEY ( "Nasua nasua" OR “South American coati” OR “ring-tailed coati” ) AND TITLE-ABS-KEY ( europe* OR "european union" OR EU OR Belgium OR France OR Germany OR Spain OR introduc* OR invasi* OR establish* OR alien OR invasive OR ias OR allochthonous OR exotic OR "Aichi target 9" OR "EU biodiversity strategy" OR "europe* biodiversity strategy" OR "EU IAS regulation" OR "Europe* IAS regulation" OR “Union List” OR "propagule pressure" OR "colonization pressure" OR "life-history trait*" OR "life history trait*" OR trait* OR "risk assessment*" OR "impact assessment*" OR "environmental impact*" OR "socio-economic impact*" OR "socio economic impact*" OR "economic impact*" ) ) AND ( LIMIT-TO ( SUBJAREA , "AGRI" ) OR LIMIT-TO ( SUBJAREA , "ENVI" ) OR LIMIT-TO ( SUBJAREA , "EART" ) ) AND ( LIMIT-TO ( LANGUAGE , "English" ) ) AND ( LIMIT-TO ( PUBYEAR , 2020 ) OR LIMIT-TO ( PUBYEAR , 2019 ) OR LIMIT-TO ( PUBYEAR , 2018 ) OR LIMIT-TO ( PUBYEAR , 2017 ) OR LIMIT-TO ( PUBYEAR , 2016 ) OR LIMIT-TO ( PUBYEAR , 2015 ) )

*Neovison vison*

( TITLE-ABS-KEY ( "Neovison vison" OR “American mink”) AND TITLE-ABS-KEY ( europe* OR "european union" OR EU OR Albania OR Andorra OR Austria OR Belarus OR Belgium OR Czechia OR “Czech Republic” OR Denmark OR Estonia OR Finland OR France OR Germany OR Greece OR Hungary OR Iceland OR Ireland OR Italy OR Latvia OR Lithuania OR Luxembourg OR Macedonia OR “North Macedonia” OR Montenegro OR Netherlands OR “The Netherlands” OR Norway OR Poland OR Portugal OR Romania OR Russia OR “Russian federation” OR Slovakia OR Slovenia OR Serbia OR Spain OR Sweden OR Switzerland OR “United Kingdom” OR UK OR Ukraine OR introduc* OR invasi* OR establish* OR alien OR invasive OR ias OR allochthonous OR exotic OR "Aichi target 9" OR "EU biodiversity strategy" OR "europe* biodiversity strategy" OR "EU IAS regulation" OR "Europe* IAS regulation" OR “Union List” OR "propagule pressure" OR "colonization pressure" OR "life-history trait*" OR "life history trait*" OR trait* OR "risk assessment*" OR "impact assessment*" OR "environmental impact*" OR "socio-economic impact*" OR "socio economic impact*" OR "economic impact*" ) ) AND ( LIMIT-TO ( SUBJAREA , "AGRI" ) OR LIMIT-TO ( SUBJAREA , "ENVI" ) OR LIMIT-TO ( SUBJAREA , "EART" ) ) AND ( LIMIT-TO ( LANGUAGE , "English" ) ) AND ( LIMIT-TO ( PUBYEAR , 2020 ) OR LIMIT-TO ( PUBYEAR , 2019 ) OR LIMIT-TO ( PUBYEAR , 2018 ) OR LIMIT-TO ( PUBYEAR , 2017 ) OR LIMIT-TO ( PUBYEAR , 2016 ) )

*Nyctereutes procyonoides*

( TITLE-ABS-KEY ( "Nyctereutes procyonoides" OR “Racoon dog*” ) AND TITLE-ABS-KEY ( europe* OR "european union" OR EU OR Albania OR Austria OR Belarus OR Belgium OR “Bosnia and Herzegovina” OR “Bosnia-Herzegovina” OR Bulgaria OR Croatia OR Czechia OR “Czech Republic” OR Denmark OR Estonia OR Finland OR France OR Germany OR Greece OR Hungary OR Italy OR Latvia OR Liechtenstein OR Lithuania OR Luxembourg OR Macedonia OR “North Macedonia” OR Moldova OR Montenegro OR Netherlands OR “The Netherlands” OR Norway OR Poland OR Romania OR Russia OR “Russian Federation” OR Serbia OR Slovakia OR Slovenia OR Sweden OR Switzerland OR Ukraine OR introduc* OR invasi* OR establish* OR alien OR invasive OR ias OR allochthonous OR exotic OR "Aichi target 9" OR "EU biodiversity strategy" OR "europe* biodiversity strategy" OR "EU IAS regulation" OR "Europe* IAS regulation" OR “Union List” OR "propagule pressure" OR "colonization pressure" OR "life-history trait*" OR "life history trait*" OR trait* OR "risk assessment*" OR "impact assessment*" OR "environmental impact*" OR "socio-economic impact*" OR "socio economic impact*" OR "economic impact*" ) ) AND ( LIMIT-TO ( SUBJAREA , "AGRI" ) OR LIMIT-TO ( SUBJAREA , "ENVI" ) OR LIMIT-TO ( SUBJAREA , "EART" ) ) AND ( LIMIT-TO ( LANGUAGE , "English" ) ) AND ( LIMIT-TO ( PUBYEAR , 2020 ) OR LIMIT-TO ( PUBYEAR , 2019 ) OR LIMIT-TO ( PUBYEAR , 2018 ) OR LIMIT-TO ( PUBYEAR , 2017 ) OR LIMIT-TO ( PUBYEAR , 2016 ) OR LIMIT-TO ( PUBYEAR , 2015 ) )

*Ondatra zibethicus*

( TITLE-ABS-KEY ( "Ondatra zibethicus" OR muskrat* ) AND TITLE-ABS-KEY ( europe* OR "european union" OR EU OR Andorra OR Austria OR Belarus OR Belgium OR “Bosnia and Herzegovina” OR “Bosnia-Herzegovina” OR Bulgaria OR Croatia OR Czechia OR “Czech Republic” OR Denmark OR Estonia OR Finland OR France OR Germany OR Greece OR Hungary OR Ireland OR Italy OR Latvia OR Liechtenstein OR Lithuania OR Luxembourg OR Moldova OR Montenegro OR Netherlands OR “The Netherlands” OR Norway OR Poland OR Romania OR Russia OR “Russian Federation” OR Serbia OR Slovakia OR Slovenia OR Spain OR Sweden OR Switzerland OR “United Kingdom” OR UK OR Ukraine OR introduc* OR invasi* OR establish* OR alien OR invasive OR ias OR allochthonous OR exotic OR "Aichi target 9" OR "EU biodiversity strategy" OR "europe* biodiversity strategy" OR "EU IAS regulation" OR "Europe* IAS regulation" OR “Union List” OR "propagule pressure" OR "colonization pressure" OR "life-history trait*" OR "life history trait*" OR trait* OR "risk assessment*" OR "impact assessment*" OR "environmental impact*" OR "socio-economic impact*" OR "socio economic impact*" OR "economic impact*" ) ) AND ( LIMIT-TO ( SUBJAREA , "AGRI" ) OR LIMIT-TO

( SUBJAREA , "ENVI" ) OR LIMIT-TO ( SUBJAREA , "EART" ) ) AND ( LIMIT-TO ( LANGUAGE , "English" ) ) AND ( LIMIT-TO ( PUBYEAR , 2020 ) OR LIMIT-TO ( PUBYEAR , 2019 ) OR LIMIT-TO ( PUBYEAR , 2018 ) OR LIMIT-TO ( PUBYEAR , 2017 ) OR LIMIT-TO ( PUBYEAR , 2016 ) OR LIMIT-TO ( PUBYEAR , 2015 ) )

*Procyon lotor*

( TITLE-ABS-KEY ( "Procyon lotor" OR raccoon* OR "Northern raccoon" AND NOT "raccoon dog") AND TITLE-ABS-KEY ( europe* OR "european union" OR EU OR Austria OR Belarus OR Belgium OR Croatia OR Czechia OR "Czech Republic" OR Denmark OR Estonia OR France OR Germany OR Hungary OR Ireland OR Italy OR Liechtenstein OR Lithuania OR Luxembourg OR Netherlands OR "The Netherlands" OR Poland OR Romania OR Russia OR "Russian Federation" OR Serbia OR Slovakia OR Slovenia OR Spain OR Switzerland OR Ukraine OR introduc* OR invasi* OR establish* OR alien OR invasive OR ias OR allochthonous OR exotic OR "Aichi target 9" OR "EU biodiversity strategy" OR "europe* biodiversity strategy" OR "EU IAS regulation" OR "Europe* IAS regulation" OR "Union List" OR "propagule pressure" OR "colonization pressure" OR "life-history trait*" OR "life history trait*" OR trait* OR "risk assessment*" OR "impact assessment*" OR "environmental impact*" OR "socio-economic impact*" OR "socio economic impact*" OR "economic impact*" ) ) AND ( LIMIT-TO ( SUBJAREA , "AGRI" ) OR LIMIT-TO ( SUBJAREA , "ENVI" ) OR LIMIT-TO ( SUBJAREA , "EART" ) ) AND ( LIMIT-TO ( LANGUAGE , "English" ) ) AND ( LIMIT-TO ( SUBJAREA , "AGRI" ) OR LIMIT-TO ( SUBJAREA , "ENVI" ) OR LIMIT-TO ( SUBJAREA , "EART" ) ) AND ( LIMIT-TO ( LANGUAGE , "English" ) ) AND ( LIMIT-TO ( PUBYEAR , 2020 ) OR LIMIT-TO ( PUBYEAR , 2019 ) OR LIMIT-TO ( PUBYEAR , 2018 ) OR LIMIT-TO ( PUBYEAR , 2017 ) OR LIMIT-TO ( PUBYEAR , 2016 ) OR LIMIT-TO ( PUBYEAR , 2015 ) OR LIMIT-TO ( PUBYEAR , 2014 ) OR LIMIT-TO ( PUBYEAR , 2013 ) OR LIMIT-TO ( PUBYEAR , 2012 ) OR LIMIT-TO ( PUBYEAR , 2011 ) )

*Sciurus carolinensis*

( TITLE-ABS-KEY ( "*Sciurus carolinensis*" OR "Eastern gr*y squirrel" OR "American gr*y squirrel" OR "gr*y squirrel") AND TITLE-ABS-KEY ( europe* OR "european union" OR EU OR Belgium OR Germany OR Ireland OR Italy OR Netherlands OR "The Netherlands" OR "United Kingdom" OR UK OR introduc* OR invasi* OR establish* OR alien OR invasive OR ias OR allochthonous OR exotic OR "Aichi target 9" OR "EU biodiversity strategy" OR "europe* biodiversity strategy" OR "EU IAS regulation" OR "Europe* IAS regulation" OR "Union List" OR "propagule pressure" OR "colonization pressure" OR "life-history trait*" OR "life history trait*" OR trait* OR "risk assessment*" OR "impact assessment*" OR "environmental impact*" OR "socio-economic impact*" OR "socio economic impact*" OR "economic impact*" ) ) AND ( LIMIT-TO ( SUBJAREA , "AGRI" ) OR LIMIT-TO ( SUBJAREA , "ENVI" ) OR LIMIT-TO ( SUBJAREA , "EART" ) ) AND ( LIMIT-TO ( LANGUAGE , "English" ) ) AND ( LIMIT-TO ( PUBYEAR , 2020 ) OR LIMIT-TO ( PUBYEAR , 2019 ) OR LIMIT-TO ( PUBYEAR , 2018 ) OR LIMIT-TO ( PUBYEAR , 2017 ) OR LIMIT-TO ( PUBYEAR , 2016 ) OR LIMIT-TO ( PUBYEAR , 2015 ) )

**Appendix S2.** Figures illustrating the trends in the published literature, species' taxonomy, traits, native zoogeographic realms, and pathogens classification.

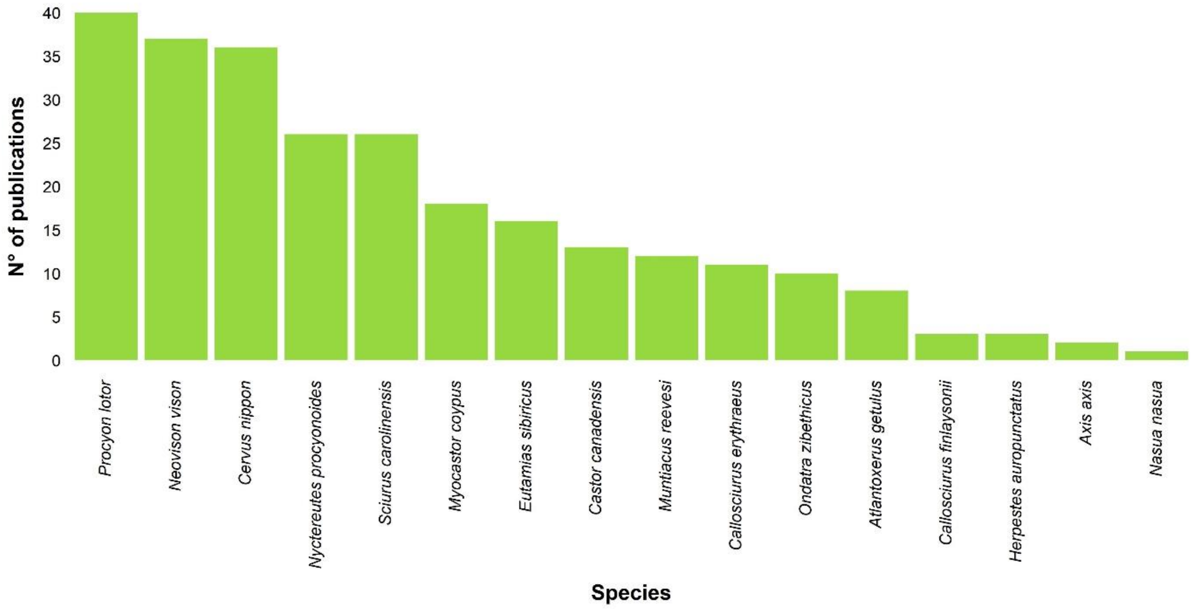


**Fig. S2.** The number of publications resulting from the literature search process collected for each study species in Europe (n = 261; one publication has no date).

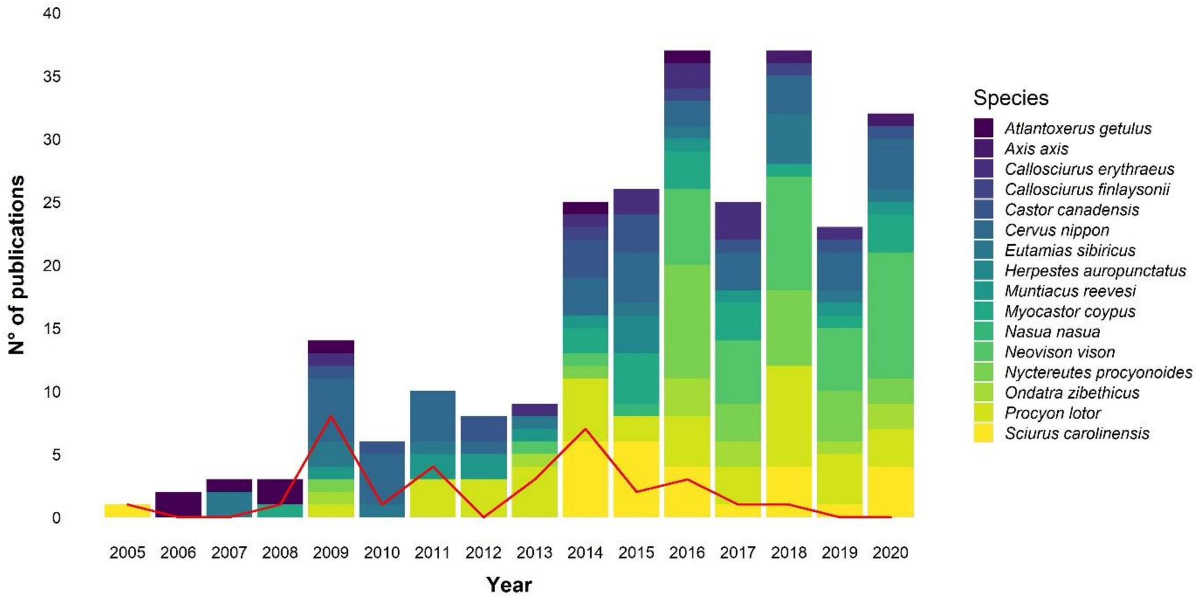


**Fig. S3.** The number of published studies per year for each study species from 2005 to 2020 (n = 261; one publication has no date). The line shows the overall temporal trend of the published datasheets.

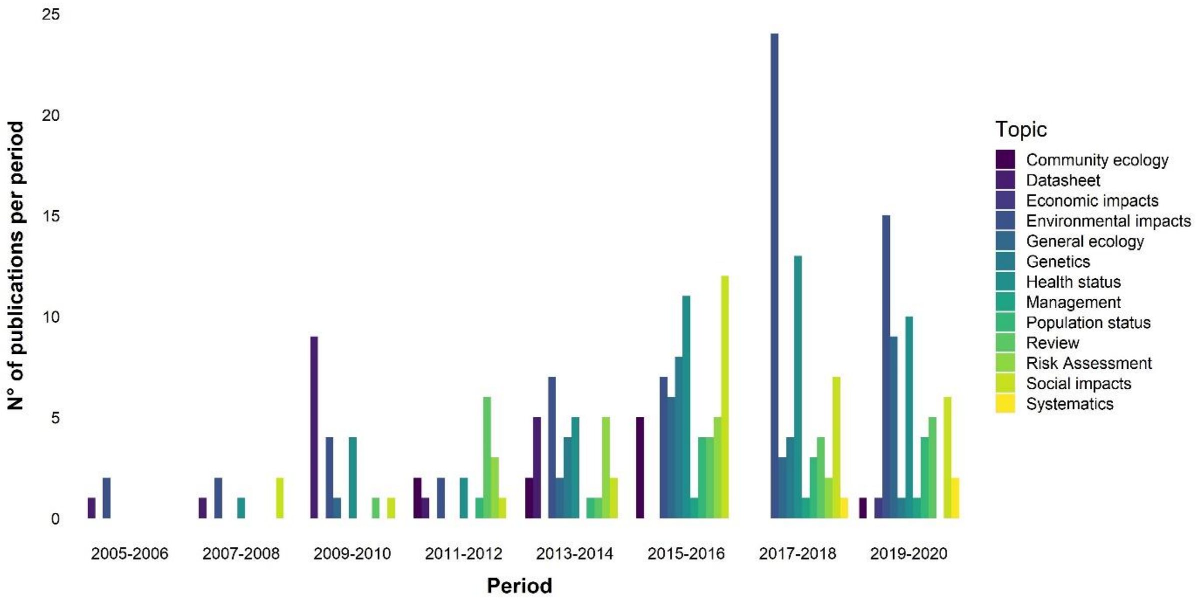


**Fig. S4.** The number of publications for each topic over a two-year period from 2005 to 2020 (n = 261; one publication has no date).

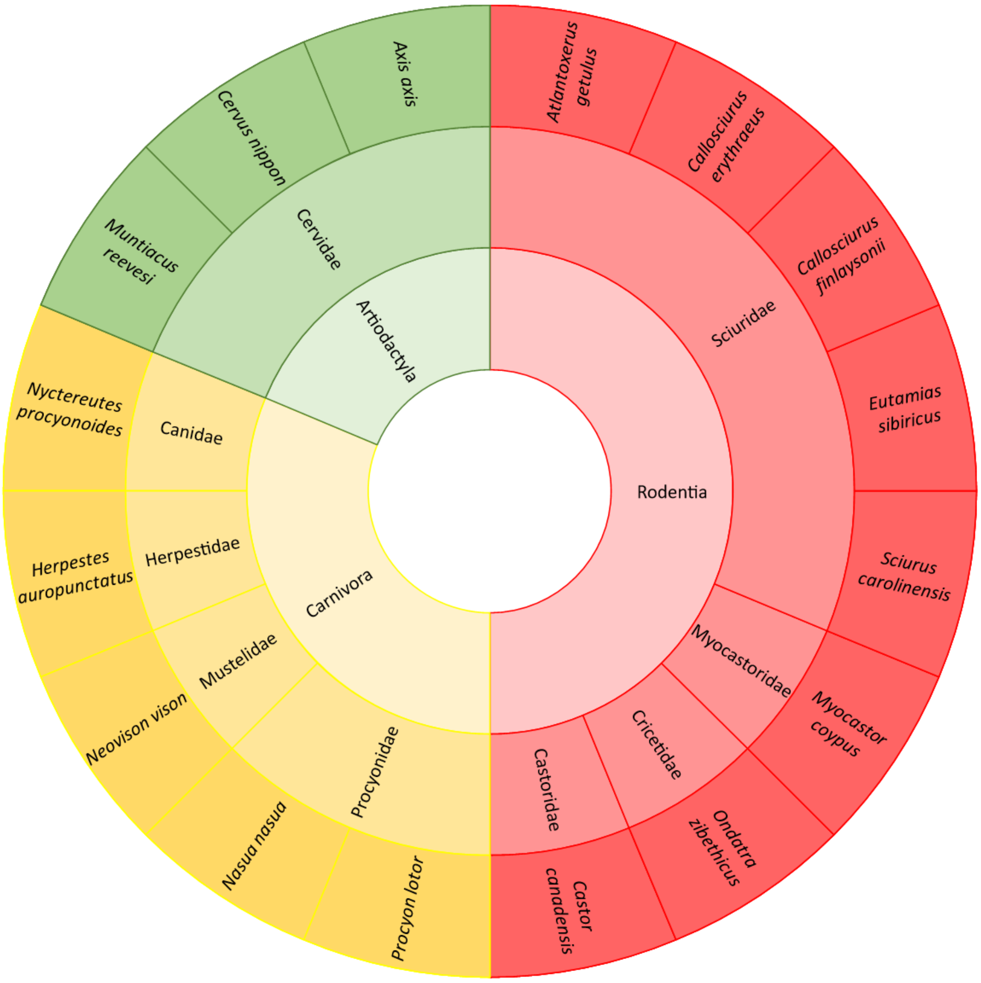


**Fig. S5.** Taxonomic assignment of the study species (n = 16). The inner circle represents the orders, the middle circle the families and the outer circle the species.

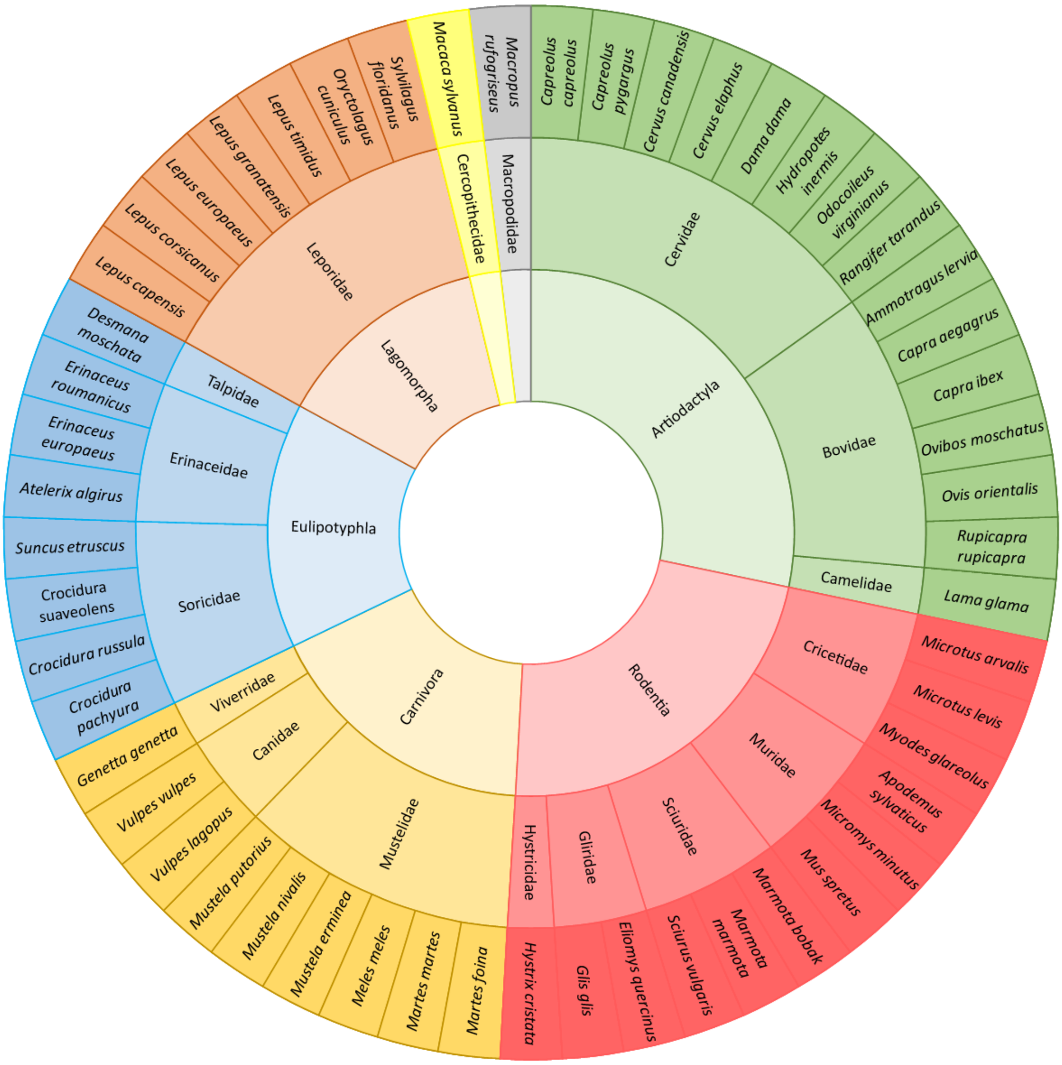


**Fig. S6.** Taxonomic assignment of all the introduced mammals established in Europe (n = 53). The inner circle represents the orders, the middle circle the families and the outer circle the species.

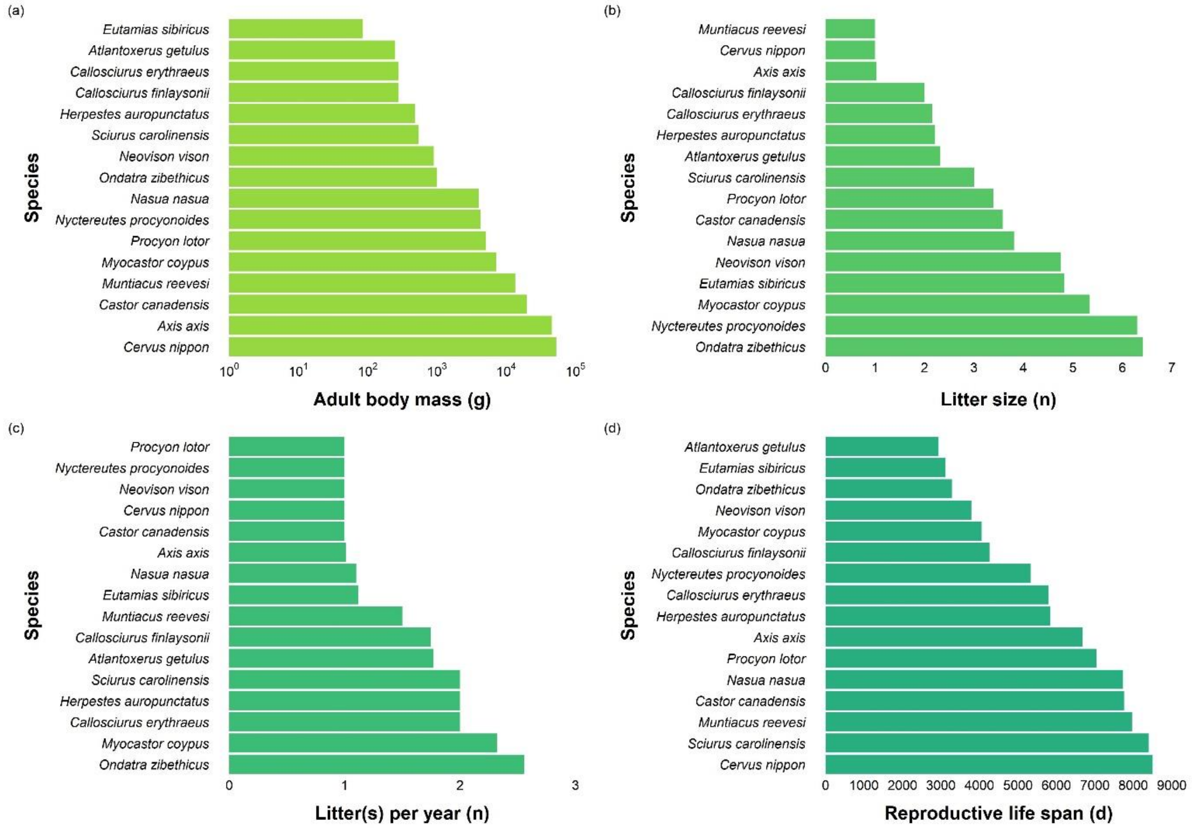


**Fig. S7.** Traits favouring the introduction, establishment, and spread (Capellini et al. 2015; Blackburn et al. 2017) of the study species: (a) adult body mass (log scale, in grams), (b) litter size, (c) litter(s) per year, and (d) generation length (in days).

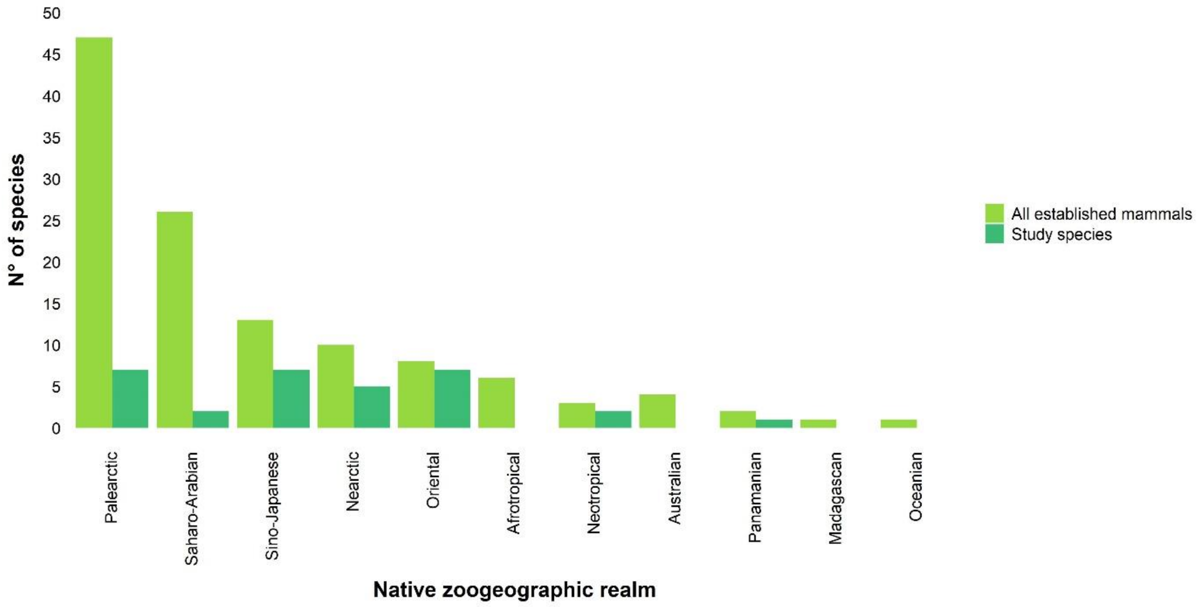


**Fig. S8.** Native zoogeographic realms (Holt et al., 2013) of all the introduced mammals established in Europe (n = 119) and of the study species (n = 31). Marginal parts of native ranges occurring in less than 1% of a zoogeographic realm were not considered.

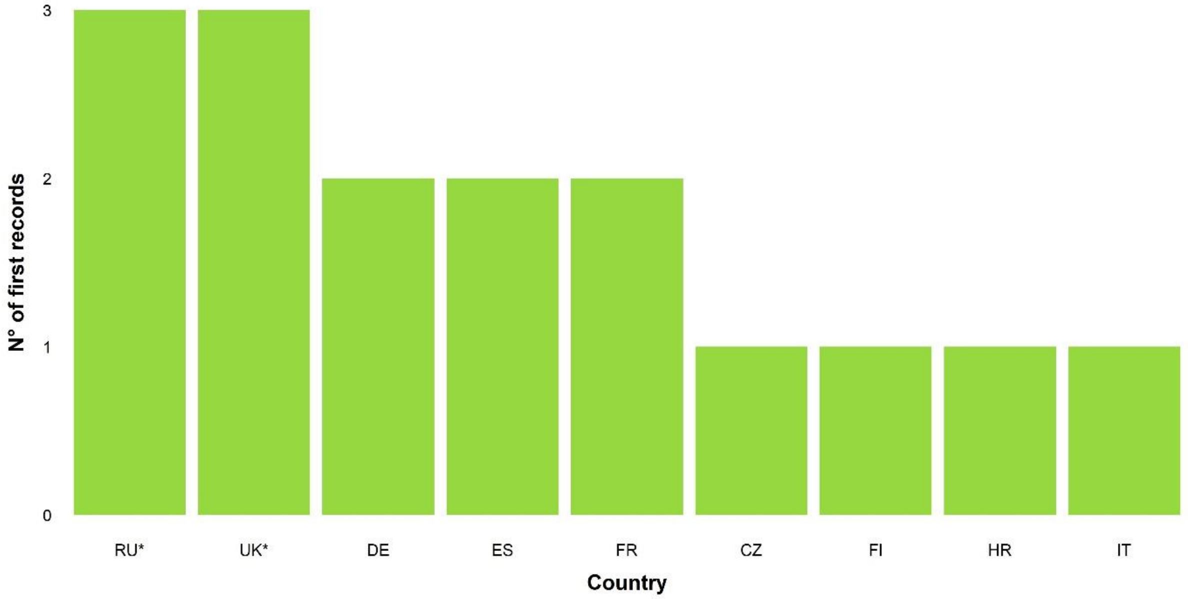


**Fig. S9.** First continental records in the countries of Europe (n = 16). Countries without invasive mammal species are not shown. Countries marked with an asterisk (*) are not Member States of the EU.

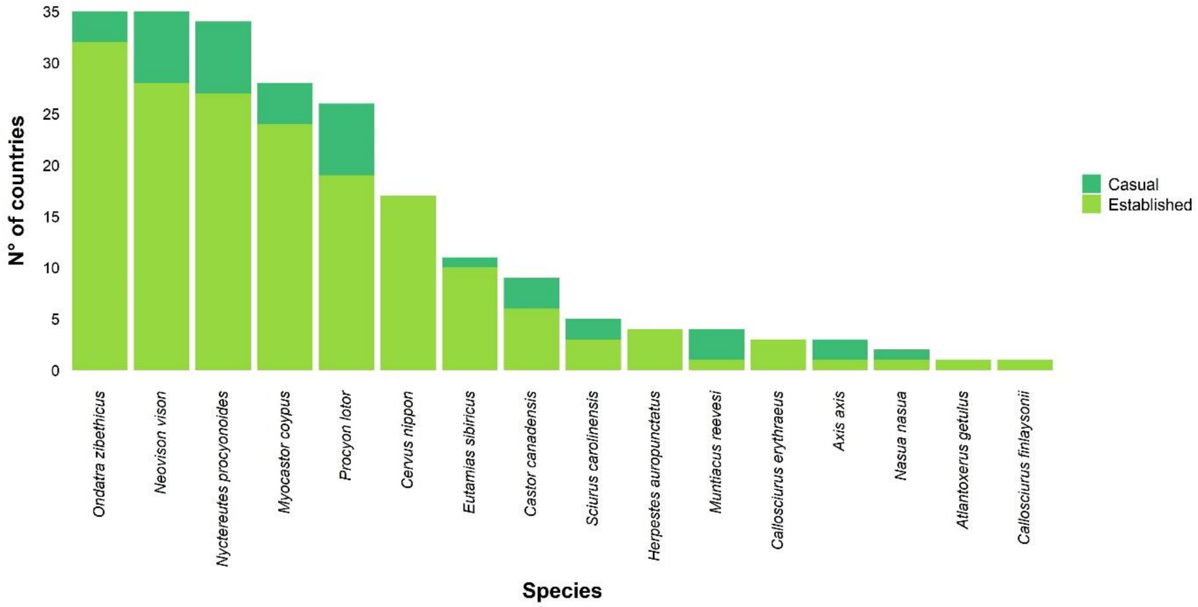


**Fig. S10.** Number of countries in Europe with established and casual presences (n = 218) of the study species.

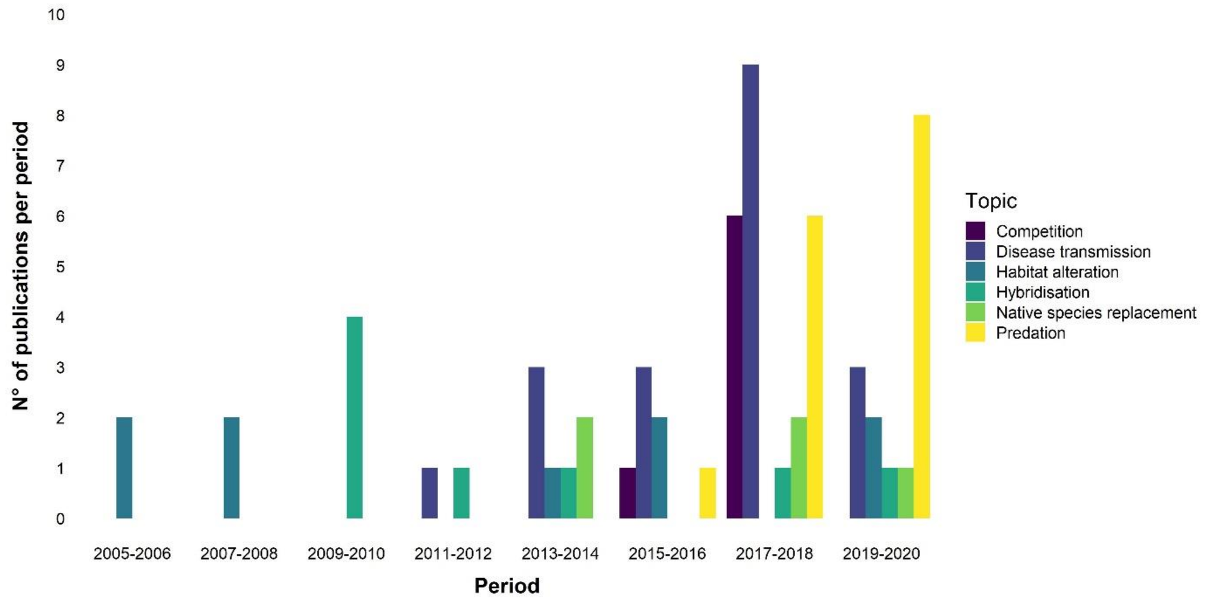


**Fig. S11.** The number of published papers (n = 63) regarding environmental impacts of invasive mammal species in Europe, divided per impact categories (following Blackburn et al., 2014).

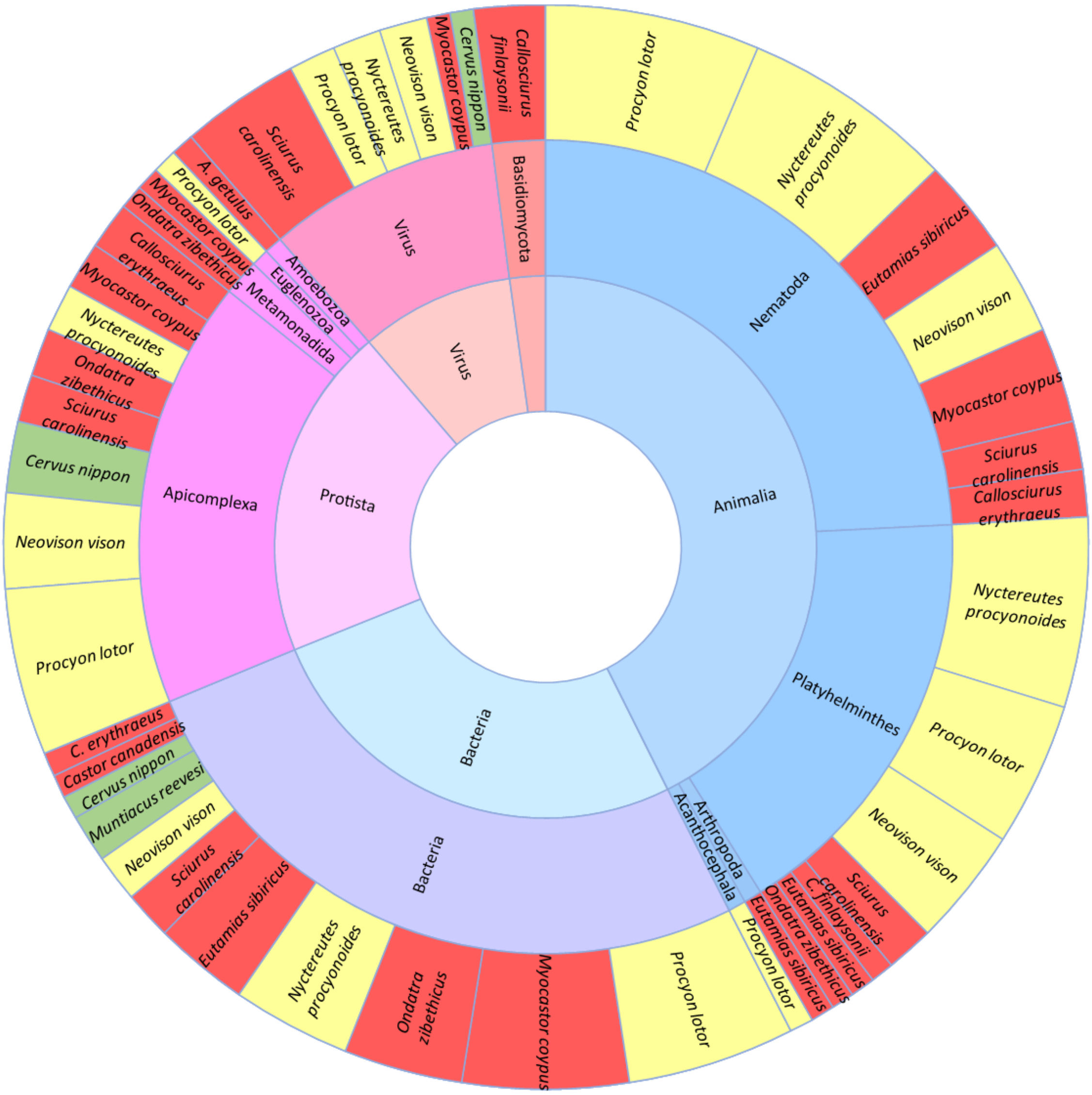


**Fig. S12.** Taxonomic assignments of the pathogens (n = 141) and study species (n = 13) known to be infected by them. The inner circle represents the Domain or the Kingdom, the middle circle the phyla, and the outer circle the species know to be infected. Species pertaining to the same order are indicated with the same colour (yellow for Carnivora, red for Rodentia, green for Artiodactyla).

**Appendix S3.** List of the papers obtained through the literature search process for each study species in Europe.

*Atlantoxerus getulus*


1. Gangoso, L., Donázar, J. A., Scholz, S., Palacios, C. J., & Hiraldo, F. (2006). Contradiction in conservation of island ecosystems: Plants, introduced herbivores and avian scavengers in the Canary Islands. Biodiversity and Conservation, 15(7), 2231–2248. https://doi.org/10.1007/s10531-004-7181-4
2. Nogales, M., Rodríguez-Luengo, J. L., & Marrero, P. (2006). Ecological effects and distribution of invasive non-native mammals on the Canary Islands. Mammal Review, 36(1), 49–65. https://doi.org/10.1111/j.1365-2907.2006.00077.x
3. Lorenzo-Morales, J., López-Darias, M., Martínez-Carretero, E., & Valladares, B. (2007). Isolation of potentially pathogenic strains of Acanthamoeba in wild squirrels from the Canary Islands and Morocco. Experimental Parasitology, 117(1), 74–79. https://doi.org/10.1016/j.exppara.2007.03.014
4. López-Darias, M., Lobo, J. M., & Gouat, P. (2008). Predicting potential distributions of invasive species: The exotic Barbary ground squirrel in the Canarian archipelago and the west Mediterranean region. Biological Invasions, 10(7), 1027–1040. https://doi.org/10.1007/s10530-007-9181-2
5. López-Darias, M., & Nogales, M. (2008). Effects of the invasive Barbary ground squirrel (*Atlantoxerus getulus*) on seed dispersal systems of insular xeric environments. Journal of Arid Environments, 72(6), 926–939. https://doi.org/10.1016/j.jaridenv.2007.12.006
6. Traveset, A., Nogales, M., Alcover, J. A., Delgado, J. D., López-Darias, M., Godoy, D., Igual, J. M., & Bover, P. (2009). A review on the effects of alien rodents in the Balearic (western Mediterranean sea) and Canary islands (eastern Atlantic ocean). Biological Invasions, 11(7), 1653–1670. https://doi.org/10.1007/s10530-008-9395-y
7. Nogales, M., Nieves, C., Illera, J. C., Padilla, D. P., & Traveset, A. (2014). Effect of native and alien vertebrate frugivores patterns of *Rubia fruticosa* viability and germination in the eastern Canary Islands (Rubiaceae). Functional Ecology, 19, 429–436. https://doi.org/10.1111/j.1365-2435.2005.00975.x
8. Di Febbraro, M., Martinoli, A., Russo, D., Preatoni, D., & Bertolino, S. (2016). Modelling the effects of climate change on the risk of invasion by alien squirrels. Hystrix, 27(1), 1–8. https://doi.org/10.4404/hystrix-27.1-11776


*Axis axis*


9. Centore, L., Ugarković, D., Scaravelli, D., Safner, T., Pandurić, K., & Sprem, N. (2018). Locomotor activity pattern of two recently introduced non-native ungulate species in a Mediterranean habitat. Folia Zoologica, 67(1), 17–24. https://doi.org/10.25225/fozo.v67.i1.a1.2018
10. Šprem, N., & Zachos, F. E. (2020). Axis Deer *Axis axis* Erxleben, 1777. In K. Hackländer & F. E. Zachos (Eds.), Handbook of the Mammals of Europe (pp. 1–9). Springer Nature Switzerland. https://doi.org/10.1007/978-3-319-65038-8_22-2


*Callosciurus erythraeus*


11. Tamura, N. (2009). Datasheet on *Callosciurus erythraeus*. Wallingford (UK): CAB International, Invasive Species Compendium. Available from: http://www.cabi.org/isc.
12. Bertolino, S., & Lurz, P. W. W. (2013). Callosciurus squirrels: Worldwide introductions, ecological impacts and recommendations to prevent the establishment of new invasive populations. Mammal Review, 43(1), 22–33. https://doi.org/10.1111/j.1365-2907.2011.00204.x
13. Mazzamuto, M. V., Wauters, L., Martinoli, A., & Bertolino, S. (2014). EU NON-NATIVE ORGANISM RISK ASSESSMENT SCHEME - *Callosciurus erythraeus*.
14. Adriaens, T., Baert, K., Breyne, P., Casaer, J., Devisscher, S., Onkelinx, T., Pieters, S., & Stuyck, J. (2015). Successful eradication of a suburban Pallas's squirrel *Callosciurus erythraeus* (Pallas 1779) (Rodentia, Sciuridae) population in Flanders (northern Belgium). Biological Invasions, 17(9), 2517–2526. https://doi.org/10.1007/s10530-015-0898-z

15. Dozières, A., Pisanu, B., Kamenova, S., Bastelica, F., Gerriet, O., & Chapuis, J. L. (2015). Range expansion of Pallas's squirrel (*Callosciurus erythraeus*) introduced in southern France: Habitat suitability and space use. Mammalian Biology, 80(6), 518–526. https://doi.org/10.1016/j.mambio.2015.08.004
16. Hofmannová, L., Romeo, C., Štohanzlová, L., Jirsová, D., Mazzamuto, M. V., Wauters, L. A., Ferrari, N., & Modrý, D. (2016). Diversity and host specificity of coccidia (Apicomplexa: Eimeriidae) in native and introduced squirrel species. European Journal of Protistology, 56, 1–14. https://doi.org/10.1016/j.ejop.2016.04.008
17. Mazzamuto, M. V., Pisanu, B., Romeo, C., Ferrari, N., Preatoni, D., Wauters, L. A., Chapuis, J. L., & Martinoli, A. (2016). Poor Parasite Community of an Invasive Alien Species: Macroparasites of Pallas's Squirrel in Italy. Annales Zoologici Fennici, 53(1–2), 103–112. https://doi.org/10.5735/086.053.0209
18. Mazzamuto, M. V., Morandini, M., Panzeri, M., Wauters, L. A., Preatoni, D. G., & Martinoli, A. (2017a). Space invaders: effects of invasive alien Pallas's squirrel on home range and body mass of native red squirrel. Biological Invasions, 19(6), 1863–1877. https://doi.org/10.1007/s10530-017-1396-2
19. Mazzamuto, M. V., Bisi, F., Wauters, L. A., Preatoni, D. G., & Martinoli, A. (2017b). Interspecific competition between alien Pallas's squirrels and Eurasian red squirrels reduces density of the native species. Biological Invasions, 19(2), 723–735. https://doi.org/10.1007/s10530-016-1310-3
20. Prediger, J., Horčičková, M., Hofmannová, L., Sak, B., Ferrari, N., Mazzamuto, M. V., Romeo, C., Wauters, L. A., McEvoy, J., & Kváč, M. (2017). Native and introduced squirrels in Italy host different Cryptosporidium spp. European Journal of Protistology, 61, 64–75. https://doi.org/10.1016/j.ejop.2017.09.007
21. Schilling, A. K., Avanzi, C., Ulrich, R. G., Busso, P., Pisanu, B., Ferrari, N., Romeo, C., Mazzamuto, M. V., McLuckie, J., Shuttleworth, C. M., Del-Pozo, J., Lurz, P. W. W., Escalante-Fuentes, W. G., Ocampo-Candiani, J., Vera-Cabrera, L., Stevenson, K., Chapuis, J. L., Meredith, A. L., & Cole, S. T. (2019). British red squirrels remain the only known wild rodent host for leprosy bacilli. Frontiers in Veterinary Science, 6(FEB), 6–11. https://doi.org/10.3389/fvets.2019.00008

*Callosciurus finlaysonii*

22. Lurz, P. (2014). Datasheet on *Callosciurus finlaysonii*. Wallingford (UK): CAB International, Invasive Species Compendium. Available from: http://www.cabi. org/isc.
23. Mori, E., Mazzoglio, P. J., Rima, P. C., Aloise, G., & Bertolino, S. (2016a). Bark-stripping damage by *Callosciurus finlaysonii* introduced into Italy. Mammalia, 80(5), 507–514. https://doi.org/10.1515/mammalia-2015-0107
24. Bertolino, S., Adriaens, T., Verzelen, Y., Rabitsch, W., Robertson, P., Kettunen, M., Chapman, D., & Scalera, R. (2018). Study on Invasive Alien Species – Development of risk assessments to tackle priority species and enhance prevention (*Callosciurus finlaysonii*).

*Castor canadensis*

25. Aldridge, V. (2009). Datasheet on *Castor canadensis*. Wallingford (UK): CAB International, Invasive Species Compendium. Available from: http://www.cabi. org/isc.
26. Nummi, P. (2010). NOBANIS - Invasive Alien Species Fact Sheet: *Castor canadensis*. https://doi.org/10.3732/ajb.1000402
27. Dewas, M., Herr, J., Schley, L., Angst, C., Manet, B., Landry, P., & Catusse, M. (2012). Recovery and status of native and introduced beavers *Castor fiber* and *Castor canadensis* in France and neighbouring countries. Mammal Review, 42(2), 144–165. https://doi.org/10.1111/j.1365-2907.2011.00196.x
28. Parker, H., Nummi, P., Hartman, G., & Rosell, F. (2012). Invasive North American beaver *Castor canadensis* in Eurasia: A review of potential consequences and a strategy for eradication. Wildlife Biology, 18(4), 354–365. https://doi.org/10.2981/12-007
29. Frosch, C., Kraus, R. H. S., Angst, C., Allgöwer, R., Michaux, J., Teubner, J., & Nowak, C. (2014). The genetic legacy of multiple beaver reintroductions in central Europe. PLoS ONE, 9(5). https://doi.org/10.1371/journal.pone.0097619

30. Holopainen, S., Nummi, P., & Pöysä, H. (2014). Breeding in the stable boreal landscape: Lake habitat variability drives brood production in the teal (Anas crecca). Freshwater Biology, 59(12), 2621–2631. https://doi.org/10.1111/fwb.12458
31. Nummi, P., & Holopainen, S. (2014). Whole-community facilitation by beaver: Ecosystem engineer increases waterbird diversity. Aquatic Conservation: Marine and Freshwater Ecosystems, 24(5), 623–633. https://doi.org/10.1002/aqc.2437
32. Sissonen, S., Rossow, H., Edvin, K., HemmilÄ, H., Henttonen, H., Isomursu, M., Kinnunen, P. M., Pelkola, K., Pelkonen, S., Tarkka, E., MyrtennÄs, K., Nikkari, S., & Forsman, M. (2015). Phylogeography of Francisella tularensis subspecies holarctica in Finland, 1993-2011. Infectious Diseases, 47(10), 701–706. https://doi.org/10.3109/23744235.2015.1049657
33. Vehkaoja, M., & Nummi, P. (2015). Beaver facilitation in the conservation of boreal anuran communities. Herpetozoa, 28(1/2), 75–87. https://doi.org/1013-4425
34. Whitfield, C. J., Baulch, H. M., Chun, K. P., & Westbrook, C. J. (2015). Beaver-mediated methane emission: The effects of population growth in Eurasia and the Americas. Ambio, 44(1), 7–15. https://doi.org/10.1007/s13280-014-0575-y
35. Hollander, H., Van Duinen, G. A., Branquart, E., De Hoop, L., De Hullu, P. C., Matthews, J., Van der Velde, G., & Leuven, R. S. E. W. (2017). Risk assessment of the alien North American beaver (*Castor canadensis*).
36. Nummi, P., Suontakanen, E. M., Holopainen, S., & Väänänen, V. M. (2019a). The effect of beaver facilitation on Common Teal: pairs and broods respond differently at the patch and landscape scales. Ibis, 161(2), 301–309. https://doi.org/10.1111/ibi.12626
37. Halley, D. J., Saveljev, A. P., & Rosell, F. (2020). Population and distribution of beavers *Castor fiber* and *Castor canadensis* in Eurasia. Mammal Review, 1–24. https://doi.org/10.1111/mam.12216

*Cervus nippon*

38. McDevitt, A. D., Edwards, C. J., O'Toole, P., O'Sullivan, P., O'Reilly, C., & Carden, R. F. (2009). Genetic structure of, and hybridisation between, red (*Cervus elaphus*) and sika (*Cervus nippon*) deer in Ireland. Mammalian Biology, 74(4), 263–273. https://doi.org/10.1016/j.mambio.2009.03.015
39. Putman, R. (2009a). Datasheet on *Cervus nippon*. Wallingford (UK): CAB International, Invasive Species Compendium. Available from: http://www.cabi. org/isc.
40. Robinson, M. T., Shaw, S. E., & Morgan, E. R. (2009). Anaplasma phagocytophilum infection in a multi-species deer community in the New Forest, England. European Journal of Wildlife Research, 55(4), 439–442. https://doi.org/10.1007/s10344-009-0261-8
41. Sedlak, K., Girma, T., & Holejsovsky, J. (2009). Pestivirus infections in cervids from the Czech Republic. Veterinarni Medicina, 54(4), 191–193. https://doi.org/10.17221/29/2009-VETMED
42. Senn, H. V., & Pemberton, J. M. (2009). Variable extent of hybridization between invasive sika (*Cervus nippon*) and native red deer (*C. elaphus*) in a small geographical area. Molecular Ecology, 18(5), 862–876. https://doi.org/10.1111/j.1365-294X.2008.04051.x
43. Acevedo, P., Ward, A. I., Real, R., & Smith, G. C. (2010). Assessing biogeographical relationships of ecologically related species using favourability functions: A case study on British deer. Diversity and Distributions, 16(4), 515–528. https://doi.org/10.1111/j.1472-4642.2010.00662.x
44. Radwan, J., Demiaszkiewicz, A. W., Kowalczyk, R., Lachowicz, J., Kawałko, A., Wójcik, J. M., Pyziel, A. M., & Babik, W. (2010). An evaluation of two potential risk factors, MHC diversity and host density, for infection by an invasive nematode Ashworthius sidemi in endangered European bison (Bison bonasus). Biological Conservation, 143(9), 2049–2053. https://doi.org/10.1016/j.biocon.2010.05.012
45. Senn, H. V., Barton, N. H., Goodman, S. J., Swanson, G. M., Abernethy, K. A., & Pemberton, J. M. (2010a). Investigating temporal changes in hybridization and introgression in a predominantly bimodal hybridizing population of invasive sika (*Cervus nippon*) and native red deer (*C. elaphus*) on the Kintyre Peninsula, Scotland. Molecular Ecology, 19(5), 910–924. https://doi.org/10.1111/j.1365-294X.2009.04497.x

46. Senn, H. V., Swanson, G. M., Goodman, S. J., Barton, N. H., & Pemberton, J. M. (2010b). Phenotypic correlates of hybridisation between red and sika deer (genus Cervus). Journal of Animal Ecology, 79(2), 414–425. https://doi.org/10.1111/j.1365-2656.2009.01633.x
47. GB Non-Native Species Secretariat. (2011). GB NON-NATIVE ORGANISM RISK ASSESSMENT SCHEME - *Cervus nippon*. https://circabc.europa.eu/sd/a/284ef858-4601-4def-969b-3276abc69a0c/*Cervus nippon* - GBNNRA.pdf
48. Carden, R. F., Carlin, C. M., Marnell, F., McElholm, D., Hetherington, J., & Gammell, M. P. (2011). Distribution and range expansion of deer in Ireland. Mammal Review, 41(4), 313–325. https://doi.org/10.1111/j.1365-2907.2010.00170.x
49. Perrin, P. M., Mitchell, F. J. G., & Kelly, D. L. (2011). Long-term deer exclusion in yew-wood and oakwood habitats in southwest Ireland: Changes in ground flora and species diversity. Forest Ecology and Management, 262(12), 2328–2337. https://doi.org/10.1016/j.foreco.2011.08.028
50. Zachos, F. E., & Hartl, G. B. (2011). Phylogeography, population genetics and conservation of the European red deer *Cervus elaphus*. Mammal Review, 41(2), 138–150. https://doi.org/10.1111/j.1365-2907.2010.00177.x
51. Biedrzycka, A., Solarz, W., & Okarma, H. (2012). Hybridization between native and introduced species of deer in Eastern Europe. Journal of Mammalogy, 93(5), 1331–1341. https://doi.org/10.1644/11-MAMM-A-022.1
52. Liu, Y., & Nieuwenhuis, M. (2014). An analysis of habitat-use patterns of fallow and sika deer based on culling data from two estates in Co. Wicklow. Irish Forestry, December, 27–49.
53. Macháček, Z., Dvořák, S., Ježek, M., & Zahradník, D. (2014). Impact of interspecific relations between native red deer (*Cervus elaphus*) and introduced sika deer (*Cervus nippon*) on their rutting season in the Doupovské hory Mts. Journal of Forest Science, 60(7), 272–280. https://doi.org/10.17221/47/2014-jfs
54. Smith, S. L., Carden, R. F., Coad, B., Birkitt, T., & Pemberton, J. M. (2014). A survey of the hybridisation status of Cervus deer species on the island of Ireland. Conservation Genetics, 15(4), 823–835. https://doi.org/10.1007/s10592-014-0582-3
55. Ambroz, R., Vacek, S., Vacek, Z., Král, J., & Štefančík, I. (2015). Current and simulated structure, growth parameters and regeneration of beech forests with different game management in the Lány Game Enclosure. Forestry Journal, 61(2), 78–88. https://doi.org/10.1515/forj-2015-0016
56. Kubankova, M., Kralik, P., Lamka, J., Zakovcik, V., Dolanský, M., & Vasickova, P. (2015). Prevalence of Hepatitis E Virus in Populations of Wild Animals in Comparison with Animals Bred in Game Enclosures. Food and Environmental Virology, 7(2), 159–163. https://doi.org/10.1007/s12560-015-9189-1
57. Larska, M., Krzysiak, M. K., Jabłoński, A., Kesik, J., Bednarski, M., & Rola, J. (2015). Hepatitis E Virus Antibody Prevalence in Wildlife in Poland. Zoonoses and Public Health, 62(2), 105–110. https://doi.org/10.1111/zph.12113
58. Lorencova, A., Lamka, J., & Slany, M. (2015). Toxoplasma gondii in wild ruminants bred in game preserves and farms with production destined for human consumption in the Czech Republic. Potravinarstvo, 9(1), 288–292. https://doi.org/10.5219/482
59. Dvořák, J., & Palyzová, L. (2016). Analysis of the development and spatial distribution of sika deer (*Cervus nippon*) populations on the territory of the Czech Republic. Acta Universitatis Agriculturae et Silviculturae Mendelianae Brunensis, 64(5), 1507–1515. https://doi.org/10.11118/actaun201664051507
60. Prakas, P., Butkauskas, D., Rudaitytė, E., Kutkienė, L., Sruoga, A., & Pūraitė, I. (2016). Morphological and molecular characterization of Sarcocystis taeniata and Sarcocystis pilosa n. sp. from the sika deer (*Cervus nippon*) in Lithuania. Parasitology Research, 115(8), 3021–3032. https://doi.org/10.1007/s00436-016-5057-7
61. Graham, D. A., Gallagher, C., Carden, R. F., Lozano, J. M., Moriarty, J., & O'Neill, R. (2017). A survey of free-ranging deer in Ireland for serological evidence of exposure to bovine viral diarrhoea virus, bovine herpes virus-1, bluetongue virus and Schmallenberg virus. Irish Veterinary Journal, 70(1), 1–11. https://doi.org/10.1186/s13620-017-0091-z
62. Kopij, G. (2017). Expansion of alien carnivore and ungulate species in SW Poland. Russian Journal of Biological Invasions, 8(3), 290–299. https://doi.org/10.1134/S2075111717030031

63. Panova, O. A., Serdyuk, N. V., Glamazdin, I. G., & Zemlyanko, I. I. (2017). Retrospective and prospective studies on helminthiases in bisons of Prioksko-Terrasny Nature Reserve (Moscow Region, Serpukhov District). Russian Journal of Theriology, 16(2), 149–156. https://doi.org/10.15298/rusjtheriol.16.2.04
64. Nechybová, S., Vejl, P., Hart, V., Melounová, M., Čílová, D., Vašek, J., Jankovská, I., Vadlejch, J., & Langrová, I. (2018a). Long-term occurrence of Trichuris species in wild ruminants in the Czech Republic. Parasitology Research, 117(6), 1699–1708. https://doi.org/10.1007/s00436-018-5841-7
65. Rudaitytė-Lukošienė, E., Prakas, P., Butkauskas, D., Kutkienė, L., Vepštaitė-Monstavičė, I., & Servienė, E. (2018). Morphological and molecular identification of Sarcocystis spp. from the sika deer (*Cervus nippon*), including two new species Sarcocystis frondea and Sarcocystis nipponi. Parasitology Research, 117(5), 1305–1315. https://doi.org/10.1007/s00436-018-5816-8
66. Smith, S. L., Senn, H. V., Pérez-Espona, S., Wyman, M. T., Heap, E., & Pemberton, J. M. (2018). Introgression of exotic Cervus (nippon and canadensis) into red deer (*Cervus elaphus*) populations in Scotland and the English Lake District. Ecology and Evolution, 8(4), 2122–2134. https://doi.org/10.1002/ece3.3767
67. Cukor, J., Vacek, Z., Linda, R., Vacek, S., Marada, P., Šimůnek, V., & Havránek, F. (2019). Effects of bark stripping on timber production and structure of Norway Spruce forests in relation to climatic factors. Forests, 10(4), 13–17. https://doi.org/10.3390/f10040320
68. Kurina, O., Kirik, H., Õunap, H., & Õunap, E. (2019). The northernmost record of a blood-sucking ectoparasite, Lipoptena fortisetosa Maa (Diptera: Hippoboscidae), in Estonia. Biodiversity Data Journal, 7. https://doi.org/10.3897/BDJ.7.E47857
69. Loy, A., Aloise, G., Ancillotto, L., Angelici, F. M., Bertolino, S., Capizzi, D., Castiglia, R., Colangelo, P., Contoli, L., Cozzi, B., Fontaneto, D., Lapini, L., Maio, N., Monaco, A., Mori, E., Nappi, A., Podestà, M., Russo, D., Sarà, M., … Amori, G. (2019). Mammals of Italy: an annotated checklist. Hystrix, Italian Journal of Mammalogy, 30(2), 87–106. https://doi.org/10.4404/hystrix
70. Hrazdilová, K., Rybářová, M., Široký, P., Votýpka, J., Zintl, A., Burgess, H., Steinbauer, V., Žákovčík, V., & Modrý, D. (2020). Diversity of Babesia spp. in cervid ungulates based on the 18S rDNA and cytochrome c oxidase subunit I phylogenies. Infection, Genetics and Evolution, 77(October 2019), 104060. https://doi.org/10.1016/j.meegid.2019.104060
71. McFarlane, S. E., Hunter, D. C., Senn, H. V., Smith, S. L., Holland, R., Huisman, J., & Pemberton, J. M. (2020). Increased genetic marker density reveals high levels of admixture between red deer and introduced Japanese sika in Kintyre, Scotland. Evolutionary Applications, 13(2), 432–441. https://doi.org/10.1111/eva.12880
72. Trojnar, E., Kästner, B., & Johne, R. (2020). No Evidence of Hepatitis E Virus Infection in Farmed Deer in Germany. Food and Environmental Virology, 12(1), 81–83. https://doi.org/10.1007/s12560-019-09407-y
73. Vacek, Z., Cukor, J., Linda, R., Vacek, S., Šimůnek, V., Brichta, J., Gallo, J., & Prokůpková, A. (2020). Bark stripping, the crucial factor affecting stem rot development and timber production of Norway spruce forests in Central Europe. Forest Ecology and Management, 474(April), 118360. https://doi.org/10.1016/j.foreco.2020.118360

*Eutamias sibiricus*

74. Pisanu, B., Jerusalem, C., Huchery, C., Marmet, J., & Chapuis, J. L. (2007). Helminth fauna of the Siberian chipmunk, *Tamias sibiricus* Laxmann (Rodentia, Sciuridae) introduced in suburban French forests. Parasitology Research, 100(6), 1375–1379. https://doi.org/10.1007/s00436-006-0389-3
75. Vourc'h, G., Marmet, J., Chassagne, M., Bord, S., & Chapuis, J. L. (2007). Borrelia burgdorferi sensu lato in Siberian chipmunks (*Tamias sibiricus*) introduced in suburban forests in France. Vector-Borne and Zoonotic Diseases, 7(4), 637–641. https://doi.org/10.1089/vbz.2007.0111
76. Chapuis, J.-L., Obolenskaya, E.V., Pisanu, B., Lissovsky, A.A. (2009). Datasheet on *Tamias sibiricus*. Wallingford (UK): CAB International, Invasive Species Compendium. Available from: http://www.cabi. org/isc.

77. Pisanu, B., Lebailleux, L., & Chapuis, J. L. (2009). Why do Siberian chipmunks *Tamias sibiricus* (Sciuridae) introduced in French forests acquired so few intestinal helminth species from native sympatric Murids? Parasitology Research, 104(3), 709–714. https://doi.org/10.1007/s00436-008-1279-7
78. Pisanu, B., Marsot, M., Marmet, J., Chapuis, J. L., Réale, D., & Vourc'h, G. (2010). Introduced Siberian chipmunks are more heavily infested by ixodid ticks than are native bank voles in a suburban forest in France. International Journal for Parasitology, 40(11), 1277–1283. https://doi.org/10.1016/j.ijpara.2010.03.012
79. Marsot, M., Sigaud, M., Chapuis, J. L., Ferquel, E., Cornet, M., & Vourc'h, G. (2011). Introduced Siberian chipmunks (*Tamias sibiricus* barberi) harbor more-diverse Borrelia burgdorferi sensu lato genospecies than native bank voles (Myodes glareolus). Applied and Environmental Microbiology, 77(16), 5716–5721. https://doi.org/10.1128/AEM.01846-10
80. Marsot, M., Chapuis, J. L., Gasqui, P., Dozières, A., Masséglia, S., Pisanu, B., Ferquel, E., & Vourc'h, G. (2013). Introduced Siberian Chipmunks (*Tamias sibiricus* barberi) Contribute More to Lyme Borreliosis Risk than Native Reservoir Rodents. PLoS ONE, 8(1), 1–8. https://doi.org/10.1371/journal.pone.0055377
81. Bonnet, S., Choumet, V., Masseglia, S., Cote, M., Ferquel, E., Lilin, T., Marsot, M., Chapuis, J. L., & Vourc'h, G. (2015). Infection of Siberian chipmunks (*Tamias sibiricus* barberi) with Borrelia sp. reveals a low reservoir competence under experimental conditions. Ticks and Tick-Borne Diseases, 6(3), 393–400. https://doi.org/10.1016/j.ttbdis.2015.03.008
82. d'Ovidio, D., Noviello, E., Pepe, P., Del Prete, L., Cringoli, G., & Rinaldi, L. (2015). Survey of Hymenolepis spp. in pet rodents in Italy. Parasitology Research, 114(12), 4381–4384. https://doi.org/10.1007/s00436-015-4675-9
83. Vourc'h, G., Abrial, D., Bord, S., Jacquot, M., Masséglia, S., Poux, V., Pisanu, B., Bailly, X., & Chapuis, J. L. (2016). Mapping human risk of infection with Borrelia burgdorferi sensu lato, the agent of Lyme borreliosis, in a periurban forest in France. Ticks and Tick-Borne Diseases, 7(5), 644–652. https://doi.org/10.1016/j.ttbdis.2016.02.008
84. Mori, E., Milanesi, P., Menchetti, M., Zozzoli, R., Monaco, A., Capizzi, D., & Nerva, L. (2018a). Genetics reveals that free-ranging chipmunks introduced to Italy have multiple origins. Hystrix, Italian Journal of Mammalogy, 29(December), 81–85. https://doi.org/10.4404/hystrix
85. Mori, E., Pisanu, B., Zozzoli, R., Solano, E., Olivieri, E., Sassera, D., & Montagna, M. (2018b). Arthropods and associated pathogens from native and introduced rodents in Northeastern Italy. Parasitology Research, 117(10), 3237–3243. https://doi.org/10.1007/s00436-018-6022-4
86. Mori, E., Zozzoli, R., & Mazza, G. (2018c). Coming in like a wrecking-ball: are native Eurasian red squirrels displacing invasive Siberian chipmunks? A study from an urban park. Urban Ecosystems, 21(5), 975–981. https://doi.org/10.1007/s11252-018-0775-5
87. Mori, E., Zozzoli, R., & Menchetti, M. (2018d). Global distribution and status of introduced Siberian chipmunks *EuTamias sibiricus*. Mammal Review, 48(2), 139–152. https://doi.org/10.1111/mam.12117
88. Di Febbraro, M., Menchetti, M., Russo, D., Ancillotto, L., Aloise, G., Roscioni, F., Preatoni, D. G., Loy, A., Martinoli, A., Bertolino, S., & Mori, E. (2019). Integrating climate and land-use change scenarios in modelling the future spread of invasive squirrels in Italy. Diversity and Distributions, 25(4), 644–659. https://doi.org/10.1111/ddi.12890
89. Andreoni, A., Augugliaro, C., Zozzoli, R., Dartora, F., & Mori, E. (2020). Diel activity patterns and overlap between Eurasian red squirrels and Siberian chipmunks in native and introduced ranges. Ethology Ecology and Evolution, 00(00), 1–7. https://doi.org/10.1080/03949370.2020.1777211

*Herpestes auropunctatus*

90. Deputy Direction of Nature. (2015). EU NON-NATIVE RISK ASSESSMENT SCHEME - *Herpestes javanicus*. 45.
91. Gaubert, P. (2015). Fate of the Mongooses and the Genet (Carnivora) in Mediterranean Europe: None Native, All Invasive? In F. M. Angelici (Ed.), Problematic Wildlife: A Cross-Disciplinary Approach (pp. 295–314). Springer International Publishing. https://doi.org/10.1007/978-3-319-22246-2

92. Müller, T., Freuling, C. M., Wysocki, P., Roumiantzeff, M., Freney, J., Mettenleiter, T. C., & Vos, A. (2015). Terrestrial rabies control in the European Union: Historical achievements and challenges ahead. Veterinary Journal, 203(1), 10–17. https://doi.org/10.1016/j.tvjl.2014.10.026

*Muntiacus reevesi*

93. Genovesi, P., Josefsson, M., Booy, O., Scalera, R., & Gallardo, B. (n.d.). GB NNRA - *Muntiacus reevesi*. https://doi.org/10.1016/j.cub.2005.09.007
94. Putman, R. (2009b). Datasheet on *Muntiacus reevesi*. Wallingford (UK): CAB International, Invasive Species Compendium. Available from: http://www.cabi. org/isc.
95. GB Non-Native Species Secretariat. (2011). GB NON-NATIVE ORGANISM RISK ASSESSMENT SCHEME - *Muntiacus reevesi*. https://circabc.europa.eu/sd/a/ad4e3149-017d-4204-b5c7-d3711b36cb83/*Muntiacus reevesi*i - GBNNRA.pdf
96. Putman, R., Langbein, J., Green, P., & Watson, P. (2011). Identifying threshold densities for wild deer in the UK above which negative impacts may occur. Mammal Review, 41(3), 175–196. https://doi.org/10.1111/j.1365-2907.2010.00173.x
97. Newson, S. E., Johnston, A., Renwick, A. R., Baillie, S. R., & Fuller, R. J. (2012). Modelling large-scale relationships between changes in woodland deer and bird populations. Journal of Applied Ecology, 49(1), 278–286. https://doi.org/10.1111/j.1365-2664.2011.02077.x
98. Ward, A. I., & Smith, G. C. (2012). Predicting the status of wild deer as hosts of Mycobacterium bovis infection in Britain. European Journal of Wildlife Research, 58(1), 127–135. https://doi.org/10.1007/s10344-011-0553-7
99. Baiwy, E., Schockert, V., & Branquart, E. (2013). Risk analysis of the Reeves' muntjac *Muntiacus reevesi*. 37.
100. O'Flynn, C., Kelly, J., & O'Rourke, E. (2014). Risk Assessment of *Muntiacus reevesi*. 24.
101. Freeman, M. S., Beatty, G. E., Dick, J. T. A., Reid, N., & Provan, J. (2016). The paradox of invasion: Reeves' muntjac deer invade the British Isles from a limited number of founding females. Journal of Zoology, 298(1), 54–63. https://doi.org/10.1111/jzo.12283
102. McKillen, J., Hogg, K., Lagan, P., Ball, C., Doherty, S., Reid, N., Collins, L., & Dick, J. T. A. (2017). Detection of a novel gammaherpesvirus (genus Rhadinovirus) in wild muntjac deer in Northern Ireland. Archives of Virology, 162(6), 1737–1740. https://doi.org/10.1007/s00705-017-3254-z
103. Croft, S., Ward, A. I., Aegerter, J. N., & Smith, G. C. (2019). Modeling current and potential distributions of mammal species using presence-only data: A case study on British deer. Ecology and Evolution, 9(15), 8724–8735. https://doi.org/10.1002/ece3.5424
104. Duscher, G. G., Battisti, E., Hodžić, A., Wäber, K., Steinbach, P., Stubbe, M., & Heddergott, M. (2020). First detection and molecular identification of Anaplasma phagocytophilum in an introduced population of Reeve's muntjac (*Muntiacus reevesi*) in United Kingdom. Molecular and Cellular Probes, 52(April), 101582. https://doi.org/10.1016/j.mcp.2020.101582

*Myocastor coypus*

105. Bertolino, S. (2008). Datasheet on *Myocastor coypus*. Wallingford (UK): CAB International, Invasive Species Compendium. Available from: http://www.cabi.org/isc.
106. Bertolino, S. (2014). GB NON-NATIVE ORGANISM RISK ASSESSMENT SCHEME - *Myocastor coypus*. 9. http://www.nonnativespecies.org/downloadDocument.cfm?id=55
107. Vein, J., Leblond, A., Belli, P., Kodjo, A., & Berny, P. J. (2014). The role of the coypu (*Myocastor coypus*), an invasive aquatic rodent species, in the epidemiological cycle of leptospirosis: A study in two wetlands in the East of France. European Journal of Wildlife Research, 60(1), 125–133. https://doi.org/10.1007/s10344-013-0758-z
108. Bertolino, S., Colangelo, P., Mori, E., & Capizzi, D. (2015). Good for management, not for conservation: An overview of research, conservation and management of Italian small mammals. Hystrix, 26(1), 1–11. https://doi.org/10.4404/hystrix-26.1-10263

109. Fratini, F., Turchi, B., Ebani, V. V., Bertelloni, F., Galiero, A., & Cerri, D. (2015). The presence of Leptospira in coypus (*Myocastor coypus*) and rats (Rattus norvegicus) living in a protected wetland in Tuscany (Italy). Veterinarski Arhiv, 85(4), 407–414.
110. Rylková, K., Tůmová, E., Brožová, A., Jankovská, I., Vadlejch, J., Čadková, Z., Frýdlová, J., Peřinková, P., Langrová, I., Chodová, D., Nechybová, S., & Schánková. (2015). Genetic and morphological characterization of Trichuris myocastoris found in *Myocastor coypus* in the Czech Republic. Parasitology Research, 114(11), 3969–3975. https://doi.org/10.1007/s00436-015-4623-8
111. Serracca, L., Battistini, R., Rossini, I., Mignone, W., Peletto, S., Boin, C., Pistone, G., Ercolini, R., & Ercolini, C. (2015). Molecular Investigation on the Presence of Hepatitis E Virus (HEV) in Wild Game in North-Western Italy. Food and Environmental Virology, 7(3), 206–212. https://doi.org/10.1007/s12560-015-9201-9
112. Adamopoulou, C., & Legakis, A. (2016). First account on the occurrence of selected invasive alien vertebrates in Greece. BioInvasions Records, 5(4), 189–196. https://doi.org/10.3391/bir.2016.5.4.01
113. Schulze, C., Heuner, K., Myrtennäs, K., Karlsson, E., Jacob, D., Kutzer, P., Große, K., Forsman, M., & Grunow, R. (2016). High and novel genetic diversity of Francisella tularensis in Germany and indication of environmental persistence. Epidemiology and Infection, 144(14), 3025–3036. https://doi.org/10.1017/S0950268816001175
114. Zanzani, S. A., Di Cerbo, A., Gazzonis, A. L., Epis, S., Invernizzi, A., Tagliabue, S., & Manfredi, M. T. (2016). Parasitic and bacterial infections of *Myocastor coypus* in a metropolitan area of northwestern Italy. Journal of Wildlife Diseases, 52(1), 126–130. https://doi.org/10.7589/2015-01-010
115. Gruychev, G. (2017). Distribution and density of coypu (*Myocastor coypus* (Molina, 1782)) in downstream of Maritsa River Southeast Bulgaria. Forestry Ideas, 23(1), 77–81.
116. Kellnerová, K., Holubová, N., Jandová, A., Vejčík, A., McEvoy, J., Sak, B., & Kváč, M. (2017). First description of Cryptosporidium ubiquitum XIIa subtype family in farmed fur animals. European Journal of Protistology, 59, 108–113. https://doi.org/10.1016/j.ejop.2017.03.007
117. Sicuro, B., Valle, E., Costa, P., Mussa, P., & Tarantola, M. (2017). The relation between exotic mammals and birds and agriculture productions in Italy: Modern containment strategies. Bulgarian Journal of Agricultural Science, 23(2), 242–251.
118. Nechybová, S., Langrová, I., & Tůmová, E. (2018b). Parasites of *Myocastor coypus* - A comparison in farm animals and their feral counterparts. Scientia Agriculturae Bohemica, 49(1), 21–25. https://doi.org/10.2478/sab-2018-0004
119. Bertelloni, F., Cilia, G., Turchi, B., Pinzauti, P., Cerri, D., & Fratini, F. (2019). Epidemiology of leptospirosis in North-Central Italy: Fifteen years of serological data (2002–2016). Comparative Immunology, Microbiology and Infectious Diseases, 65(January), 14–22. https://doi.org/10.1016/j.cimid.2019.04.001
120. Ayral, F., Kodjo, A., Guédon, G., Boué, F., & Richomme, C. (2020). Muskrats are greater carriers of pathogenic Leptospira than coypus in ecosystems with temperate climates. PLoS ONE, 15(2), 1–8. https://doi.org/10.1371/journal.pone.0228577
121. Gethöffer, F., & Siebert, U. (2020). Current knowledge of the Neozoa Nutria and Muskrat in Europe and their environmental impacts. Journal of Wildlife and Biodiversity, 4(2), 1–12. https://doi.org/10.22120/JWB.2019.109875.1074
122. Schertler, A., Rabitsch, W., Moser, D., Wessely, J., & Essl, F. (2020). The potential current distribution of the coypu (*Myocastor coypus*) in Europe and climate change induced shifts in the near future. NeoBiota, 58, 129–160. https://doi.org/10.3897/neobiota.58.33118

*Nasua nasua*

123. Deputy Direction of Nature. (2015). EU NON-NATIVE RISK ASSESSMENT SCHEME - *Nasua nasua*. 27.

*Neovison vison*

124. Bouros, G., Dekker, J., Gómez, A., Harrington, L. A., Hegyeli, Z., Hodor, C., Kauhala, K., Kranz, A., Korpimäki, E., Haye, M. La, Lambin, X., Macdonald, D., Mañas, S., Maran, T., Michaux, J. R., Moreno, L.,

Palazón, S., Põdra, M., Salo, P., … Zuberogoitia, I. (2016). EU NON-NATIVE ORGANISM RISK ASSESSMENT SCHEME - *Neovison vison*. 60.

125. Palazón, S. (2014). Datasheet on *Neovison vison*. Wallingford (UK): CAB International, Invasive Species Compendium. Available from: http://www.cabi. org/isc.

126. Barros, Á., Romero, R., Munilla, I., Pérez, C., & Velando, A. (2016). Behavioural plasticity in nest-site selection of a colonial seabird in response to an invasive carnivore. Biological Invasions, 18(11), 3149–3161. https://doi.org/10.1007/s10530-016-1205-3

127. Bouros, G., Dekker, J., Gómez, A., Harrington, L. A., Hegyeli, Z., Hodor, C., Kauhala, K., Kranz, A., Korpimäki, E., Haye, M. La, Lambin, X., Macdonald, D., Mañas, S., Maran, T., Michaux, J. R., Moreno, L., Palazón, S., Põdra, M., Salo, P., … Zuberogoitia, I. (2016). EU NON-NATIVE ORGANISM RISK ASSESSMENT SCHEME - *Neovison vison*.

128. Heddergott, M., Pohl, D., Steinbach, P., Salazar, L. C., Müller, F., & Frantz, A. C. (2016). Determinants and effects of sinus worm Skrjabingylus nasicola (Nematoda: Metastrongyloidae) infestation in invasive American mink *Neovison vison* in Germany. Parasitology Research, 115(9), 3449–3457. https://doi.org/10.1007/s00436-016-5107-1

129. Hurníková, Z., Kołodziej-Sobocińska, M., Dvorožňáková, E., Niemczynowicz, A., & Zalewski, A. (2016). An invasive species as an additional parasite reservoir: Trichinella in introduced American mink (*Neovison vison*). Veterinary Parasitology, 231, 106–109. https://doi.org/10.1016/j.vetpar.2016.06.010

130. Iordan, F., Lapini, L., Pavanello, M., Polednik, L., & Rieppi, C. (2016). Evidence for naturalization of the American mink (*Neovison vison*) in Friuli Venezia Giulia, NE Italy. Mammalia, 81(1), 91–94. https://doi.org/10.1515/mammalia-2015-0044

131. Manikowska-Ślepowrońska, B., Szydzik, B., & Jakubas, D. (2016). Determinants of the presence of conflict bird and mammal species at pond fisheries in western Poland. Aquatic Ecology, 50(1), 87–95. https://doi.org/10.1007/s10452-015-9554-z

132. Gholipour, H., Busquets, N., Fernández-Aguilar, X., Sánchez, A., Ribas, M. P., De Pedro, G., Lizarraga, P., Alarcia-Alejos, O., Temiño, C., & Cabezón, O. (2017). Influenza A Virus Surveillance in the Invasive American Mink (*Neovison vison*) from Freshwater Ecosystems, Northern Spain. Zoonoses and Public Health, 64(5), 363–369. https://doi.org/10.1111/zph.12316

133. Martínez-Rondán, F. J., Ruiz de Ybáñez, M. R., Tizzani, P., López-Beceiro, A. M., Fidalgo, L. E., & Martínez-Carrasco, C. (2017). The American mink (*Neovison vison*) is a competent host for native European parasites. Veterinary Parasitology, 247(October), 93–99. https://doi.org/10.1016/j.vetpar.2017.10.004

134. Miranda, C., Santos, N., Parrish, C., & Thompson, G. (2017). Genetic characterization of canine parvovirus in sympatric free-ranging wild carnivores in Portugal. Journal of Wildlife Diseases, 53(4), 824–831. https://doi.org/10.7589/2016-08-194

135. Niemczynowicz, A., Świętochowski, P., Brzeziński, M., & Zalewski, A. (2017). Non-native predator control increases the nesting success of birds: American mink preying on wader nests. Biological Conservation, 212(May), 86–95. https://doi.org/10.1016/j.biocon.2017.05.032

136. Nugaraite, D., Mazeika, V., & Paulauskas, A. (2017). Molecular and morphological characterization of Isthmiophora melis (Schrank, 1788) Luhe, 1909 (Digenea: Echinostomatidae) from American mink (*Neovison vison*) and European polecat (Mustela putorius) in Lithuania. Helminthologia (Poland), 54(2), 97–104. https://doi.org/10.1515/helm-2017-0012

137. Brzeziński, M., Ignatiuk, P., Żmihorski, M., & Zalewski, A. (2018a). An invasive predator affects habitat use by native prey: American mink and water vole co-existence in riparian habitats. Journal of Zoology, 304(2), 109–116. https://doi.org/10.1111/jzo.12500

138. Brzeziński, M., Chibowski, P., Gornia, J., Górecki, G., & Zalewski, A. (2018b). Spatio-temporal variation in nesting success of colonial waterbirds under the impact of a non-native invasive predator. Oecologia, 188(4), 1037–1047. https://doi.org/10.1007/s00442-018-4270-8

139. Criado-Fornelio, A., Martín-Pérez, T., Verdú-Expósito, C., Reinoso-Ortiz, S. A., & Pérez-Serrano, J. (2018). Molecular epidemiology of parasitic protozoa and Ehrlichia canis in wildlife in Madrid (central Spain). Parasitology Research, 117(7), 2291–2298. https://doi.org/10.1007/s00436-018-5919-2

140. Nugaraitė, D., Mažeika, V., & Paulauskas, A. (2018). Helminths of mustelids with overlapping ecological niches: Eurasian otter Lutra lutra (Linnaeus, 1758), American mink *Neovison vison* Schreber, 1777, and

European polecat Mustela putorius Linnaeus, 1758. Helminthologia, 56(1), 66–74. https://doi.org/10.2478/helm-2018-0035
141. Petersen, H. H., Nielsen, S. T., Larsen, G., Holm, E., & Chriél, M. (2018b). Prevalence of Capillaria plica in Danish wild carnivores. International Journal for Parasitology: Parasites and Wildlife, 7(3), 360–363. https://doi.org/10.1016/j.ijppaw.2018.09.006
142. Põdra, M., & Gómez, A. (2018). Rapid expansion of the American mink poses a serious threat to the European mink in Spain. Mammalia, 82(6), 580–588. https://doi.org/10.1515/mammalia-2017-0013
143. Prakas, P., Strazdaitė-Žielienė, Ž., Rudaitytė-Lukošienė, E., Servienė, E., & Butkauskas, D. (2018). Molecular identification of Sarcocystis lutrae (Apicomplexa: Sarcocystidae) in muscles of five species of the family Mustelidae. Parasitology Research, 117(6), 1989–1993. https://doi.org/10.1007/s00436-018-5880-0
144. Ribas, M. P., Almería, S., Fernández-Aguilar, X., De Pedro, G., Lizarraga, P., Alarcia-Alejos, O., Molina-López, R., Obón, E., Gholipour, H., Temiño, C., Dubey, J. P., & Cabezón, O. (2018). Tracking Toxoplasma gondii in freshwater ecosystems: interaction with the invasive American mink (*Neovison vison*) in Spain. Parasitology Research, 117(7), 2275–2281. https://doi.org/10.1007/s00436-018-5916-5
145. Roos, S., Smart, J., Gibbons, D. W., & Wilson, J. D. (2018). A review of predation as a limiting factor for bird populations in mesopredator-rich landscapes: a case study of the UK. Biological Reviews, 93(4), 1915–1937. https://doi.org/10.1111/brv.12426
146. Brzeziński, M., Pyrlik, J., Churski, M., Komar, E., & Zalewski, A. (2019a). The influence of American mink odour on the spatial distribution and behaviour of water voles. Ethology, 125(11), 791–801. https://doi.org/10.1111/eth.12933
147. Brzeziński, M., Żmihorski, M., Zarzycka, A., & Zalewski, A. (2019b). Expansion and population dynamics of a non-native invasive species: the 40-year history of American mink colonisation of Poland. Biological Invasions, 21(2), 531–545. https://doi.org/10.1007/s10530-018-1844-7
148. Koshev, Y. S. (2019). Occurrence of the American Mink *Neovison vison* (Schreber, 1777) (Carnivora: Mustelidae) in Bulgaria. Acta Zoologica Bulgarica, 71(3), 417–425.
149. Mori, E., & Mazza, G. (2019). Diet of a semiaquatic invasive mammal in northern Italy: Could it be an alarming threat to the endemic water vole? Mammalian Biology, 97, 88–94. https://doi.org/10.1016/j.mambio.2019.05.003
150. Sroka, J., Karamon, J., Wójcik-Fatla, A., Dutkiewicz, J., Bilska-Zając, E., Zając, V., Piotrowska, W., & Cencek, T. (2019). Toxoplasma gondii infection in selected species of free-living animals in Poland. Annals of Agricultural and Environmental Medicine, 26(4), 656–660. https://doi.org/10.26444/aaem/114930
151. Brzeziński, M., Żmihorski, M., Nieoczym, M., Wilniewczyc, P., & Zalewski, A. (2020). The expansion wave of an invasive predator leaves declining waterbird populations behind. Diversity and Distributions, 26(1), 138–150. https://doi.org/10.1111/ddi.13003
152. Flávio, H., Caballero, P., Jepsen, N., & Aarestrup, K. (2020). Atlantic salmon living on the edge: Smolt behaviour and survival during seaward migration in River Minho. Ecology of Freshwater Fish, April, 1–12. https://doi.org/10.1111/eff.12564
153. Garcìa, K., Sanpera, C., Lluìs, J., Palazón, S., Gosàlbez, J., Gòrski, K., & Melero, Y. (2020). High Trophic Niche Overlap between a Native and Invasive Mink Does Not Drive Trophic Displacement of the Native Mink during an Invasion Process. Animals, 10(1387). https://doi.org/10.3390/ani10081387
154. Hansen, J. E., Stegger, M., Pedersen, K., Sieber, R. N., Larsen, J., Larsen, G., Lilje, B., Chriél, M., Andersen, P. S., & Larsen, A. R. (2020). Spread of LA-MRSA CC398 in Danish mink (*Neovison vison*) and mink farm workers. Veterinary Microbiology, 245(October 2019), 108705. https://doi.org/10.1016/j.vetmic.2020.108705
155. Harrington, L. A., Birks, J., Chanin, P., & Tansley, D. (2020). Current status of American mink *Neovison vison* in Great Britain: a review of the evidence for a population decline. Mammal Review, 50(2), 157–169. https://doi.org/10.1111/mam.12184
156. Kołodziej-Sobocińska, M., Dvorožňáková, E., Hurníková, Z., Reiterová, K., & Zalewski, A. (2020). Seroprevalence of Echinococcus spp. and Toxocara spp. in Invasive Non-native American Mink. EcoHealth, 17(1), 13–27. https://doi.org/10.1007/s10393-020-01470-3

157. Lemming, L., Jørgensen, A. C., Nielsen, L. B., Nielsen, S. T., Mejer, H., Chriél, M., & Petersen, H. H. (2020). Cardiopulmonary nematodes of wild carnivores from Denmark: Do they serve as reservoir hosts for infections in domestic animals? International Journal for Parasitology: Parasites and Wildlife, 13(August), 90–97. https://doi.org/10.1016/j.ijppaw.2020.08.001
158. Molenaar, R. J., Vreman, S., Hakze-van der Honing, R. W., Zwart, R., de Rond, J., Weesendorp, E., Smit, L. A. M., Koopmans, M., Bouwstra, R., Stegeman, A., & van der Poel, W. H. M. (2020). Clinical and Pathological Findings in SARS-CoV-2 Disease Outbreaks in Farmed Mink (*Neovison vison*). Veterinary Pathology, 57(5), 653–657. https://doi.org/10.1177/0300985820943535
159. Petersen, H. H., Yang, R., Chriel, M., Liu, D., Hansen, M. S., & Ryan, U. M. (2020). Morphological and molecular characterization of Cystoisospora laidlawi oocysts (Apicomplexa: Eimeriidae) in farmed American mink (*Neovison vison*) in Denmark. Parasitology Research. https://doi.org/10.1007/s00436-020-06846-6
160. Oreshkova, N., Moelnaar, R. J., Vreman, S., Harders, F., Munnink, B. B. O., Van Der Honin, R. W. H., Gerhards, N., Tolsma, P., Bouwstra, R., Sikkema, R. S., Tacken, M. G. J., Rooij, M. M. T. De, Weesendorp, E., Engelsma, M. Y., Bruschke, C. J., Smit, L. A., Koopman, M., Van der Poel, W. H., & Stegeman, A. (2020). SARS-CoV-2 infection in farmed minks, the Netherlands, April and May 2020. Euro Surveillance, 25 (23)(May), 1–7. https://doi.org/10.2807/1560-7917.ES.2020.25.23.2001005

*Nyctereutes procyonoides*

161. Kauhala, K. (2009) Datasheet on *Nyctereutes procyonoides*. Wallingford (UK): CAB International, Invasive Species Compendium. Available from: http://www.cabi. org/isc.
162. Kowalczyk, R. (2014). NOBANIS - Invasive Alien Species Fact Sheet - *Nyctereutes procyonoides*. Online Database of the European Network on Invasive Alien Species - NOBANIS, Lv, 1–10.
163. Deputy Direction of Nature. (2016). EU NON-NATIVE ORGANISM RISK ASSESSMENT SCHEME - *Nyctereutes procyonoides*. 52.
164. Bagrade, G., Deksne, G., Ozoliņa, Z., Howlett, S. J., Interisano, M., Casulli, A., & Pozio, E. (2016). Echinococcus multilocularis in foxes and raccoon dogs: an increasing concern for Baltic countries. Parasites and Vectors, 9(1), 1–9. https://doi.org/10.1186/s13071-016-1891-9
165. Drygala, F., Korablev, N., Ansorge, H., Fickel, J., Isomursu, M., Elmeros, M., Kowalczyk, R., Baltrunaite, L., Balciauskas, L., Saarma, U., Schulze, C., Borkenhagen, P., & Frantz, A. C. (2016). Homogenous population genetic structure of the non-native raccoon dog (*Nyctereutes procyonoides*) in Europe as a result of rapid population expansion. PLoS ONE, 11(4), 1–17. https://doi.org/10.1371/journal.pone.0153098
166. Griciuviene, L., Paulauskas, A., Radzijevskaja, J., Žukauskiene, J., & Puraite, I. (2016). Impact of anthropogenic pressure on the formation of population structure and genetic diversity of raccoon dog *Nyctereutes procyonoides*. Current Zoology, 62(5), 413–420. https://doi.org/10.1093/cz/zow038
167. Karamon, J., Samorek-Pieróg, M., Moskwa, B., Rózycki, M., Bilska-Zajac, E., Zdybel, J., & Włodarczyk, M. (2016). Intestinal helminths of raccoon dogs (*Nyctereutes procyonoides*) and red foxes (Vulpes vulpes) from the Augustów Primeval Forest (north-eastern Poland). Journal of Veterinary Research (Poland), 60(3), 273–277. https://doi.org/10.1515/jvetres-2016-0042
168. Laurimaa, L., Süld, K., Davison, J., Moks, E., Valdmann, H., & Saarma, U. (2016). Alien species and their zoonotic parasites in native and introduced ranges: The raccoon dog example. Veterinary Parasitology, 219, 24–33. https://doi.org/10.1016/j.vetpar.2016.01.020
169. Maas, M., van den End, S., van Roon, A., Mulder, J., Franssen, F., Dam-Deisz, C., Montizaan, M., & van der Giessen, J. (2016). First findings of Trichinella spiralis and DNA of Echinococcus multilocularis in wild raccoon dogs in the Netherlands. International Journal for Parasitology: Parasites and Wildlife, 5(3), 277–279. https://doi.org/10.1016/j.ijppaw.2016.09.001
170. Oksanen, A., Siles-Lucas, M., Karamon, J., Possenti, A., Conraths, F. J., Romig, T., Wysocki, P., Mannocci, A., Mipatrini, D., La Torre, G., Boufana, B., & Casulli, A. (2016). The geographical distribution and prevalence of Echinococcus multilocularis in animals in the European Union and adjacent countries: A systematic review and meta-analysis. Parasites and Vectors, 9(1), 1–23. https://doi.org/10.1186/s13071-016-1746-4

171. Wodecka, B., Michalik, J., Lane, R. S., Nowak-Chmura, M., & Wierzbicka, A. (2016). Differential associations of Borrelia species with European badgers (Meles meles) and raccoon dogs (*Nyctereutes procyonoides*) in western Poland. Ticks and Tick-Borne Diseases, 7(5), 1010–1016. https://doi.org/10.1016/j.ttbdis.2016.05.008
172. Duscher, T., Hodžić, A., Glawischnig, W., & Duscher, G. G. (2017). The raccoon dog (*Nyctereutes procyonoides*) and the raccoon (*Procyon lotor*)—their role and impact of maintaining and transmitting zoonotic diseases in Austria, Central Europe. Parasitology Research, 116(4), 1411–1416. https://doi.org/10.1007/s00436-017-5405-2
173. Kärssin, A., Häkkinen, L., Niin, E., Peik, K., Vilem, A., Jokelainen, P., & Lassen, B. (2017). Trichinella spp. biomass has increased in raccoon dogs (*Nyctereutes procyonoides*) and red foxes (Vulpes vulpes) in Estonia. Parasites and Vectors, 10(1), 0–12. https://doi.org/10.1186/s13071-017-2571-0
174. Suld, K., Saarma, U., & Valdmann, H. (2017). Home ranges of raccoon dogs in managed and natural areas. PLoS ONE, 12(3), 1–10. https://doi.org/10.1371/journal.pone.0171805
175. Dähnert, L., Conraths, F. J., Reimer, N., Groschup, M. H., & Eiden, M. (2018). Molecular and serological surveillance of Hepatitis E virus in wild and domestic carnivores in Brandenburg, Germany. Transboundary and Emerging Diseases, 65(5), 1377–1380. https://doi.org/10.1111/tbed.12877
176. Elmeros, M., Mikkelsen, D. M. G., Nørgaard, L. S., Pertoldi, C., Jensen, T. H., & Chriél, M. (2018). The diet of feral raccoon dog (*Nyctereutes procyonoides*) and native badger (Meles meles) and red fox (Vulpes vulpes) in Denmark. Mammal Research, 63(4), 405–413. https://doi.org/10.1007/s13364-018-0372-2
177. Hildebrand, J., Buńkowska-Gawlik, K., Adamczyk, M., Gajda, E., Merta, D., Popiołek, M., & Perec-Matysiak, A. (2018). The occurrence of Anaplasmataceae in European populations of invasive carnivores. Ticks and Tick-Borne Diseases, 9(4), 934–937. https://doi.org/10.1016/j.ttbdis.2018.03.018
178. Krüger, H., Väänänen, V. M., Holopainen, S., & Nummi, P. (2018). The new faces of nest predation in agricultural landscapes—a wildlife camera survey with artificial nests. European Journal of Wildlife Research, 64(6). https://doi.org/10.1007/s10344-018-1233-7
179. Petersen, H. H., Al-Sabi, M. N. S., Enemark, H. L., Kapel, C. M. O., Jørgensen, J. A., & Chriél, M. (2018a). Echinococcus multilocularis in Denmark 2012–2015: high local prevalence in red foxes. Parasitology Research, 117(8), 2577–2584. https://doi.org/10.1007/s00436-018-5947-y
180. Tammeleht, E., & Kuuspu, M. (2018). Effect of competition and landscape characteristics on mesocarnivore cohabitation in badger setts. Journal of Zoology, 305(1), 8–16. https://doi.org/10.1111/jzo.12529
181. Cybulska, A., Kornacka, A., & Moskwa, B. (2019). The occurrence and muscle distribution of Trichinella britovi in raccoon dogs (*Nyctereutes procyonoides*) in wildlife in the Głęboki Bród Forest District, Poland. International Journal for Parasitology: Parasites and Wildlife, 9(February), 149–153. https://doi.org/10.1016/j.ijppaw.2019.05.003
182. Dahl, F., & Åhlén, P. A. (2019). Nest predation by raccoon dog *Nyctereutes procyonoides* in the archipelago of northern Sweden. Biological Invasions, 21(3), 743–755. https://doi.org/10.1007/s10530-018-1855-4
183. Ksyonz, I. M., Zezekalo, V. K., Peredera, S. B., Shcherbakova, N. C., Peredera, Z. O., Kone, M. S., Rak, T. M., Kravchenko, S. O., & Kanivets, N. S. (2019). Chlamydial Infection Monitoring Within Wild Mammals in Ukraine. World of Medicine and Biology, 15(67), 227. https://doi.org/10.26724/2079-8334-2019-1-67-227
184. Nummi, P., Väänänen, V. M., Pekkarinen, A. J., Eronen, V., Mikkola-Roos, M., Nurmi, J., Rautiainen, A., & Rusanen, P. (2019b). Alien predation in wetlands – The raccoon dog and waterbird breeding success. Baltic Forestry, 25(2), 228–237. https://doi.org/10.46490/vol25iss2pp228
185. Holopainen, S., Väänänen, V. M., & Fox, A. D. (2020). Landscape and habitat affect frequency of artificial duck nest predation by native species, but not by an alien predator. Basic and Applied Ecology, 48, 52–60. https://doi.org/10.1016/j.baae.2020.07.004
186. Uusitalo, R., Siljander, M., Dub, T., Sane, J., Sormunen, J. J., Pellikka, P., & Vapalahti, O. (2020). Modelling habitat suitability for occurrence of human tick-borne encephalitis (TBE) cases in Finland. Ticks and Tick-Borne Diseases, 11(5), 101457. https://doi.org/10.1016/j.ttbdis.2020.101457

*Ondatra zibethicus*

187. Triplet, P. (2009). Datasheet on *Ondatra zibethicus*. Wallingford (UK): CAB International, Invasive Species Compendium. Available from: http://www.cabi.org/isc.
188. Birnbaum, C. (2013). NOBANIS - Invasive Alien Species Fact Sheet - *Ondatra zibethicus*. Online Database of the European Network on Invasive Alien Species - NOBANIS, 1–11. www.nobanis.org
189. Deputy Direction of Nature. (2016). EU NON-NATIVE ORGANISM RISK ASSESSMENT SCHEME - *Ondatra zibethicus*. 37.
190. Adriana, G., Zsuzsa, K., Mirabela Oana, D., Mircea, G. C., & Viorica, M. (2016). *Giardia duodenalis* genotypes in domestic and wild animals from Romania identified by PCR-RFLP targeting the gdh gene. Veterinary Parasitology, 217, 71–75. https://doi.org/10.1016/j.vetpar.2015.10.017
191. Vermaat, J. E., Bos, B., & Van Der Burg, P. (2016). Why do reed beds decline and fail to re-establish? A case study of Dutch peat lakes. Freshwater Biology, 61(9), 1580–1589. https://doi.org/10.1111/fwb.12801
192. Hurd, J., Berke, O., Poljak, Z., & Runge, M. (2017). Spatial analysis of Leptospira infection in muskrats in Lower Saxony, Germany, and the association with human leptospirosis. Research in Veterinary Science, 114(June), 351–354. https://doi.org/10.1016/j.rvsc.2017.06.015
193. van Loon, E. E., Bos, D., van Hellenberg Hubar, C. J., & Ydenberg, R. C. (2017). A historical perspective on the effects of trapping and controlling the muskrat (*Ondatra zibethicus*) in the Netherlands. Pest Management Science, 73(2), 305–312. https://doi.org/10.1002/ps.4270
194. Ydenberg, R. C., Loon, E. E. Van, Bos, D., & Hemert, H. Van. (2019). Damage to dykes and levees in the - Netherlands is extensive and increases with muskrat (*Ondatra zibethicus*) density. Lutra, 62(1), 39–53.
195. Krügel, M., Pfeffer, M., Król, N., Imholt, C., Baert, K., Ulrich, R. G., & Obiegala, A. (2020). Rats as potential reservoirs for neglected zoonotic Bartonella species in Flanders, Belgium. Parasites and Vectors, 13(1), 1–12. https://doi.org/10.1186/s13071-020-04098-y
196. Stoeckl, K., Denic, M., & Geist, J. (2020). Conservation status of two endangered freshwater mussel species in Bavaria, Germany: Habitat quality, threats, and implications for conservation management. Aquatic Conservation: Marine and Freshwater Ecosystems, 30(4), 647–661. https://doi.org/10.1002/aqc.3310

*Procyon lotor*

197. Gehrt, S. (2009). Datasheet on *Procyon lotor*. Wallingford (UK): CAB International, Invasive Species Compendium. Available from: http://www.cabi.org/isc.
198. Bartoszewicz, M. (2011). NOBANIS - Invasive Alien Species Fact Sheet - *Procyon lotor*. Online Database of the European Network on Invasive Alien Species - NOBANIS, 1–9. http://www.nobanis.org/files/factsheets/Procyon_lotor.pdf
199. Popiołek, M., Szczęsna-Staśkiewicz, J., Bartoszewicz, M., Okarma, H., Smalec, B., & Zalewski, A. (2011). Helminth parasites of an introduced invasive carnivore species, the raccoon (*Procyon lotor* L.), from the Warta Mouth National Park (Poland). Journal of Parasitology, 97(2), 357–360. https://doi.org/10.1645/GE-2525.1
200. Zalewski, A. (2011). GB NON-NATIVE SPECIES RISK ASSESSMENT - *Procyon lotor*.
201. Beltrán-Beck, B., García, F. J., & Gortázar, C. (2012). Raccoons in Europe: Disease hazards due to the establishment of an invasive species. European Journal of Wildlife Research, 58(1), 5–15. https://doi.org/10.1007/s10344-011-0600-4
202. García, J. T., García, F. J., Alda, F., González, J. L., Aramburu, M. J., Cortés, Y., Prieto, B., Pliego, B., Pérez, M., Herrera, J., & García-Román, L. (2012). Recent invasion and status of the raccoon (*Procyon lotor*) in Spain. Biological Invasions, 14(7), 1305–1310. https://doi.org/10.1007/s10530-011-0157-x
203. Vos, A., Ortmann, S., Kretzschmar, A. S., Köhnemann, B., & Michler, F. (2012). The raccoon (*Procyon lotor*) as potential rabies reservoir species in Germany: A risk assessment. Berliner Und Munchener Tierarztliche Wochenschrift, 125(5/6), 228–235. https://doi.org/10.2376/0005-9366-125-222
204. Alda, F., Ruiz-López, M. J., García, F. J., Gompper, M. E., Eggert, L. S., & García, J. T. (2013). Genetic evidence for multiple introduction events of raccoons (*Procyon lotor*) in Spain. Biological Invasions, 15(3), 687–698. https://doi.org/10.1007/s10530-012-0318-6

205. Frantz, A. C., Heddergott, M., Lang, J., Schulze, C., Ansorge, H., Runge, M., Braune, S., Michler, F. U., Wittstatt, U., Hoffmann, L., Hohmann, U., Michler, B. A., Van Den Berge, K., & Horsburgh, G. J. (2013). Limited mitochondrial DNA diversity is indicative of a small number of founders of the German raccoon (*Procyon lotor*) population. European Journal of Wildlife Research, 59(5), 665–674. https://doi.org/10.1007/s10344-013-0719-6
206. Rentería-Solís, Z. M., Hamedy, A., Michler, F. U., Michler, B. A., Lücker, E., Stier, N., Wibbelt, G., & Riehn, K. (2013). Alaria alata mesocercariae in raccoons (*Procyon lotor*) in Germany. Parasitology Research, 112(10), 3595–3600. https://doi.org/10.1007/s00436-013-3547-4
207. Vos, A., Nolden, T., Habla, C., Finke, S., Freuling, C. M., Teifke, J., & Müller, T. (2013). Raccoons (*Procyon lotor*) in Germany as potential reservoir species for Lyssaviruses. European Journal of Wildlife Research, 59(5), 637–643. https://doi.org/10.1007/s10344-013-0714-y
208. Biedrzycka, A., Zalewski, A., Bartoszewicz, M., Okarma, H., & Jędrzejewska, E. (2014). The genetic structure of raccoon introduced in Central Europe reflects multiple invasion pathways. Biological Invasions, 16(8), 1611–1625. https://doi.org/10.1007/s10530-013-0595-8
209. Gabrys, G., Nowaczyk, J., Wazna, A., Koscielska, A., Nowakowski, K., & Cichocki, J. (2014). Expansion of the raccoon *Procyon lotor* in Poland. Zeszyty Naukowe Uniwersytetu Szczecinskiego, 844, 169–181.
210. Karamon, J., Kochanowski, M., Cencek, T., Bartoszewicz, M., & Kusyk, P. (2014). Gastrointestinal helminths of raccoons (*Procyon lotor*) in western Poland (Lubuskie province) - with particular regard to Baylisascaris procyonis. Bulletin of the Veterinary Institute in Pulawy, 58(4), 547–552. https://doi.org/10.2478/bvip-2014-0084
211. Rentería-Solís, Z., Min, A. M., Alasaad, S., Müller, K., Michler, F. U., Schmäschke, R., Wittstatt, U., Rossi, L., & Wibbelt, G. (2014a). Genetic epidemiology and pathology of raccoon-derived Sarcoptes mites from urban areas of Germany. Medical and Veterinary Entomology, 28(SUPPL.1), 98–103. https://doi.org/10.1111/mve.12079
212. Rentería-Solís, Z., Förster, C., Aue, A., Wittstatt, U., Wibbelt, G., & König, M. (2014b). Canine distemper outbreak in raccoons suggests pathogen interspecies transmission amongst alien and native carnivores in urban areas from Germany. Veterinary Microbiology, 174(1–2), 50–59. https://doi.org/10.1016/j.vetmic.2014.08.034
213. Fischer, M. L., Hochkirch, A., Heddergott, M., Schulze, C., Anheyer-Behmenburg, H. E., Lang, J., Michler, F. U., Hohmann, U., Ansorge, H., Hoffmann, L., Klein, R., & Frantz, A. C. (2015). Historical invasion records can be misleading: Genetic evidence for multiple introductions of invasive raccoons (*Procyon lotor*) in Germany. PLoS ONE, 10(5), 1–17. https://doi.org/10.1371/journal.pone.0125441
214. Mori, E., Mazza, G., Menchetti, M., Panzeri, M., Gager, Y., Bertolino, S., & Di Febbraro, M. (2015). The masked invader strikes again: The conquest of Italy by the Northern raccoon. Hystrix, 26(1), 1–5. https://doi.org/10.4404/hystrix-26.1-11035
215. Farashi, A., Naderi, M., & Safavian, S. (2016). Predicting the potential invasive range of raccoon in the world. Polish Journal of Ecology, 64(4), 594–600. https://doi.org/10.3161/15052249PJE2016.64.4.014
216. Fischer, M. L., Sullivan, M. J. P., Greiser, G., Guerrero-Casado, J., Heddergott, M., Hohmann, U., Keuling, O., Lang, J., Martin, I., Michler, F. U., Winter, A., & Klein, R. (2016). Assessing and predicting the spread of non-native raccoons in Germany using hunting bag data and dispersal weighted models. Biological Invasions, 18(1), 57–71. https://doi.org/10.1007/s10530-015-0989-x
217. Leśniańska, K., Perec-Matysiak, A., Hildebrand, J., Buńkowska-Gawlik, K., Piróg, A., & Popiołek, M. (2016). Cryptosporidium spp. and Enterocytozoon bieneusi in introduced raccoons (*Procyon lotor*)—first evidence from Poland and Germany. Parasitology Research, 115(12), 4535–4541. https://doi.org/10.1007/s00436-016-5245-5
218. Nowakiewicz, A., Zieba, P., Ziółkowska, G., Gnat, S., Muszyńska, M., Tomczuk, K., Dziedzic, B. M., Ulbrych, Ł., & Trościańczyk, A. (2016). Free-living species of carnivorous mammals in Poland: Red fox, beech marten, and raccoon as a potential reservoir of Salmonella, Yersinia, Listeria spp. and coagulase-positive Staphylococcus. PLoS ONE, 11(5), 1–16. https://doi.org/10.1371/journal.pone.0155533
219. Fischer, M. L., Salgado, I., Beninde, J., Klein, R., Frantz, A. C., Heddergott, M., Cullingham, C. I., Kyle, C. J., & Hochkirch, A. (2017). Multiple founder effects are followed by range expansion and admixture during the invasion process of the raccoon (*Procyon lotor*) in Europe. Diversity and Distributions, 23(4), 409–420. https://doi.org/10.1111/ddi.12538

220. Hechinger, S., Scheffold, S., Hamann, H. P., & Zschöck, M. (2017). Detection of canine adenovirus 1 in red foxes (Vulpes vulpes) and raccoons (*Procyon lotor*) in Germany with a TaqMan real-time PCR assay. Journal of Veterinary Diagnostic Investigation, 29(5), 741–746. https://doi.org/10.1177/1040638717712331
221. Heddergott, M., Frantz, A. C., Stubbe, M., Stubbe, A., Ansorge, H., & Osten-Sacken, N. (2017). Seroprevalence and risk factors of Toxoplasma gondii infection in invasive raccoons (*Procyon lotor*) in Central Europe. Parasitology Research, 116(8), 2335–2340. https://doi.org/10.1007/s00436-017-5518-7
222. Bencatel, J., Ferreira, C. C., Márcia Barbosa, A., Rosalino, L. M., & Álvares, F. (2018). Research trends and geographical distribution of mammalian carnivores in Portugal (SW Europe). PLoS ONE, 13(11), 1–20. https://doi.org/10.1371/journal.pone.0207866
223. Cybulska, A., Skopek, R., Kornacka, A., Popiołek, M., Piróg, A., Laskowski, Z., & Moskwa, B. (2018). First detection of Trichinella pseudospiralis infection in raccoon (*Procyon lotor*) in Central Europe. Veterinary Parasitology, 254(March), 114–119. https://doi.org/10.1016/j.vetpar.2018.03.007
224. Kornacka, A., Cybulska, A., Popiołek, M., Kuśmierek, N., & Moskwa, B. (2018). Survey of Toxoplasma gondii and Neospora caninum in raccoons (*Procyon lotor*) from the Czech Republic, Germany and Poland. Veterinary Parasitology, 262, 47–50. https://doi.org/10.1016/j.vetpar.2018.09.006
225. Litvinchuk, S. N., & Kidov, A. A. (2018). Distribution and conservation status of the caucasian parsley frog, pelodytes caucasicus (amphibia: Anura). Nature Conservation Research, 3, 51–60. https://doi.org/10.24189/ncr.2018.053
226. Osten-Sacken, N., Heddergott, M., Schleimer, A., Anheyer-Behmenburg, H. E., Runge, M., Horsburgh, G. J., Camp, L., Nadler, S. A., & Frantz, A. C. (2018). Similar yet different: co-analysis of the genetic diversity and structure of an invasive nematode parasite and its invasive mammalian host. International Journal for Parasitology, 48(3–4), 233–243. https://doi.org/10.1016/j.ijpara.2017.08.013
227. Rentería-Solís, Z., Birka, S., Schmäschke, R., Król, N., & Obiegala, A. (2018). First detection of Baylisascaris procyonis in wild raccoons (*Procyon lotor*) from Leipzig, Saxony, Eastern Germany. Parasitology Research, 117(10), 3289–3292. https://doi.org/10.1007/s00436-018-5988-2
228. Risueño, J., Ortuño, M., Pérez-Cutillas, P., Goyena, E., Maia, C., Cortes, S., Campino, L., Bernal, L. J., Muñoz, C., Arcenillas, I., Martínez-Rondán, F. J., Gonzálvez, M., Collantes, F., Ortiz, J., Martínez-Carrasco, C., & Berriatua, E. (2018). Epidemiological and genetic studies suggest a common Leishmania infantum transmission cycle in wildlife, dogs and humans associated to vector abundance in Southeast Spain. Veterinary Parasitology, 259(May), 61–67. https://doi.org/10.1016/j.vetpar.2018.05.012
229. Salgado, I. (2018). Is the raccoon (*Procyon lotor*) out of control in Europe? Biodiversity and Conservation, 27(9), 2243–2256. https://doi.org/10.1007/s10531-018-1535-9
230. Boscherini, A., Mazza, G., Menchetti, M., Laurenzi, A., & Mori, E. (2019). Time is running out! Rapid range expansion of the invasive northern raccoon in central Italy. Mammalia. https://doi.org/10.1515/mammalia-2018-0151
231. Fiderer, C., Göttert, T., & Zeller, U. (2019). Spatial interrelations between raccoons (*Procyon lotor*), red foxes (Vulpes vulpes), and ground-nesting birds in a Special Protection Area of Germany. European Journal of Wildlife Research, 65(1). https://doi.org/10.1007/s10344-018-1249-z
232. Louppe, V., Leroy, B., Herrel, A., & Veron, G. (2019). Current and future climatic regions favourable for a globally introduced wild carnivore, the raccoon *Procyon lotor*. Scientific Reports, 9(1), 1–13. https://doi.org/10.1038/s41598-019-45713-y
233. Schulze, C., Schatz, J., Dohrmann, E., & Wohlsein, P. (2019). Molecular epidemiology of canine adeno-virus type 1 (Cadv-1) in free-ranging small carnivores in the berlin-brandenburg region, Germany. A preliminary study. Berliner Und Munchener Tierarztliche Wochenschrift, 132(9–10), 476–480. https://doi.org/10.2376/0005-9366-18071
234. Heddergott, M., Frantz, A. C., Pohl, D., Osten-Sacken, N., & Steinbach, P. (2020a). Detection of Cryptosporidium spp. Infection in Wild Raccoons (*Procyon lotor*) from Luxembourg Using an ELISA Approach. Acta Parasitologica, June. https://doi.org/10.2478/s11686-020-00234-x
235. Heddergott, M., Steinbach, P., Schwarz, S., Anheyer-Behmenburg, H. E., Sutor, A., Schliephake, A., Jeschke, D., Striese, M., Müller, F., Meyer-Kayser, E., Stubbe, M., Osten-Sacken, N., Krüger, S., Gaede, W., Runge, M., Hoffmann, L., Ansorge, H., Conraths, F. J., & Frantz, A. C. (2020b). Geographic

distribution of raccoon roundworm, Baylisascaris procyonis, Germany and Luxembourg. Emerging Infectious Diseases, 26(4), 821–823. https://doi.org/10.3201/eid2604.191670

236. Mazzamuto, M. V., Panzeri, M., Bisi, F., Wauters, L. A., Preatoni, D., & Martinoli, A. (2020). When management meets science: adaptive analysis for the optimization of the eradication of the Northern raccoon (*Procyon lotor*). Biological Invasions, 22(10), 3119–3130. https://doi.org/10.1007/s10530-020-02313-6

*Sciurus carolinensis*

237. IUCN/SSC Invasive Species Specialist Group (ISSG) (2005). Datasheet on *Sciurus carolinensis*. Wallingford (UK): CAB International, Invasive Species Compendium. Available from: http://www.cabi. org/isc.
238. Bertolino, S., Martinoli, A., & Wauters, L. (2014a). Risk Assessment for *Sciurus carolinensis* (Grey Squirrel). 251–291.
239. Bertolino, S., di Montezemolo, N. C., Preatoni, D. G., Wauters, L. A., & Martinoli, A. (2014b). A grey future for Europe: *Sciurus carolinensis* is replacing native red squirrels in Italy. Biological Invasions, 16(1), 53–62. https://doi.org/10.1007/s10530-013-0502-3
240. Collins, L. M., Warnock, N. D., Tosh, D. G., McInnes, C., Everest, D., Montgomery, W. I., Scantlebury, M., Marks, N., Dick, J. T. A., & Reid, N. (2014). Squirrelpox virus: Assessing prevalence, transmission and environmental degradation. PLoS ONE, 9(2), 1–8. https://doi.org/10.1371/journal.pone.0089521
241. Gurnell, J., Lurz, P., & Bertoldi, W. (2014). The changing patterns in the distribution of red and grey squirrels in the North of England and Scotland between 1991 and 2010 based on volunteer surveys. Hystrix, 25(2), 83–89. https://doi.org/10.4404/hystrix-25.2-9988
242. Romeo, C., Wauters, L. A., Ferrari, N., Lanfranchi, P., Martinoli, A., Pisanu, B., Preatoni, D. G., & Saino, N. (2014a). Macroparasite fauna of alien grey squirrels (*Sciurus carolinensis*): Composition, variability and implications for native species. PLoS ONE, 9(2), 1–8. https://doi.org/10.1371/journal.pone.0088002
243. Romeo, C., Ferrari, N., Rossi, C., Everest, D. J., Grierson, S. S., Lanfranchi, P., Martinoli, A., Saino, N., Wauters, L. A., & Hauffe, H. C. (2014b). Ljungan virus and an adenovirus in Italian squirrel populations. Journal of Wildlife Diseases, 50(2), 409–411. https://doi.org/10.7589/2013-10-260
244. Bonnington, C., Gaston, K. J., & Evans, K. L. (2015). Ecological traps and behavioural adjustments of urban songbirds to fine-scale spatial variation in predator activity. Animal Conservation, 18(6), 529–538. https://doi.org/10.1111/acv.12206
245. Millins, C., Magierecka, A., Gilbert, L., Edoff, A., Brereton, A., Kilbride, E., Denwood, M., Birtles, R., & Bieka, R. (2015). An invasive mammal (the gray squirrel, *Sciurus carolinensis*) commonly hosts diverse and atypical genotypes of the zoonotic pathogen Borrelia burgdorferi Sensu lato. Applied and Environmental Microbiology, 81(13), 4236–4245. https://doi.org/10.1128/AEM.00109-15
246. Romeo, C., Ferrari, N., Lanfranchi, P., Saino, N., Santicchia, F., Martinoli, A., & Wauters, L. A. (2015). Biodiversity threats from outside to inside: effects of alien grey squirrel (*Sciurus carolinensis*) on helminth community of native red squirrel (*Sciurus vulgaris*). Parasitology Research, 114(7), 2621–2628. https://doi.org/10.1007/s00436-015-4466-3
247. Shuttleworth, C. M., Signorile, A. L., Everest, D. J., Duff, J. P., & Lurz, P. W. W. (2015). Assessing causes and significance of red squirrel (*Sciurus vulgaris*) mortality during regional population restoration: An applied conservation perspective. Hystrix, 26(2), 69–75. https://doi.org/10.4404/hystrix-26.2-11166
248. Stritch, C., Naulty, F., Zintl, A., Callanan, J. J., McCullough, M., Deane, D., Marnell, F., & McMahon, B. J. (2015). Squirrelpox virus reservoir expansion on the east coast of Ireland. European Journal of Wildlife Research, 61(3), 483–486. https://doi.org/10.1007/s10344-015-0909-5
249. Goldstein, E. A., Butler, F., & Lawton, C. (2016). Modeling future range expansion and management strategies for an invasive squirrel species. Biological Invasions, 18(5), 1431–1450. https://doi.org/10.1007/s10530-016-1092-7
250. Mori, E., Amerini, R., Mazza, G., Bertolino, S., Battiston, R., Sforzi, A., & Menchetti, M. (2016b). Alien shades of grey: New occurrences and relevant spread of *Sciurus carolinensis* in Italy. European Journal of Ecology, 2(1), 13–20. https://doi.org/10.1515/eje-2016-0002

251. Signorile, A. L., Lurz, P. W. W., Wang, J., Reuman, D. C., & Carbone, C. (2016a). Mixture or mosaic? Genetic patterns in UK grey squirrels support a human-mediated "long-jump" invasion mechanism. Diversity and Distributions, 22(5), 566–577. https://doi.org/10.1111/ddi.12424
252. Signorile, A. L., Reuman, D. C., Lurz, P. W. W., Bertolino, S., Carbone, C., & Wang, J. (2016b). Using DNA profiling to investigate human-mediated translocations of an invasive species. Biological Conservation, 195, 97–105. https://doi.org/10.1016/j.biocon.2015.12.026
253. Hanmer, H. J., Thomas, R. L., & Fellowes, M. D. E. (2017). Provision of supplementary food for wild birds may increase the risk of local nest predation. Ibis, 159(1), 158–167. https://doi.org/10.1111/ibi.12432
254. Hanmer, H. J., Thomas, R. L., & Fellowes, M. D. E. (2018). Introduced Grey Squirrels subvert supplementary feeding of suburban wild birds. Landscape and Urban Planning, 177(March 2017), 10–18. https://doi.org/10.1016/j.landurbplan.2018.04.004
255. Romeo, C., Lecollinet, S., Caballero, J., Isla, J., Luzzago, C., Ferrari, N., & García-Bocanegra, I. (2018). Are tree squirrels involved in the circulation of flaviviruses in Italy? Transboundary and Emerging Diseases, 65(5), 1372–1376. https://doi.org/10.1111/tbed.12874
256. Santicchia, F., Dantzer, B., van Kesteren, F., Palme, R., Martinoli, A., Ferrari, N., & Wauters, L. A. (2018). Stress in biological invasions: Introduced invasive grey squirrels increase physiological stress in native Eurasian red squirrels. Journal of Animal Ecology, 87(5), 1342–1352. https://doi.org/10.1111/1365-2656.12853
257. Sheehy, E., Sutherland, C., O'Reilly, C., & Lambin, X. (2018). The enemy of my enemy is my friend: Native pine marten recovery reverses the decline of the red squirrel by suppressing grey squirrel populations. Proceedings of the Royal Society B: Biological Sciences, 285(1874). https://doi.org/10.1098/rspb.2017.2603
258. Romeo, C., McInnes, C. J., Dale, T. D., Shuttleworth, C., Bertolino, S., Wauters, L. A., & Ferrari, N. (2019). Disease, invasions and conservation: no evidence of squirrelpox virus in grey squirrels introduced to Italy. Animal Conservation, 22(1), 14–23. https://doi.org/10.1111/acv.12433
259. Broughton, R. K. (2020). Current and future impacts of nest predation and nest-site competition by invasive eastern grey squirrels *Sciurus carolinensis* on European birds. Mammal Review, 50(1), 38–51. https://doi.org/10.1111/mam.12174
260. McNicol, C. M., Bavin, D., Bearhop, S., Ferryman, M., Gill, R., Goodwin, C. E. D., MacPherson, J., Silk, M. J., & McDonald, R. A. (2020). Translocated native pine martens Martes martes alter short-term space use by invasive non-native grey squirrels *Sciurus carolinensis*. Journal of Applied Ecology, 57(5), 903–913. https://doi.org/10.1111/1365-2664.13598
261. Santicchia, F., Wauters, L. A., Piscitelli, A. P., Van Dongen, S., Martinoli, A., Preatoni, D., Romeo, C., & Ferrari, N. (2020). Spillover of an alien parasite reduces expression of costly behaviour in native host species. Journal of Animal Ecology, 89(7), 1559–1569. https://doi.org/10.1111/1365-2656.13219
262. Twining, J. P., Montgomery, W. I., Price, L., Kunc, H. P., & Tosh, D. G. (2020). Native and invasive squirrels show different behavioural responses to scent of a shared native predator. Royal Society Open Science, 7(2). https://doi.org/10.1098/rsos.191841

1189

**Appendix S5.** List of pathogens known to have been recorded to infect the study species in Europe and list of additional references.

| Species | Pathogen | Zoonotic | Country | Prevalence | Reference | Notes |
|---|---|---|---|---|---|---|
| *Atlantoxerus getulus* | *Acanthamoeba* spp. | YES | ES | 23.50% | Lorenzo-Morales *et al.*, 2007 | |
| *Callosciurus erythraeus* | Capillariinae | YES | IT | 1% | Mazzamuto *et al.*, 2016 | |
| | *Ceratophyllus s. sciurorum* | NO | IT | 50% | Mazzamuto *et al.*, 2016 | |
| | *Cryptosporidium* spp. | YES | IT | 2.80% | Prediger *et al.*, 2017 | |
| | *Ctenophtalmus agyrtes sardiniensis* | NO | IT | 1% | Mazzamuto *et al.*, 2016 | |
| | *Ctenophtalmus* sp. | NO | IT | 1% | Mazzamuto *et al.*, 2016 | |
| | *Eimeria* spp. | YES | IT | 4.10% | Hofmannová *et al.*, 2016 | |
| | *Ixodes ricinus* | | IT | 47% | Mazzamuto *et al.*, 2016 | |
| | *Mycobacterium leprae* | YES | IT | 0% | Schilling *et al.*, 2019 | |
| | *Mycobacterium leprae* | YES | FR | 0% | Schilling *et al.*, 2019 | |
| | Spiruridae | NO | IT | 1% | Mazzamuto *et al.*, 2016 | |
| | *Strongyloides callosciureus* | NO | IT | 1% | Mazzamuto *et al.*, 2016 | |
| | *Strongyloides* sp. | YES | IT | 1% | Mazzamuto *et al.*, 2016 | |
| | *Trichuris muris* | NO | IT | 4% | Mazzamuto *et al.*, 2016 | |
| | Trombiculidae | NO | IT | 7% | Mazzamuto *et al.*, 2016 | |
| | *Trypanoxyuris sciuri* | NO | IT | 5% | Mazzamuto *et al.*, 2016 | |
| *Callosciurus finlaysonii* | *Cryptococcus neoformans* | YES | IT | 5.60% | Iatta *et al.*, 2015 | |
| | *Debaryomyces hansenii* | YES | IT | 0.80% | Iatta *et al.*, 2015 | |
| | *Dicrocoelium dendriticum* | YES | IT | 33.30% | d'Ovidio *et al.* 2014 | |
| | *Hanseniaspora thailandica* | NO | IT | 3.20% | Iatta *et al.*, 2015 | |

| Species | Pathogen | Zoonotic | Country | Prevalence | Reference | Notes |
|---|---|---|---|---|---|---|
| *Callosciurus finlaysonii* | *Meyerozyma guilliermondii* | YES | IT | 0.80% | Iatta *et al.*, 2015 | |
| *Castor canadensis* | *Francisella tularensis* | YES | SE | | Sissonen *et al.*, 2015 | Samples analysed were already infected. |
| *Cervus nippon* | *Anaplasma phagocytophilum* | YES | UK | 50% | Robinson *et al.*, 2009 | |
| | *Ashworthius sidemi* | NO | RU | | Panova *et al.*, 2017 | Probably introduced in Europe with *C. nippon.* |
| | *Babesia* spp. | YES | CZ | 21.90% | Hrazdilová *et al.*, 2020 | |
| | *BlueTongue Virus* (BTV) | NO | IE | 0% | Graham *et al.,* 2017 | Pooled prevalence: sika + fallow + red deer. |
| | *Border Disease Virus* (BDV) | NO | CZ | 0% | Sedlak *et al.*, 2009 | |
| | *Bovine HerpesVirus-1* (BoHV-1) | NO | IE | 1.80% | Graham *et al.,* 2017 | Pooled prevalence: sika + fallow + red deer. |
| | *Bovine Viral Diarrhoea Virus* (BVDV) | NO | CZ | 0% | Sedlak *et al.*, 2009 | |
| | *Bovine Viral Diarrhoea Virus* (BVDV) | NO | IE | 1.50% | Graham *et al.,* 2017 | Pooled prevalence: sika + fallow + red deer. |
| | *Hepatitis E Virus* (HEV) | YES | DE | 0% | Trojnar *et al.*, 2020 | |
| | *Hepatitis E Virus* (HEV) | YES | PL | 0% | Larska *et al.*, 2015 | |
| | *Hepatitis E Virus* (HEV) | YES | CZ | 0% | Kubankova *et al.*, 2015 | |
| | *Lipoptena fortisetosa* | NO | EE | | Mihalca *et al.* 2019 | Probably introduced in Europe with *C. nippon.* |
| | *Onchocerca flexuosa* | NO | CZ | 16.70% | Dykova & Blazek, 1972 | Can be a host. |
| | *Sarcocystis* spp. | YES | LT | 100% | Prakas *et al.*, 2016 | Farm bred animals. |
| | *Sarcocystis* spp. | YES | LT | 92% | Rudaitytė-Lukošienė *et al.*, 2018 | |
| | *Schmallenberg Virus* (SBV) | NO | IE | 9.70% | Graham *et al.,* 2017 | Pooled prevalence: sika + fallow + red deer. |
| | *Toxoplasma gondii* | YES | CZ | 50% | Lorencova *et al.,* 2015 | Antibodies. DNA prevalence: 0%. |
| | *Trichuris discolor* | NO | CZ | 5.20% | Nechybová *et al.*, 2018 | |
| | *Trichuris ovis* | NO | CZ | 1.70% | Nechybová *et al.*, 2018 | |
| | *Wehrdickmansia cervipedis* | NO | CZ | 16.70% | Dykova & Blazek, 1972 | |
| *Eutamias sibiricus* | *Aonchotheca annulosa* | NO | FR | 47% | Pisanu *et al.*, 2007 | |

| Species | Pathogen | Zoonotic | Country | Prevalence | Reference | Notes |
|---|---|---|---|---|---|---|
| *Eutamias sibiricus* | *Aonchotheca annulosa* | NO | FR | 40.50% | Pisanu *et al.*, 2009 | |
| | Ascaroidea | YES | FR | 2.40% | Pisanu *et al.*, 2009 | |
| | *Borrelia burgdoferi sensu lato* | YES | FR | 33.30% | Vourc'h *et al.,* 2007 | |
| | *Borrelia burgdoferi sensu lato* | YES | FR | 35% | Marsot *et al.,* 2011 | |
| | *Borrelia burgdoferi sensu lato* | YES | FR | 5%-60% | Marsot *et al.,* 2013 | |
| | *Borrelia lusitaniae* | YES | IT | | Mori *et al.*, 2018b | *B. lusitaniae* and *R. monacensis* were present in ticks of the chipmunks. |
| | *Brevistriata skrjabini* | | FR | 90.5% | Pisanu *et al.,* 2009 | |
| | *Brevistriata skrjabini* | | FR | 87% | Pisanu *et al.,* 2007 | |
| | *Hymenolepis* spp. | YES | IT | 0% | d'Ovidio *et al.,* 2015 | |
| | *Mycobacterium leprae* | YES | FR | 0% | Schilling *et al.,* 2019 | |
| | Oxyuridea | YES | FR | 2.40% | Pisanu *et al.*, 2009 | |
| | *Rickettsia monacensis* | YES | IT | | Mori *et al.*, 2018b | *B. lusitaniae* and *R. monacensis* were present in ticks of the chipmunks. |
| | *Strongyloides callosciureus* | NO | FR | 19% | Pisanu *et al.,* 2009 | |
| | *Trichostrongyloidea* sp. | YES | FR | 7.10% | Pisanu *et al.*, 2009 | |
| | *Trichuris* sp. | YES | FR | 9.50% | Pisanu *et al.*, 2009 | |
| *Muntiacus reevesi* | *Anaplasma phagocytophilum* | YES | UK | 1% | Duscher *et al.,* 2020 | |
| | *Bovine Viral Diarrhoea Virus* (BVDV) | NO | IE | | McKillen *et al.*, 2017 | Can act as a reservoir. Preliminary study. |
| | *Foot and Mouth Disease Virus* (FMDV) | NO | UK | | Gibbs *et al.* 1975 | Samples analysed were already infected. |
| | *Ixodes ricinus* | | UK | | GB Non-Native Species Secretariat, 2011 | |
| | *Mycobacterium bovis* | YES | UK | | Ward & Smith, 2012 | Can act as a host. |
| *Myocastor coypus* | *Cryptosporidium* spp. | YES | IT | 0% | Zanzani *et al.*, 2016 | |
| | *Cryptosporidium* spp. | YES | CZ | 0% | Kellnerová *et al.,* 2017 | |
| | *Eimeria coypi* | NO | CZ | 37% | Nechybová *et al.*, 2018 | Faecal analysis of farm-bred animals. |

| Species | Pathogen | Zoonotic | Country | Prevalence | Reference | Notes |
|---|---|---|---|---|---|---|
| *Myocastor coypus* | *Eimeria coypi* | NO | CZ | 60% | Nechybová *et al.*, 2018 | Faecal analysis of wild animals. |
| | *Eimeria coypi* | NO | IT | 86.30% | Zanzani *et al.*, 2016 | |
| | *Eimeria myopotami* | NO | CZ | 5% | Nechybová *et al.*, 2018 | Faecal analysis of farm-bred animals. |
| | *Eimeria nutriae* | NO | CZ | 45% | Nechybová *et al.*, 2018 | Faecal analysis of wild animals. |
| | *Eimeria nutriae* | NO | CZ | 23% | Nechybová *et al.*, 2018 | Faecal analysis of farm-bred animals. |
| | *Eimeria seideli* | NO | CZ | 26% | Nechybová *et al.*, 2018 | Faecal analysis of farm-bred animals. |
| | *Eimeria seideli* | NO | IT | 6.80% | Zanzani *et al.*, 2016 | |
| | *Escherichia coli* | YES | IT | 4.50% | Zanzani *et al.*, 2016 | |
| | *Francisella tularensis* | YES | DE | 0% | Schulze *et al.,* 2016 | |
| | *Giardia duodenalis (Giardia lamblia)* | YES | IT | 0% | Zanzani *et al.*, 2016 | |
| | *Hepatitis E Virus* (HEV) | YES | IT | 0% | Serracca *et al.,* 2015 | |
| | *Leptospira interrogans* | YES | IT | 44.90% | Zanzani *et al.*, 2016 | Antibodies. Humans are accidental hosts. |
| | *Leptospira* spp. | YES | IT | 32.90% | Bertelloni *et al.*, 2019 | |
| | *Leptospira* spp. | YES | IT | 44.90% | Zanzani *et al.*, 2016 | |
| | *Leptospira* spp. | YES | IT | 27.90% | Fratini *et al.*, 2015 | Antibodies. Prevalence 9.8% by PCR, 0% by bacteriological examination. |
| | *Leptospira* spp. | YES | FR | 64%-76% | Vein *et al.,* 2014 | Antibodies. |
| | *Leptospira* spp. | YES | FR | 42% | Ayral *et al.,* 2020 | Antibodies. |
| | *Leptospira* spp. | YES | FR | 16.50%-66% | Michel *et al.*, 2001 | Antibodies. |
| | *Salmonella* spp. | YES | IT | 0% | Zanzani *et al.*, 2016 | |
| | *Staphylococcus aureus* | YES | IT | 10.10% | Zanzani *et al.*, 2016 | |
| | *Streptococcus* spp. | YES | IT | 3.40% | Zanzani *et al.*, 2016 | |
| | *Strongyloides myopotami* | YES | CZ | 25% | Nechybová *et al.,* 2018 | Necropsy on farm-bred animals. |
| | *Strongyloides myopotami* | YES | IT | 63.40% | Zanzani *et al.*, 2016 | |
| | *Strongyloides* sp. | YES | CZ | 30% | Nechybová *et al.,* 2018 | Faecal analysis of wild animals. |
| | *Strongyloides* sp. | YES | CZ | 11.50% | Nechybová *et al.,* 2018 | Faecal analysis of farm-bred animals. |

| Species | Pathogen | Zoonotic | Country | Prevalence | Reference | Notes |
|---|---|---|---|---|---|---|
| *Myocastor coypus* | *Toxoplasma gondii* | YES | IT | 28.9% | Zanzani *et al.*, 2016 | Antibodies. |
| | *Toxoplasma gondii* | YES | IT | 59.40% | Nardoni *et al.*, 2011 | Antibodies. Prevalence 52.2% by PCR. |
| | *Trichostrongylus duretteae* | NO | IT | 28.10% | Zanzani *et al.*, 2016 | |
| | *Trichostrongylus* sp. | YES | CZ | 4% | Nechybová *et al.*, 2018 | Faecal analysis of farm-bred animals. |
| | *Trichuris myocastoris* | | CZ | 40% | Nechybová *et al.*, 2018 | Necropsy on farm-bred animals. |
| | *Trichuris* sp. | YES | CZ | 5% | Nechybová *et al.,* 2018 | Faecal analysis of wild animals. |
| | *Trichuris* sp. | YES | CZ | 57% | Nechybová *et al.,* 2018 | Faecal analysis of farm-bred animals. |
| *Neovison vison* | *Aelurostrongylus* spp. | NO | ES | 2% | Martínez-Rondán *et al.*, 2017 | |
| | *Alaria alata* | YES | LT | 7.60% | Nugaraitė *et al.*, 2018 | Mesocercariae. |
| | *Aleutian Disease Virus* (ADV) | NO | ES | | Mañas *et al.* 2001 | ADV DNA was detected by PCR in 28.57% of the carcasses tested. |
| | *Angiostrongylus daskalovi* | NO | ES | 6% | Martínez-Rondán *et al.*, 2017 | |
| | *Angiostrongylus vasorum* | NO | DK | 0.80% | Lemming *et al.,* 2020 | |
| | *Aonchotheca annulosa* | NO | ES | 8% | Martínez-Rondán *et al.*, 2017 | |
| | *Aonchotheca putorii* | YES | ES | 54% | Martínez-Rondán *et al.*, 2017 | |
| | *Aonchotheca putorii* | YES | LT | 33.30%-50% | Nugaraitė *et al.*, 2018 | |
| | *Canine ParvoVirus* (CPV) | NO | PT | 0% | Miranda *et al.,* 2017 | |
| | *Capillaria plica (Pearsonema plica)* | NO | DK | 0% | Petersen *et al.,* 2018b | |
| | *Crenosoma melesi* | | ES | 10% | Martínez-Rondán *et al.*, 2017 | |
| | *Crenosoma schachmatovae* | | LT | 10.20%-15% | Nugaraitė *et al.*, 2018 | |
| | *Crenosoma vulpis* | NO | DK | 5.70% | Lemming *et al.,* 2020 | |
| | *Cryptosporidium* spp. | YES | CZ | 1% | Kellnerová *et al.,* 2017 | |
| | *Cystoisospora* spp. | YES | DK | 11% | Petersen *et al.*, 2020 | |
| | *Echinococcus* spp. | YES | PL | 14.20% | Kołodziej-Sobocińska *et al.,* 2020 | |
| | *Ehrlichia canis* | YES | ES | 0% | Criado-Fornelio *et al.,* 2018 | |

| Species | Pathogen | Zoonotic | Country | Prevalence | Reference | Notes |
|---|---|---|---|---|---|---|
| *Neovison vison* | *Eucoleus aerophilus* | YES | LT | 10%-15.30% | Nugaraitė *et al.*, 2018 | |
| | *Francisella tularensis* | YES | DE | 0% | Schulze *et al.,* 2016 | |
| | *Hepatozoon* spp. | NO | ES | 0% | Criado-Fornelio *et al.,* 2018 | |
| | *Influenza A Viruses* (IAV) | YES | ES | 2.20% | Gholipour *et al.,* 2017 | |
| | *Isthmiophora melis* | NO | LT | 75% | Nugaraitė *et al.*, 2017 | |
| | *Isthmiophora melis* | NO | LT | 70%-77% | Nugaraitė *et al.*, 2018 | |
| | *Mesocestoides* spp. | YES | LT | 5%-7.60% | Nugaraitė *et al.*, 2018 | |
| | *Molineus patens* | NO | LT | 12.80%-20% | Nugaraitė *et al.*, 2018 | |
| | *Molineus patens* | NO | ES | 68% | Martínez-Rondán *et al.*, 2017 | |
| | *Pseudamphistomum truncatum* | YES | LT | 17.90%-30% | Nugaraitė *et al.*, 2018 | |
| | *Sarcosystis lutrae* | NO | LT | 13.60% | Prakas *et al.,* 2018 | |
| | *SARS-CoV-2* | YES | NL | 19.40% | Oreshkova *et al.*, 2020 | Dead mink positive for viral RNA. Prevalence 100% of the throat swabs of dead animals. |
| | *Skrjabingylus nasicola* | NO | DE | 53.30% | Heddergott *et al.,* 2016 | |
| | *Staphylococcus aureus methicillin-resistant* (LA-MRSA) | YES | DK | 34%-40% | Hansen *et al.*, 2017 | |
| | *Strigea strigis* | | LT | 28.20%-30% | Nugaraitė *et al.*, 2018 | Metacercariae. |
| | *Taenia martis* | YES | LT | 2.50% | Nugaraitė *et al.*, 2018 | |
| | *Toxocara* spp. | YES | PL | 21.70% | Kołodziej-Sobocińska *et al.,* 2020 | |
| | *Toxoplasma gondii* | YES | PL | 25% | Sroka *et al.,* 2019 | |
| | *Toxoplasma gondii* | YES | ES | 78.80% | Ribas *et al.*, 2018 | |
| | *Toxoplasma gondii* | YES | ES | 0% | Criado-Fornelio *et al.,* 2018 | |
| | *Trichinella* spp. | YES | PL | 3.30% | Hurníková *et al.,* 2016 | |
| | *Troglotrema acutum* | | ES | 2% | Martínez-Rondán *et al.*, 2017 | |
| | Unidentified trematode | | ES | 2% | Martínez-Rondán *et al.*, 2017 | |

| Species | Pathogen | Zoonotic | Country | Prevalence | Reference | Notes |
|---|---|---|---|---|---|---|
| *Nyctereutes procyonoides* | *Aelurostrongylus abstrusus* | NO | DK | 0% | Lemming *et al.*, 2020 | |
| | *Alaria alata* | YES | AT | 30% | Duscher *et al.*, 2017 | |
| | *Alaria alata* | YES | EE | 13.30% | Laurimaa *et al.*, 2016 | Metacercariae. |
| | *Alaria alata* | YES | EE | 68.30% | Laurimaa *et al.*, 2016 | |
| | *Alaria alata* | YES | PL | 94.30% | Karamon *et al.*, 2016 | |
| | Anaplasmataceae | YES | AT | 0% | Duscher *et al.*, 2017 | |
| | *Angiostrongylus vasorum* | NO | EE | 1.30% | Laurimaa *et al.*, 2016 | |
| | *Angiostrongylus vasorum* | NO | DK | 3.20% | Lemming *et al.*, 2020 | |
| | *Aonchotheca putorii* | YES | EE | 3.60% | Laurimaa *et al.*, 2016 | |
| | *Apophallus* spp. | YES | PL | 15.10% | Karamon *et al.*, 2016 | |
| | *Babesia cf microti* | YES | AT | 62.50% | Duscher *et al.*, 2017 | |
| | *Borrelia* spp. | YES | PL | 25% | Wodecka *et al.*, 2016 | |
| | *Candidatus Neoerlichia* sp. | YES | PL | 30% | Hildebrand *et al.*, 2018 | |
| | *Capillaria aerophila (Eucoleus aerophilus)* | YES | DK | 1.90% | Lemming *et al.*, 2020 | |
| | *Capillaria plica (Pearsonema plica)* | NO | DK | 0.50% | Petersen *et al.*, 2018b | |
| | *Chlamydia* spp. | YES | UA | 0% | Ksyonz *et al.*, 2019 | |
| | *Crenosoma vulpis* | NO | EE | 15% | Laurimaa *et al.*, 2016 | |
| | *Crenosoma vulpis* | NO | DK | 5.30% | Lemming *et al.*, 2020 | |
| | *Dipylidium caninum* | NO | AT | 20% | Duscher *et al.*, 2017 | |
| | *Echinococcus multilocularis* | YES | AT | 10% | Duscher *et al.*, 2017 | |
| | *Echinococcus multilocularis* | YES | DK | 0.70% | Petersen *et al.*, 2018a | |
| | *Echinococcus multilocularis* | YES | EE | 1.60% | Laurimaa *et al.*, 2016 | |

| Species | Pathogen | Zoonotic | Country | Prevalence | Reference | Notes |
|---|---|---|---|---|---|---|
| *Nyctereutes procyonoides* | *Echinococcus multilocularis* | YES | NL | 11.10% | Maas *et al.,* 2016 | PCR. |
| | *Echinococcus multilocularis* | YES | DK | 0% | Oksanen *et al.*, 2016 | Pooled prevalence from Enemark, 2013; Al-Sabi *et al.*, 2013; EFSA, 2015. |
| | *Echinococcus multilocularis* | YES | DE | 2.50% | Oksanen *et al.*, 2016 | Pooled prevalence from Thiess *et al.*, 2001; Thiess, 2004; Schwarz *et al.*, 2011. |
| | *Echinococcus multilocularis* | YES | NL | 0% | EFSA, 2015 | |
| | *Echinococcus multilocularis* | YES | FI | 0% | Oksanen *et al.*, 2016 | Pooled prevalence from EFSA, 2013, 2014, 2015. |
| | *Echinococcus multilocularis* | YES | PL | 10.40% | Oksanen *et al.*, 2016 | Pooled prevalence from Machnicka-Rowińska *et al.*, 2002; Machnicka *et al.*, 2003; EFSA, 2015. |
| | *Echinococcus multilocularis* | YES | PL | 0% | Karamon *et al.*, 2016 | |
| | *Echinococcus multilocularis* | YES | SE | 0% | Wahlström *et al.*, 2011 | |
| | *Echinococcus multilocularis* | YES | SK | 28% | Oksanen *et al.*, 2016 | Pooled prevalence from Letková *et al.*, 2008; Hurníková *et al.*, 2009; EFSA, 2015. |
| | *Echinococcus multilocularis* | YES | LV | 8.10% | Bagrade *et al.,* 2016 | |
| | *Echinococcus multilocularis* | YES | LV | 21% | Bagrade *et al.*, 2008 | |
| | *Echinococcus multilocularis* | YES | LT | 8.20% | Bružinskaitė-Schmidhalter *et al.*, 2012 | |
| | *Echinococcus multilocularis* | YES | UA | 0% | Kornyushin *et al.*, 2011 | |
| | Echinostomatidae | YES | PL | 18.90% | Karamon *et al.,* 2016 | |
| | *Eucoleus aerophilus* | YES | EE | 30% | Laurimaa *et al.*, 2016 | |
| | *Francisella tularensis* | YES | DE | 16.70% | Schulze *et al.,* 2016 | |
| | *Hepatitis E Virus (HEV)* | YES | DE | 34.30% | Dahnert *et al.,* 2018 | |
| | Hookworms | YES | PL | 83% | Karamon *et al.,* 2016 | |
| | *Isthmiophora melis* | NO | AT | 20% | Duscher *et al.*, 2017 | |

| Species | Pathogen | Zoonotic | Country | Prevalence | Reference | Notes |
|---|---|---|---|---|---|---|
| *Nyctereutes procyonoides* | *Isthmiophora melis* | NO | EE | 6% | Laurimaa *et al.*, 2016 | |
| | *Ixodes ricinus* | | PL | | Wodecka *et al.,* 2016 | Raccoon dogs harbor seven-fold more ticks than badgers. |
| | *Mesocestoides* spp. | YES | AT | 40% | Duscher *et al.*, 2017 | |
| | *Mesocestoides* spp. | YES | EE | 21.30% | Laurimaa *et al.*, 2016 | *M. lineatus, M. litteratus* |
| | *Metorchis bilis* | YES | EE | 19.50% | Laurimaa *et al.*, 2016 | |
| | *Molineus patens* | NO | EE | 13.70% | Laurimaa *et al.*, 2016 | |
| | *Molineus* spp. | NO | AT | 30% | Duscher *et al.*, 2017 | |
| | *Molineus* spp. | NO | PL | 41.50% | Karamon *et al.,* 2016 | |
| | *Pearsonema plica* | NO | EE | 10.80% | Laurimaa *et al.*, 2016 | |
| | *Plagiorchis elegans* | NO | EE | 0.80% | Laurimaa *et al.*, 2016 | |
| | *Pygidiopsis summa* | YES | DK | 3% | Al-Sabi *et al.,* 2013 | |
| | *Taenia policantha* | NO | EE | 8.40% | Laurimaa *et al.*, 2016 | |
| | *Taenia* spp. | YES | AT | 20% | Duscher *et al.*, 2017 | |
| | *Tick-Borne Encephalitis Virus* (TBEV) | YES | FI | | Uusitalo *et al.*, 2020 | Has a role in the cycle. |
| | *Toxocara canis* | YES | AT | 20% | Duscher *et al.*, 2017 | |
| | *Toxocara leonina* | YES | AT | 10% | Duscher *et al.*, 2017 | |
| | *Toxocara* spp. | YES | EE | 8% | Laurimaa *et al.*, 2016 | *T. canis, T. leonina* |
| | *Toxocara* spp. | YES | PL | 15.10% | Karamon *et al.,* 2016 | |
| | *Toxoplasma gondii* | YES | PL | 7.70% | Sroka *et al.,* 2019 | |
| | *Trichinella* spp. | YES | AT | 0% | Duscher *et al.*, 2017 | |
| | *Trichinella* spp. | YES | EE | 57.50% | Kärssin *et al.*, 2017 | |
| | *Trichinella* spp. | YES | NL | 11.10% | Maas *et al.,* 2016 | |
| | *Trichinella* spp. | YES | PL | 39.80% | Cybulska *et al.,* 2019 | |
| | *Uncinaria stenocephala* | YES | AT | 40% | Duscher *et al.*, 2017 | |
| | *Uncinaria stenocephala* | YES | EE | 97.60% | Laurimaa *et al.*, 2016 | |
| | Unidentified lungworm | | DK | 0.80% | Lemming *et al.*, 2020 | |

| Species | Pathogen | Zoonotic | Country | Prevalence | Reference | Notes |
|---|---|---|---|---|---|---|
| *Ondatra zibethicus* | *Bartonella* spp. | YES | BE | | Krügel *et al.,* 2020 | Detection in a by-caught specimen. |
| | *Chlamydia* spp. | YES | UA | 33.30% | Ksyonz *et al.,* 2019 | |
| | *Cryptosporidium* spp. | YES | DE | | Petri *et al.,* 1997 | Muskrat can contaminate waters. |
| | *Echinococcus multilocularis* | YES | BE | 11.20% | Hanosset *et al.,* 2008 | |
| | *Echinococcus multilocularis* | YES | NL | 0.10% | Borgsteede *et al.,* 2003 | |
| | *Francisella tularensis* | YES | DE | 0% | Schulze *et al.,* 2016 | |
| | *Giardia duodenalis (Giardia lamblia)* | YES | RO | 100% | Adriana *et al.,* 2016 | One sample analysed. |
| | *Leptospira spp.* | YES | DE | 5.90% | Hurd *et al.,* 2017 | |
| | *Toxoplasma gondii* | YES | PL | 6.30% | Sroka *et al.,* 2019 | |
| | *Yersinia pestis* | YES | | | Anon., 1940 | Appears to be susceptible to plague. |
| *Procyon lotor* | *Acanthocephala* | YES | PL | 1.90% | Karamon *et al.,* 2014 | |
| | *Alaria alata* | YES | AT | 0% | Duscher *et al.*, 2017 | |
| | *Alaria alata* | YES | DE | 33.30% | Rentería-Solís *et al.,* 2013 | |
| | *Anaplasma phagocytophilum* | YES | PL | 0.80% | Hildebrand *et al.,* 2018 | |
| | Anaplasmataceae | YES | AT | 0% | Duscher *et al.,* 2017 | |
| | *Ancylostoma* spp. | YES | PL | 4.40% | Popiołek *et al.,* 2011 | |
| | *Babesia cf microti* | YES | AT | 0% | Duscher *et al.*, 2017 | |
| | *Baylisascaris procyonis* | YES | DE | 43.60% | Heddergott *et al.,* 2020b | |
| | *Baylisascaris procyonis* | YES | DE | 76.20% | Rentería-Solís *et al.*, 2018 | |
| | *Baylisascaris procyonis* | YES | DE | 39% | Winter, 2005 | |
| | *Baylisascaris procyonis* | YES | DE | 71.40% | Gey, 1998 | |
| | *Baylisascaris procyonis* | YES | DE | 80% | Hohmann *et al.*, 2002 | |
| | *Baylisascaris procyonis* | YES | DK | 11% | Al-Sabi *et al.*, 2016 | |
| | *Baylisascaris procyonis* | YES | PL | 3.30% | Popiołek *et al.*, 2011 | |
| | *Baylisascaris procyonis* | YES | PL | 1.90% | Karamon *et al.,* 2014 | |

| Species | Pathogen | Zoonotic | Country | Prevalence | Reference | Notes |
|---|---|---|---|---|---|---|
| *Procyon lotor* | *Baylisascaris procyonis* | YES | PL | 3.70% | Bartoszewicz *et al.*, 2008 | |
| | *Canine Adenovirus 1* (CAdV-1) | NO | DE | 0% | Schulze *et al.*, 2019 | |
| | *Canine Adenovirus 1* (CAdV-1) | NO | DE | 0% | Hechinger *et al.,* 2017 | |
| | *Canine Adenovirus 1* (CAdV-2) | NO | DE | 0% | Schulze *et al.*, 2019 | |
| | *Canine Distemper Virus* (CDV) | NO | DE | 10.80% | Wibbelt *et al.*, 2008 | |
| | *Canine Distemper Virus* (CDV) | NO | DE | 46% | Hechinger *et al.,* 2017 | |
| | *Canine Distemper Virus* (CDV) | NO | DE | 76.30% | Rentería-Solís *et al.,* 2014b | |
| | *Capillaria* spp. | YES | PL | 25.50% | Karamon *et al.,* 2014 | |
| | Capillaridae | YES | PL | 33.30% | Popiołek *et al.,* 2011 | |
| | *Cryptosporidium* spp. | YES | LU | 12.40% | Heddergott *et al.,* 2020a | |
| | *Cryptosporidium* spp. | YES | PL | 34.70% | Leśniańska *et al.,* 2016 | |
| | *Cryptosporidium* spp. | YES | DE | 34.70% | Leśniańska *et al.,* 2016 | |
| | *Dipylidium caninum* | NO | AT | 0% | Duscher *et al.*, 2017 | |
| | *Echinococcus multilocularis* | YES | AT | 0% | Duscher *et al.*, 2017 | |
| | *Echinostoma* sp. | YES | PL | 2.20% | Popiołek *et al.,* 2011 | |
| | Echinostomatidae | YES | PL | 34.50% | Karamon *et al.,* 2014 | |
| | *Ehrlichia canis* | YES | ES | 2.60% | Criado-Fornelio *et al.,* 2018 | |
| | *Eimeria* spp. | YES | DE | 1.50% | Gey, 1998 | |
| | *Eimeria* spp. | YES | DE | 1.80% | Winter, 2005 | |
| | *Enterocytozoon bieneusi* | YES | PL | 4.10% | Leśniańska *et al.,* 2016 | |
| | *Francisella tularensis* | YES | DE | 0% | Schulze *et al.,* 2016 | |
| | *Hepatitis E Virus (HEV)* | YES | DE | 53.80% | Dahnert *et al.*, 2018 | |
| | *Hepatozoon canis* | NO | ES | 2.60% | Criado-Fornelio *et al.,* 2018 | |

| Species | Pathogen | Zoonotic | Country | Prevalence | Reference | Notes |
|---|---|---|---|---|---|---|
| *Procyon lotor* | *Isthmiophora melis* | NO | AT | 0% | Duscher *et al.*, 2017 | |
| | *Leishmania infantum* | YES | ES | 0% | Risueño *et al.,* 2018 | |
| | *Listeria* spp. | YES | PL | 7.10% | Nowakiewicz *et al.,* 2016 | |
| | *Lyssavirus rabies* | YES | Europe | | The Rabies Information System of the WHO Collaboration Centre for Rabies Surveillance and Research | 142 cases reported in Europe. http://rbe.fli.bund.de/ Default.aspx |
| | *Mesocestoides* spp. | YES | AT | 0% | Duscher *et al.*, 2017 | |
| | *Mesocestoides* spp. | YES | PL | 67.30% | Karamon *et al.,* 2014 | |
| | *Molineus* spp. | NO | AT | 13% | Duscher *et al.*, 2017 | |
| | *Neospora caninum* | NO | CZ | 17.60% | Kornacka *et al.*, 2018 | Antibodies. Prevalence 0% by PCR. |
| | *Neospora caninum* | NO | DE | 16.70% | Kornacka *et al.*, 2018 | Antibodies. Prevalence 0% by PCR. |
| | *Neospora caninum* | NO | PL | 13.30% | Kornacka *et al.*, 2018 | Antibodies. Prevalence 0% by PCR. |
| | *Placoconus lotoris* | NO | PL | 4.40% | Popiołek *et al.,* 2011 | |
| | *Salmonella* spp. | YES | PL | 5.70% | Nowakiewicz *et al.,* 2016 | |
| | *Sarcocystis* spp. | YES | DE | 4.40% | Stolte *et al.*, 1996 | |
| | *Sarcocystis* spp. | | DE | | Rentería-Solís *et al.*, 2014 | Cross-transmission of *S. scabiei* mites has been recorded. |
| | *Spirocerca lupi* | NO | PL | 8.80% | Popiołek *et al.,* 2011 | |
| | *Staphylococcus coagulase-positive* | YES | PL | 35.70% | Nowakiewicz *et al.,* 2016 | |
| | *Strongyloides procyonis* | YES | PL | 14.80% | Bartoszewicz *et al.*, 2008 | |
| | *Strongyloides procyonis* | YES | PL | 11% | Popiołek *et al.,* 2011 | |
| | *Taenia* spp. | YES | AT | 0% | Duscher *et al.*, 2017 | |
| | *Toxocara canis* | YES | AT | 0% | Duscher *et al.*, 2017 | |
| | *Toxocara leonina* | YES | AT | 0% | Duscher *et al.*, 2017 | |
| | *Toxoplasma gondii* | YES | CZ | 0% | Kornacka *et al.*, 2018 | Antibodies. Prevalence 47.1% by PCR. |
| | *Toxoplasma gondii* | YES | DE | 26% | Gey, 1998 | |
| | *Toxoplasma gondii* | YES | DE | 33.30% | Kornacka *et al.*, 2018 | Antibodies. Prevalence 33.3% by PCR. |
| | *Toxoplasma gondii* | YES | DE | 38.30% | Heddergot *et al.,* 2017 | Antibodies. |
| | *Toxoplasma gondii* | YES | LU | 19% | Heddergot *et al.,* 2017 | Antibodies. |

| Species | Pathogen | Zoonotic | Country | Prevalence | Reference | Notes |
|---|---|---|---|---|---|---|
| *Procyon lotor* | *Toxoplasma gondii* | YES | PL | 13.10% | Sroka *et al.,* 2019 | |
| | *Toxoplasma gondii* | YES | PL | 13.30% | Kornacka *et al.*, 2018 | Antibodies. Prevalence 40% by PCR. |
| | *Toxoplasma gondii* | YES | ES | 3.60% | Criado-Fornelio *et al.,* 2018 | |
| | *Trichinella* spp. | YES | AT | 0% | Duscher *et al.*, 2017 | |
| | *Trichinella* spp. | YES | CZ | 9.10% | Cybulska *et al.,* 2018 | |
| | *Trichinella* spp. | YES | DE | 0% | Cybulska *et al.,* 2018 | |
| | *Trichinella* spp. | YES | PL | 6% | Cybulska *et al.,* 2018 | |
| | *Uncinaria stenocephala* | YES | AT | 0% | Duscher *et al.*, 2017 | |
| | *Yersinia* spp. | YES | PL | 4.30% | Nowakiewicz *et al.,* 2016 | |
| *Sciurus carolinensis* | *Adenoviridae* | YES | IT | 0.90% | Romeo *et al.*, 2014b | |
| | *Aonchotheca annulosa* | NO | IT | 1.50% | Romeo *et al.*, 2014a | |
| | *Borrelia burgdoferi sensu lato* | YES | UK | 11.90% | Millins *et al.*, 2015 | |
| | *Cryptosporidium* spp. | YES | IT | 3.70% | Prediger *et al.,* 2017 | |
| | *Eimeria* spp. | YES | IT | 95.70% | Hofmannová *et al.*, 2016 | Plus the successful introduction of *E. lancasterensis.* |
| | Hymenolepididae | YES | IT | 0.40% | Romeo *et al.*, 2014a | |
| | *Hymenolepis* spp. | YES | IT | 0% | d'Ovidio *et al.*, 2015 | |
| | *Ljungan Virus* (LV) | YES | IT | 0% | Romeo *et al.*, 2014b | |
| | *Mycobacterium leprae* | YES | IT | 0% | Schilling *et al.*, 2019 | |
| | *Mycobacterium leprae* | YES | UK | 0% | Schilling *et al.*, 2019 | |
| | Oxyurida | YES | IT | 0.90% | Romeo *et al.*, 2014a | |
| | *Squirrelpox poxvirus* (SQPV) | NO | IE | 29% | Stritch *et al.*, 2015 | |
| | *Squirrelpox poxvirus* (SQPV) | NO | IE | 25% | Collins *et al.*, 2014 | Antibodies. Prevalence 10% by PCR. |
| | Strongylida | YES | IT | 4.40% | Romeo *et al.*, 2014a | |
| | *Strongyloides robustus* | NO | IT | 56.50% | Romeo *et al.*, 2014a | |
| | *Tick-Borne Encephalitis Virus* (TBEV) | YES | IT | 1.90%-2.50% | Romeo *et al.*, 2018 | |

| Species | Pathogen | Zoonotic | Country | Prevalence | Reference | Notes |
|---|---|---|---|---|---|---|
| *Sciurus carolinensis* | *Trichostrongylus calcaratus* | NO | IT | 6.50% | Romeo *et al.*, 2014a | |
| | *Trichostrongylus retortaeformis* | NO | IT | 0.80% | Romeo *et al.*, 2014a | |
| | *Trichuris muris* | NO | IT | 4.20% | Romeo *et al.*, 2014a | |
| | *Trypanoxyuris sciuri* | NO | IT | 2.30% | Romeo *et al.*, 2014a | |
| | *Usutu Virus* (USUV) | YES | IT | 3.20%-3.80% | Romeo *et al.*, 2018 | |
| | *West Nile Virus* (WNV) | YES | IT | 0.60% | Romeo *et al.*, 2018 | |

**List of the additional references for the pathogen studies (studies not directly included in the review)**

*Cervus nippon*

Dykova I, Blazek K (1972). Subcutaneous filariasis in red deer. *Acta Veterinaria* 41: 117-124.

Gibbs EPJ, Herniman KAJ, Lawman MJP (1975) Studies with foot-and-mouth disease virus in British deer (muntjac and sika): Clinical disease, recovery of virus and serological response. *Journal of Comparative Pathology* 85(3): 361-366

Mihalca AD, Păstrav IR, Sándor AD, Deak G, Gherman CM, Sarmaşi A, Votýpka J (2019) First report of the dog louse fly *Hippobosca longipennis* in Romania. *Medical and Veterinary Entomology* 33(4): 530-535.

Nechybová S, Vejl P, Hart V, Melounová M, Čílová D, Vašek J, Jankovská I, Vadlejch J, Langrova I (2018) Long-term occurrence of Trichuris species in wild ruminants in the Czech Republic. Parasitology Research 117(6): 1699-1708.

Rudaitytė-Lukošienė E, Prakas P, Butkauskas, D, Kutkienė L, Vepštaitė-Monstavičė I, Servienė E (2018) Morphological and molecular identification of *Sarcocystis* spp. from the sika deer (*Cervus nippon*), including two new species *Sarcocystis frondea* and *Sarcocystis nipponi*. *Parasitology Research* 117(5): 1305-1315.

*Myocastor coypus*

Michel V, Ruveon-Clouet N, Menard A, Sonrier C, Fillonneau C, Rakotovao F, Ganière JP, André-Fontaine G (2001) Role of the coypu (*Myocastor coypus*) in the epidemiology of leptospirosis in domestic animals and humans in France. *European Journal of Epidemiology* 17: 111-121.

*Neovison vison*

Hansen JE, Larsen AR, Skov RL, Chriél M, Larsen G, Angen Ø, Larsen J, Lassen DCK, Pedersen K (2017). Livestock-associated methicillin-resistant *Staphylococcus aureus* is widespread in farmed mink (*Neovison vison*). *Veterinary Microbiology* 207: 44-49.

Mañas S, Carlos Ceña J, Ruiz-Olmo J, Palazón S, Domingo M, Wolfinbarger JB, Bloom ME (2001) Aleutian mink disease parvovirus in wild riparian carnivores in Spain. *Journal of Wildlife Diseases* 37(1):138-144.

Martínez-Rondán F, Ruiz de Ybañez R, Tizzani P, López-Beceiro A, Fidalgo L, Martínez-Carrasco Pleite C (2017) The American mink (*Neovison vison*) is a competent host for native European parasites. *Veterinary Parasitology* 247: 93-99.

*Nyctereutes procyonoides*

Al-Sabi MNS, Chriél M,Hammer Jensen T, Larsen Enemark H (2013). Endoparasites of the raccoon dog (*Nyctereutes procyonoides*) and the red fox (*Vulpes vulpes*) in Denmark 2009–2012 – A comparative study. *International Journal for Parasitology: Parasites and Wildlife* 2: 144-151.

Bagrade G, Snabel V, Romig T, Ozolins J, Huettner M, Miterpáková M, et al. (2008) *Echinococcus multilocularis* is a frequent parasite of red foxes (*Vulpes vulpes*) in Latvia. *Helminthologia* 45: 157-161.

Bružinskaitė-Schmidhalter R, Šarkūnas M, Malakauskas A, Mathis A, Torgerson PR, Deplazes P. (2012) Helminths of red foxes (*Vulpes vulpes*) and raccoon dogs (*Nyctereutes procyonoides*) in Lithuania. *Parasitology* 139: 120-127.

Dähnert L, Conraths F, Reimer N, Groschup M, Eiden M (2018) Molecular and serological surveillance of Hepatitis E virus in wild and domestic carnivores in Brandenburg, Germany. *Transboundary and Emerging Diseases* 65(5).

EFSA (2015) Scientific opinion – Update on oral vaccination of foxes and raccoon dogs against rabies. *EFSA Journal* 13: 70.

Kornyushin VV, Malyshko EI, Malega AM (2011) The Helminths of wild predatory mammals of Ukraine. Cestodes. *Vestnik Zoologii* 45: 4-11.

Wahlström H, Lindberg A, Lindh J, Wallensten A, Lindqvist R, Plym-Forshell L et al. (2012) Investigations and actions taken during 2011 due to the first finding of *Echinococcus multilocularis* in Sweden. *Eurosurveillance* 17: 1-7.

*Ondatra zibethicus*

Anon, 1940. The possible role of the muskrat (*Ondatra zibethica* L.) in the epidemiology of plague. *Vestnik Mikrobiologii, Epidemiologii i Parazitologii*, 19(2).

Borgsteede FHM, van der Tibben JH, Giessen JWB (2003). The muskrat (*Ondatra zibethicus*) as intermediate host of cestodes in the Netherlands. *Veterinary Parasitology* 117: 29-36.

Hanosset R, Saegerman C, Adant S, Massart L, Losson B (2008) *Echinococcus multilocularis* in Belgium: prevalence in red foxes (*Vulpes vulpes*) and in different species of potential intermediate hosts. *Veterinary Parasitology* 151(2/4): 212-217.

Petri C, Karanis P, Renoth S (1997) Cryptosporidium infections in muskrat (*Ondatra zibethica*). *Parasite* 4(4): 369-371.

*Procyon lotor*

Bartoszewicz M, Okarma H, Zalewski A, Szczesna J (2008). Ecology of the raccoon (*Procyon lotor*) from western Poland. *Annales Zoologici Fennici* 45(4): 291-298.

Dähnert L, Conraths FJ, Reimer N, Groschup MH, Eiden M (2018) Molecular and serological surveillance of hepatitis E virus in wild and domestic carnivores in Brandenburg, Germany. *Transboundary Emerging Diseases* 65: 1377–1380.

Gey AB (1998). Endoparasite fauna of the raccoon (*Procyon lotor*) in Hesse, Germany. PhD thesis, Justus-Liebieg University Giesen, Germany. [in German with English summary].

Hohmann U, Voig S, Andreas U (2002) Raccoons take the offensive. A current assessment. In: *Biologische Invasionen*, edited by Kowarik I, Starfinger U, *Neobiota1*, 191-192.

Stolte M, Odening K, Walter G, Bockhardt I (1996) The raccoon as intermediate host of three Sarcocystis species in Europe. *Comparative parasitology* 63(1): 145-149.

Wibbelt G, Speck S, Fickel J, Köhnemann B, Michler F.-U. (2008) Outbreak of canine distemper in raccoons (*Procyon lotor*) in Germany. 8th Conference of the Wildlife Disease Association, Rovinj/Kroatien, pp 22.

Winter M, Stubbe M, Heidecke D (2005) Zur Ökologie des Waschbären (*Procyon lotor* L., 1758) in Sachsen-Anhalt. *Beitr Jagd Wildforsch* 30: 303-322. [in German].

1262
